# Hybridization promotes asexual reproduction in *Caenorhabditis* nematodes

**DOI:** 10.1101/588152

**Authors:** Piero Lamelza, Janet M. Young, Luke M. Noble, Lews Caro, Arielle Isakharov, Meenakshi Palanisamy, Matthew V. Rockman, Harmit S. Malik, Michael Ailion

**Author notes:** Institut de Biologie, École Normale Supérieure, Centre National de la Recherche Scientifique (CNRS) UMR 8197, Institut National de la Santé et de la Recherche Médicale (INSERM) U1024, Paris, France, F-75005.

## Abstract

Although most unicellular organisms reproduce asexually, most multicellular eukaryotes are obligately sexual. This implies that there are strong barriers that prevent the origin or maintenance of asexuality arising from an obligately sexual ancestor. By studying rare asexual animal species we can gain a better understanding of the circumstances that facilitate their evolution from a sexual ancestor. Of the known asexual animal species, many originated by hybridization between two ancestral sexual species. The balance hypothesis predicts that genetic incompatibilities between the divergent genomes in hybrids can modify meiosis and facilitate asexual reproduction, but there are few instances where this has been shown. Here we report that hybridizing two sexual *Caenorhabditis* nematode species (*C. nouraguensis* females and *C. becei* males) alters the normal inheritance of the maternal and paternal genomes during the formation of hybrid zygotes. Most offspring of this interspecies cross die during embryogenesis, exhibiting inheritance of a diploid *C. nouraguensis* maternal genome and incomplete inheritance of *C. becei* paternal DNA. However, a small fraction of offspring develop into viable adults that can be either fertile or sterile. Fertile offspring are produced asexually by sperm-dependent parthenogenesis (also called gynogenesis or pseudogamy); these progeny inherit a diploid maternal genome but fail to inherit a paternal genome. Sterile offspring are hybrids that inherit both a diploid maternal genome and a haploid paternal genome. Whole-genome sequencing of individual viable worms shows that diploid maternal inheritance in both fertile and sterile offspring results from an altered meiosis in *C. nouraguensis* oocytes and the inheritance of two randomly selected homologous chromatids. We hypothesize that hybrid incompatibility between *C. nouraguensis* and *C. becei* modifies maternal and paternal genome inheritance and indirectly induces gynogenetic reproduction. This system can be used to dissect the molecular mechanisms by which hybrid incompatibilities can facilitate the emergence of asexual reproduction.

**AUTHOR SUMMARY:** Eukaryotes employ two major reproductive strategies: sexual and asexual reproduction. Both types of reproduction have distinct theoretical costs and benefits, and most unicellular eukaryotes can switch between both modes. However, most multicellular eukaryotes are obligately sexual, implying that there are barriers to the evolution of asexuality from a sexual ancestor. Of the few asexual animal species, many are hybrids of two ancestral sexual species, suggesting that novel genetic interactions in hybrids facilitate the evolution of asexuality. One model suggests that genetic incompatibilities between divergent genomes in hybrids can modify female meiosis and paternal genome inheritance to facilitate asexual reproduction. While studying interspecies hybridizations of *Caenorhabditis* nematodes, we found that crossing two sexual species (*C. nouraguensis* and *C. becei*) disrupts female meiosis and paternal genome inheritance. Most offspring die during embryogenesis, but on rare occasions develop into viable and fertile adults that are produced asexually. This asexual reproduction involves the unusual production of eggs carrying two sets of maternal chromosomes and the loss of the paternal set of chromosomes. We hypothesize that genetic incompatibility between these two species disrupts maternal and paternal genome inheritance. This interspecies hybridization may serve as a model to study how genetic incompatibilities facilitate the emergence of asexuality.

## INTRODUCTION

Sex is the most common form of reproduction in multicellular eukaryotes. In sex, diploid females and males generate haploid eggs and sperm that fuse to produce diploid offspring. Sex has theoretical benefits that may explain its prevalence. Namely, recombination during meiosis generates genotypic diversity that can be used to adapt to changing environments or purge deleterious alleles [1].

Despite the predicted advantages of sex, it also has theoretical costs, such as the need to find a mate and the production of offspring that inherit only half of an individual’s genome [2, 3]. By contrast, females of asexual species independently generate their own offspring and pass on their entire genome. Consistent with sex being costly, most unicellular organisms reproduce by either obligate or facultative asexuality [4, 5], which involves being able to reproduce either sexually or asexually as conditions dictate. However, because obligate sex is the predominant form of reproduction in multicellular organisms [6], it is likely there are strong barriers that prevent the origin of asexuality from a sexual ancestor or its persistence [7]. By studying rare cases of asexual animal species, we can better understand the circumstances that facilitate the evolution of asexual reproduction.

Here we focus on two key aspects of how asexuality evolves from sexually reproducing organisms: diploid maternal inheritance and paternal genome loss. In contrast to sexual females that produce haploid eggs, asexual females produce diploid eggs that either develop independently of sperm fertilization, known as parthenogenesis [8], or require fertilization but fail to inherit the paternal genome, known as sperm-dependent parthenogenesis, gynogenesis, or pseudogamy [9]. Thus, understanding how egg production can be modified to result in diploid maternal inheritance and how the paternal genome is lost in gynogenetic reproduction should provide insight into how asexuality can arise.

Interestingly, many asexual animal species are hybrids of two ancestral sexually reproducing species, suggesting that hybridization may play a role in the generation of asexual species. However, hybridization is typically accompanied by deleterious incompatibilities between the two divergent genomes. For example, hybrid incompatibilities are known to disrupt gametogenesis and cause sterility [10, 11], or perturb paternal genome replication and inheritance and cause aneuploidy and lethality [12–14]. One model of how hybridization might lead to asexuality is known as the “balance hypothesis.” This model posits that in asexual hybrid species, interspecies incompatibilities disrupt female meiosis enough to produce diploid eggs, but do not significantly reduce the fertility or viability of hybrids [15, 16]. Other interspecies incompatibilities, such as incompatibility between components of the maternal egg cytoplasm and the paternal DNA [12, 14], may also facilitate paternal genome loss during gynogenetic reproduction. In support of this possibility, the females of many gynogenetic species hybridize with males of a different species [9, 17]. Thus, incompatibilities between two species that alter meiosis and the fidelity of paternal genome inheritance may actually in combination facilitate the generation of asexually-produced viable progeny, even though such incompatibilities on their own would be deleterious and lead to sterility or lethality.

In support of the balance hypothesis, there is good evidence that genetic incompatibilities facilitate diploid maternal inheritance in the hybrid parthenogenetic lizard *Aspidoscelis neomexicana* [18]. In most *A. neomexicana* female oocytes, the divergent homologous chromosomes fail to properly pair, resulting in arrest during meiotic prophase. However, instead of these hybrid females being sterile, a small number of oocytes duplicate their genomes and become tetraploid. Pairing and recombination in these tetraploid oocytes occur between genetically identical chromosomes, allowing these oocytes to progress through meiosis and generate diploid eggs that develop independently of fertilization.

It is unclear how often interspecies incompatibilities facilitate asexual reproduction, and the mechanisms that allow such hybrids to be viable and fertile remain largely unknown. We looked for examples of asexual reproduction in interspecies crosses within the genus of *Caenorhabditis* nematodes. Here we report that a cross between two sexual *Caenorhabditis* species (*C. nouraguensis* and *C. becei)* results in most offspring dying during embryogenesis, but also rare viable progeny that are asexually produced. We hypothesize that genetic incompatibility between these two *Caenorhabditis* species facilitates diploid maternal inheritance and paternal genome loss. Both processes usually result in aneuploidy that leads to sterility or death of hybrid embryos, but on rare occasions can result in production of viable offspring by gynogenetic reproduction. The hybridization between these two *Caenorhabditis* species might serve as an excellent cellular and genetic model to study how interspecies incompatibilities can modify normal sexual reproduction to facilitate the evolution of asexuality.

## RESULTS

### Reciprocal *C. nouraguensis* x *C. becei* crosses exhibit distinct F1 embryonic lethal phenotypes

*C. nouraguensis* and *C. becei* are obligately outcrossing sister species consisting of female and male individuals (Fig 1A). We initially studied the hybridization of *C. nouraguensis* strain JU1825 with *C. becei* strain QG711. The mating of individuals within the same strain results in a high proportion of viable progeny (91-99% viability, Fig S1A, Videos S1 and S2). However, mating *C. becei* females (QG711) to *C. nouraguensis* males (JU1825) results in 100% embryonic death with a stereotypical arrest at approximately the 32-cell stage (Fig 1B, Video S3). In the reciprocal cross between *C. nouraguensis* females (JU1825) and *C. becei* males (QG711), most F1 embryos also die (∼99.9%), but some embryos arrest early in development (approximately 1-4 cell stage) whereas some arrest at later stages (approximately 44-87 cell stage) (Video S4). From the cross of *C. nouraguensis* females (JU1825) and *C. becei* males (QG711), approximately 0.1% of F1 progeny survived and became viable adults (Fig 1B). Fecundity of *C. nouraguensis* females crossed to *C. becei* males was similar to *C. nouraguensis* females in intraspecific crosses on the first day of egg-laying but was mildly reduced on the second and third days of egg-laying (Fig S1B), consistent with possible deleterious effects of interspecific sperm [19].

**Fig 1.**
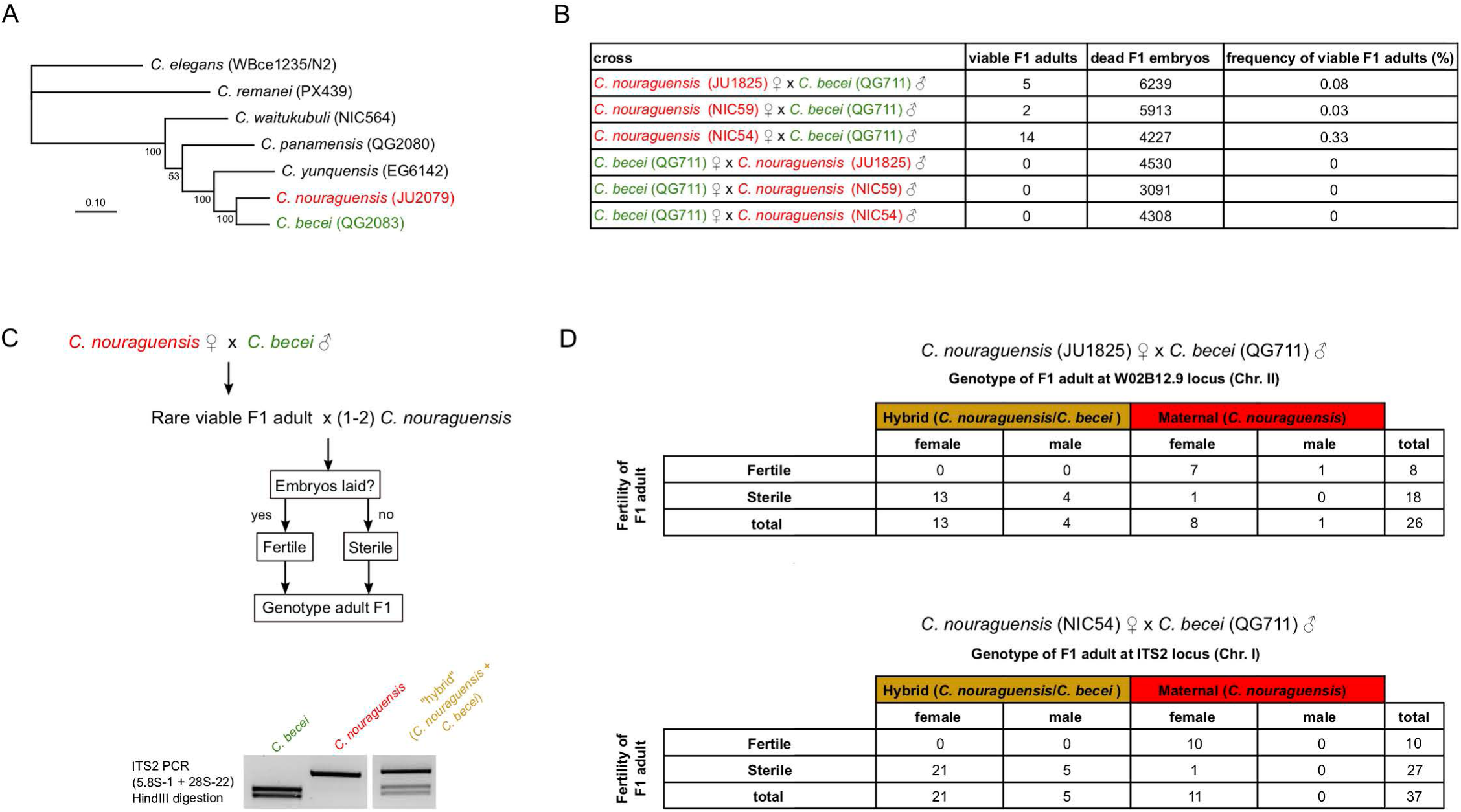
Crossing *C. nouraguensis* females to *C. becei* males results in sterile F1 with hybrid genotypes and fertile F1 with only-maternal genotypes. **(A)** A maximum likelihood phylogeny of several *Caenorhabditis* species closely related to *C. nouraguensis* and *C. becei*, with strain names in parentheses. *C. elegans and C. remanei* were used as outgroups. The scale bar represents 0.10 substitutions per site. Bootstrap support values are indicated to the left of each node (percent of 100 bootstrap replicates). See Stevens et al. (2019) for a more complete *Caenorhabditis* phylogeny [49]. **(B)** Frequency of viable F1 adults in crosses between *C. nouraguensis* and *C. becei*. NIC54 females yield a significantly higher proportion of viable F1 adults when crossed to *C. becei* (QG711) males than either JU1825 or NIC59 females (Chi-square with Yates correction, P=0.0065 NIC54 vs. JU1825, P=0.0005 NIC54 vs. NIC59). There was not a significant difference in the frequency of rare viable F1 adults produced by JU1825 and NIC59 females (Chi-square with Yates correction, P=0.49). The crosses between *C. becei* females and *C. nouraguensis* males serve as controls for accidental contamination of plates with embryos or larvae from either parental species -- no viable adults were found among more than 12,000 F1 screened for each cross (Fig S2). **(C)** Flowchart showing how fertility of rare viable F1 was tested (also see Materials and Methods). The gel shows how a PCR-RFLP assay distinguishes between *C. nouraguensis* and *C. becei* alleles at the ITS2 locus. **(D)** Tables showing the relationship between the fertility, genotype, and sex of rare viable F1 derived from crossing either *C. nouraguensis* (JU1825) females to *C. becei* (QG711) males (top table, genotyped at the W02B12.9 locus) or *C. nouraguensis* (NIC54) females to *C. becei* (QG711) males (bottom table, genotyped at the ITS2 locus). Both sterile and fertile F1 exhibited a more strongly female-biased sex ratio than that seen in intraspecies crosses (Fig S1A).

### Crossing *C. nouraguensis* females to *C. becei* males results in rare asexually-produced progeny

Interestingly, a small fraction of F1 progeny from *C. nouraguensis* female x *C. becei* male crosses develop into viable adults. Such viable progeny were found using three different *C. nouraguensis* strains and two *C. becei* strains (Fig S2), indicating that the production of viable progeny is a general feature of *C. nouraguensis* female x *C. becei* male crosses. By contrast, no surviving F1 were found in the reciprocal *C. becei* female x *C. nouraguensis* male crosses or crosses between *C. nouraguensis* females and males of other closely related *Caenorhabditis* species (Fig 1B and Fig S2). Thus, production of viable F1 progeny is specific to the *C. nouraguensis* female x *C. becei* male cross. We tested the fertility of each viable F1 animal and found that ∼2/3 of F1 were sterile and ∼1/3 were fertile (JU1825 x QG711: 18 sterile, 8 fertile; NIC54 x QG711: 27 sterile, 10 fertile) (Fig 1D).

We next tested if the rare viable F1 were indeed interspecies hybrids. We genotyped each F1 at one of two autosomal loci using PCR-restriction digest assays that distinguish between *C. nouraguensis* and *C. becei* alleles (Fig 1C). We found that about two-thirds of viable F1 inherited both parental alleles and therefore are genetic hybrids (JU1825 x QG711: 17/26 F1; NIC54 x QG711: 26/37 F1). However, to our surprise, we found that the remaining viable F1 adults carry only the maternal *C. nouraguensis* allele and no paternal *C. becei* allele (JU1825 x QG711: 9/26 F1; NIC54 x QG711: 11/37 F1) (Fig 1D). Interestingly, all fertile F1 had only the maternal *C. nouraguensis* allele and not the paternal *C. becei* allele, while all but two sterile F1 had a hybrid genotype with alleles from both parents. We observed a similar correlation between genotype and fertility among rare viable F1 animals when a different paternal *C. becei* strain was used (Fig S3).

Interspecies hybrids are often sterile [10], but it was surprising to find fertile F1 that inherited only maternal *C. nouraguensis* DNA. These fertile F1 behave like *C. nouraguensis* when backcrossed to either parental species. Backcrossing fertile F1 to *C. nouraguensis* results in many viable F2 progeny that develop to adulthood. By contrast, backcrossing fertile F1 to *C. becei* results in almost all F2 dying during embryogenesis other than the few rare viable F2 progeny in crosses to *C. becei* males (as in the original interspecies cross). Thus, it appears that interspecies hybridization between *C. nouraguensis* females and *C. becei* males produces a low frequency of fertile F1 that carry only maternal *C. nouraguensis* DNA and thus are produced asexually.

One way *C. nouraguensis* females might produce fertile F1 progeny with a maternal genotype would be to generate sperm and reproduce as hermaphrodites at a low frequency. To test this possibility, we monitored 60 virgin females from two *C. nouraguensis* strains (JU1825 and NIC59) and two *C. becei* strains (QG704 and QG711) for five days but failed to find any progeny. Thus, production of fertile F1 is not due to cryptic hermaphroditism but requires mating to *C. becei* males. We conclude that fertile F1 are the result of gynogenetic reproduction in which *C. nouraguensis* oocytes require fertilization by *C. becei* sperm to initiate development, but do not inherit the paternal *C. becei* genome.

### Asexually-produced F1 females are diploid

How do the fertile F1 compensate for the lack of a paternal haploid genome? We presumed that the autosomes of the fertile F1 had to be at least diploid for viability because haploid *C. elegans* individuals die during embryogenesis [20], and all well-studied *Caenorhabditis* species have five diploid autosomes and an X chromosome that is either diploid (females) or haploid (males) [21–23].

We determined the ploidy of fertile F1 females by counting the number of chromosomes in their oocytes. In *C. elegans*, the female germline produces oocytes containing chromosomes that have undergone prophase of meiosis I (DNA replication and recombination) but have yet to undergo the two meiotic divisions. The oocyte contains six bivalents, each of which is composed of a pair of homologous chromosomes and their sister chromatids held together by a combination of recombination and sister chromatid cohesion (Fig 2A). Therefore, diploid individuals have six bivalents in their oocytes, which appear as six distinct DAPI-staining bodies (Fig 2B). In *C. elegans*, meiotic recombination occurs only once per homologous set of chromosomes [24]. Thus, a triploid individual will have two homologous chromosomes that recombine and form a bivalent as well as a non-recombinant homolog as a univalent. Therefore, triploid individuals have six bivalents and six univalents in mature oocytes, which appear as twelve DAPI-staining bodies (Fig 2B) [25].

**Fig 2.**
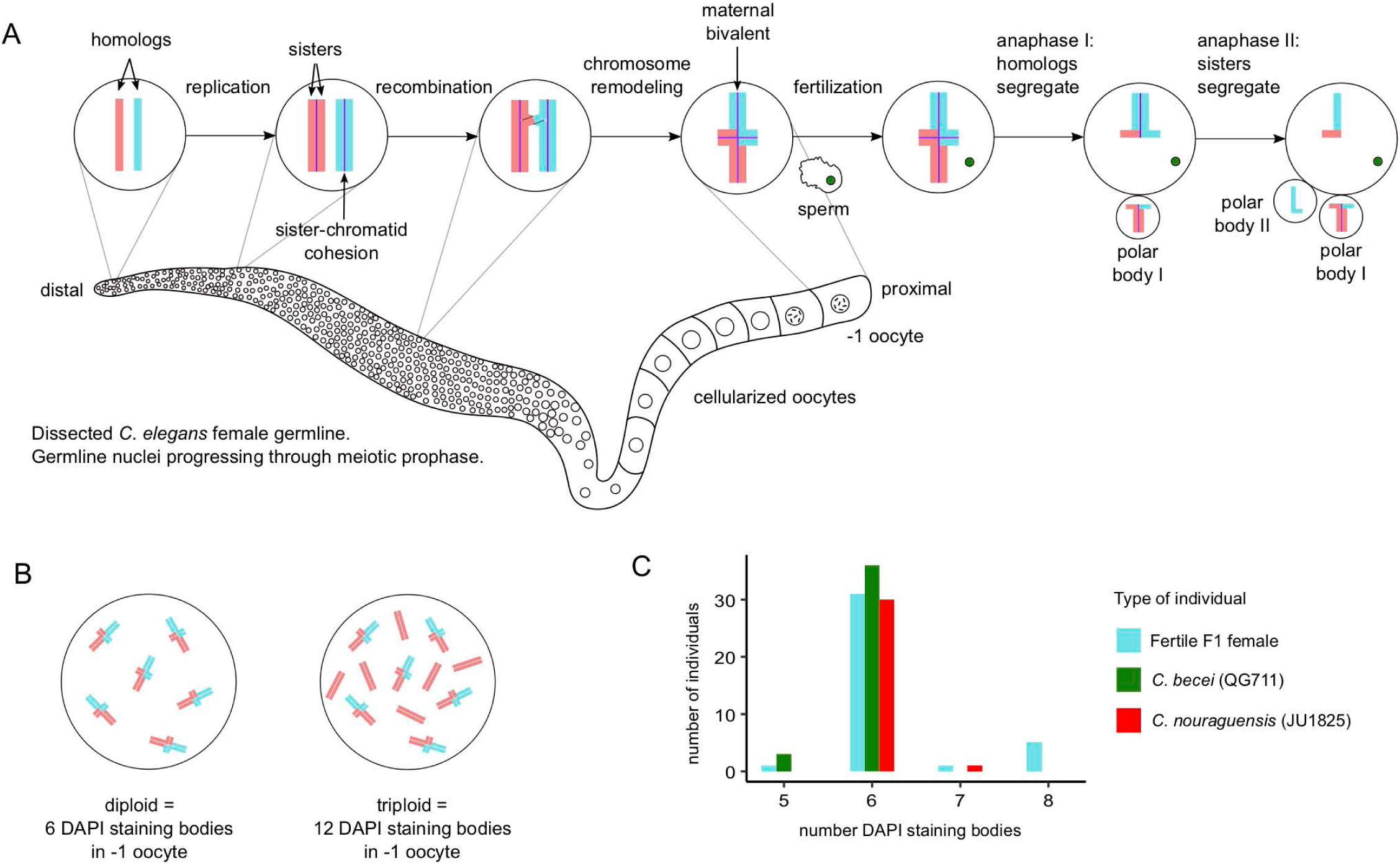
Asexually-produced F1 females are diploid. **(A)** Schematic of one of the two germlines of a *Caenorhabditis* female. The germline is a syncytial tube with nuclei (depicted as small circles) hugging its circumference. Germline nuclei are generated from mitotically dividing stem cells at the distal tip and migrate proximally while undergoing meiotic prophase. Homologs of each of the six chromosomes replicate their DNA, with the resulting sister chromatids held together by sister chromatid cohesion (purple lines). Homologous chromosomes undergo one crossover biased towards a chromosome end and are remodeled into a cruciform structure. Eventually, cell membranes form around the nuclei, resulting in mature oocytes. In mature oocytes, homologous chromosomes and their sister chromatids form bivalents that are held together by a combination of sister chromatid cohesion and recombination. The −1 oocyte is fertilized by sperm, triggering the reductional first meiotic division in which sister chromatid cohesion is lost between homologs, allowing them to segregate during anaphase I. One set of homologs is segregated into the first polar body. During the second meiotic division, sister chromatid cohesion is lost between sister chromatids, which then segregate during anaphase II. One chromatid is segregated into the second polar body and the other is inherited by the oocyte. **(B)** Because only one crossover occurs per homologous set of chromosomes, diploid individuals have six bivalents (6 DAPI-staining bodies) while triploids have six bivalents and six univalents (12 DAPI-staining bodies). **(C)** Most *C. nouraguensis* (JU1825), *C. becei* (QG711) and fertile F1 females have 6 DAPI-staining bodies, consistent with being diploid for six chromosomes. However, some fertile F1 females have a slightly higher number of DAPI-staining bodies, suggesting they are mostly diploid, but inherit some extra pieces of DNA. See Fig S4 for examples of DAPI staining.

We crossed *C. nouraguensis* (JU1825) females to *C. becei* (QG711) males, collected fertile F1 females and dissected and stained their germlines with DAPI. As controls, we examined the germlines of JU1825 and QG711 females after mating with conspecific males. Both parental species mostly have six DAPI-staining bodies, consistent with six diploid chromosomes (like *C. elegans*). Most fertile F1 females also have six DAPI-staining bodies, indicating they are diploid (Fig 2C and Fig S4).

### Asexually-produced F1 inherit two random homologous chromatids from each maternal *C. nouraguensis* bivalent

There are three mechanisms by which the fertile F1 animals could result from diploid maternal inheritance. They could inherit two chromatids from each maternal bivalent in a *C. nouraguensis* oocyte, inherit one chromatid from each bivalent and undergo genome-wide diploidization by endoreplication, or inherit the original two maternal chromatids by producing diploid eggs by mitosis (apomixis). To distinguish between these possibilities, we used whole-genome sequencing of individual worms to determine the genetic identity of the chromatids inherited by fertile F1 individuals.

All four of the chromatids within a bivalent are genetically distinct; of the six possible ways of combining two chromatids, four have distinct chromosomal genotypes and two are indistinguishable by Illumina whole-genome sequencing (Fig 3A, 3B and Fig S5A). Thus, the genotype of the fertile F1 can indicate which two chromatids were inherited. Because recombination in *Caenorhabditis* species occurs predominantly on chromosome arms [26, 27], inheriting two sister chromatids would usually result in homozygosity in the centers of chromosomes and heterozygosity on one of the arms, whereas inheriting two homologous chromatids would result in heterozygosity in the center of each chromosome (Fig 3B). By contrast, endoreplication of single chromatids would lead to homozygosity along all chromosomes, and apomixis would lead to heterozygosity along all chromosomes (Fig S6).

**Fig 3.**
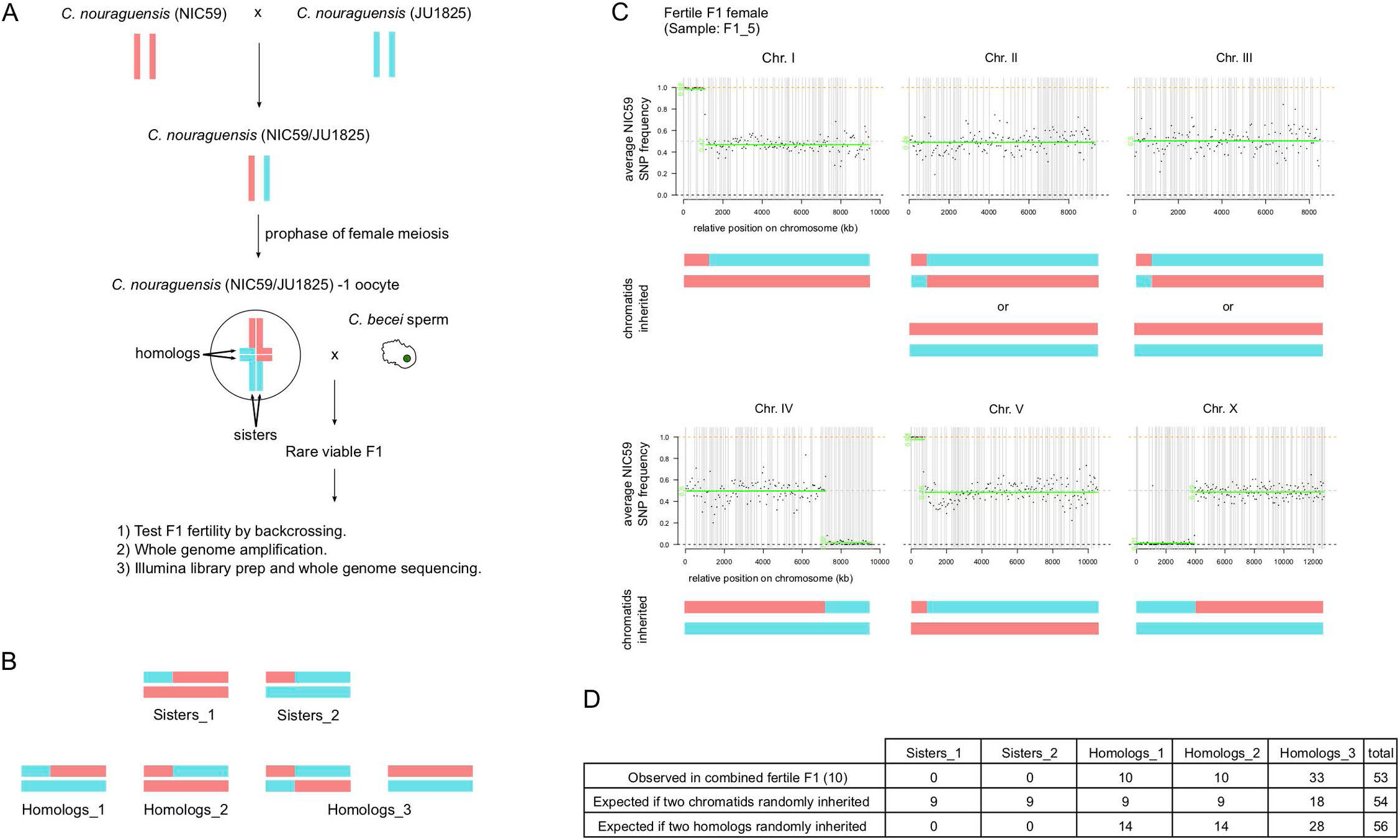
Asexually-produced fertile interspecific F1 inherit two randomly selected homologous chromatids from each maternal bivalent. **(A)** Schematic of how we determined which two chromatids are inherited from each maternal *C. nouraguensis* bivalent. Two genetically distinct strains of *C. nouraguensis*, NIC59 and JU1825, were crossed to make heterozygous NIC59/JU1825 females, which were then crossed to *C. becei* (QG711) males. Viable F1 progeny were collected, assayed for fertility by backcrossing, and prepared for whole-genome amplification and sequencing. **(B)** The six possible ways to inherit two chromatids from a bivalent. Four have distinct chromosomal genotypes (Sisters_1, Sisters_2, Homologs_1 and Homologs_2), and two have genotypes that cannot be distinguished in our short-read sequencing data because they both result in heterozygosity across the entire chromosome (Homologs_3). **(C)** An example of whole-genome genotyping data from a single fertile F1 female (F1_5). Each plot represents one of the six chromosomes. Each point represents the average NIC59 SNP frequency in 50-kb windows ordered along the chromosome; haplotype change points and average allele frequency for each segment are shown by the green horizontal lines. The reference assembly is fragmented into scaffolds that we ordered and oriented on chromosomes using synteny; gray vertical lines represent breaks between scaffolds. The combination of two chromatids that best matches the genotyping data is shown underneath each plot. Genotyping and coverage plots for all F1 are shown in Figs S5 and S7. One fertile F1 gave ambiguous genotypes and was excluded from further analysis (F1_25); this individual had a particularly high percentage of contaminating bacterial reads and low coverage of the *C. nouraguensis* genome (Table S1). **(D)** Frequency of the five distinguishable chromosome genotypes when combining all fertile F1 genotyping data. The table also shows the expected frequency of the five genotypes if any two chromatids were randomly inherited and if any two homologous chromatids were inherited. The frequencies observed in the fertile F1 are not different from those expected from random inheritance of two homologous chromatids (Fisher’s exact test, P=0.41). Hemizygous X chromosomes in males and triploid autosomes were excluded from the analysis.

We first crossed two genetically distinct *C. nouraguensis* strains, NIC59 and JU1825, to generate heterozygous *C. nouraguensis* (NIC59/JU1825) females, which we then crossed to *C. becei* (QG711) males. We collected eleven fertile F1 and performed whole-genome amplification and sequencing of each one (Fig 3A). We determined the genotype of each fertile individual’s chromosomes by calculating average NIC59 SNP frequencies in 50-kb windows across their genome (see Materials and Methods for details). An approximately 0.50 NIC59 allele frequency was inferred as heterozygosity (NIC59/JU1825), whereas an approximately 1.0 or 0.0 NIC59 allele frequency was inferred as homozygous NIC59 or JU1825, respectively. This experiment also allowed us to determine whether fertile F1 inherit only maternal *C. nouraguensis* DNA or whether some paternal *C. becei* DNA is also present.

We found that nearly 100% of each fertile F1’s reads map to the *C. nouraguensis* genome assembly; only a small percentage (0.1-0.2%) map to the *C. becei* genome assembly (Table S1 and Fig S7). These results resemble our controls that contain only *C. nouraguensis* DNA (samples: F1_NIC59_JU1825 and NIC59plusJU1825, Table S1 and Fig S7). Therefore, our sequencing results confirm that fertile F1 inherit only maternal *C. nouraguensis* DNA. Strikingly, whole-chromosome genotyping data shows that the fertile F1 always inherited two homologous chromatids from each maternal bivalent (Fig 3C, Fig S5 and Fig S7). Combining chromosome genotypes from the fertile F1 indicates that two randomly selected homologous chromatids were inherited (Fig 3D). Based on these findings, we can rule out endoreplication and apomixis, or a failure of the second meiotic division that would result in inheritance of two sister chromatids. Instead, we conclude that diploid, asexually-produced F1 inherit two homologous chromatids from each maternal bivalent in *C. nouraguensis* oocytes.

### Sterile F1 inherit a diploid *C. nouraguensis* genome and a haploid *C. becei* genome

Having established that fertile asexually-produced F1 adults inherit only a diploid maternal genome, we next examined the genome-wide inheritance pattern in sterile hybrid F1 adults. We performed whole-genome sequencing of ten sterile F1 adults derived from the same interspecies crosses mentioned in the previous section (Fig 3A).

Surprisingly, five of the sterile F1 appear to be triploid hybrids that inherited a diploid *C. nouraguensis* maternal genome and a haploid *C. becei* paternal genome (females F1_6, F1_12, F1_20; males F1_17 and F1_21, Table S1). In these individuals, the normalized coverage of *C. nouraguensis* autosomes was approximately double that of the *C. becei* autosomes (Fig 4A). Furthermore, genotyping of the *C. nouraguensis* chromosomes revealed that these individuals inherited two homologous chromatids from each maternal *C. nouraguensis* bivalent, just like the fertile F1 (Fig 4B and Fig S7). Three other sterile F1 males are also hybrids that inherited two homologous chromatids from each maternal bivalent. However, they are unlikely to be true, full triploids because the normalized coverage of the *C. becei* autosomes is considerably less than half that of the *C. nouraguensis* autosomes (males: F1_10, F1_16 and F1_23, Fig 4A). Instead, we hypothesize that the reduced coverage of the *C. becei* genome results from mosaicism, in which only a subset of cells in these individuals contain the paternal *C. becei* DNA. The *C. becei* autosomes within each individual hybrid mosaic exhibit similar levels of coverage (except perhaps Chr. V of F1_16), suggesting that most hybrid cells within these individuals have a complete haploid copy of the *C. becei* autosomal genome (diploid-triploid mosaic hybrids) (Fig 4A). By scoring viable animals, we may be selecting against individuals that have partial losses of the paternal genome since such animals would be aneuploid in at least some cells. Alternatively, the cellular mechanism of paternal genome loss may lead to loss of the complete paternal set of chromosomes.

**Fig 4.**
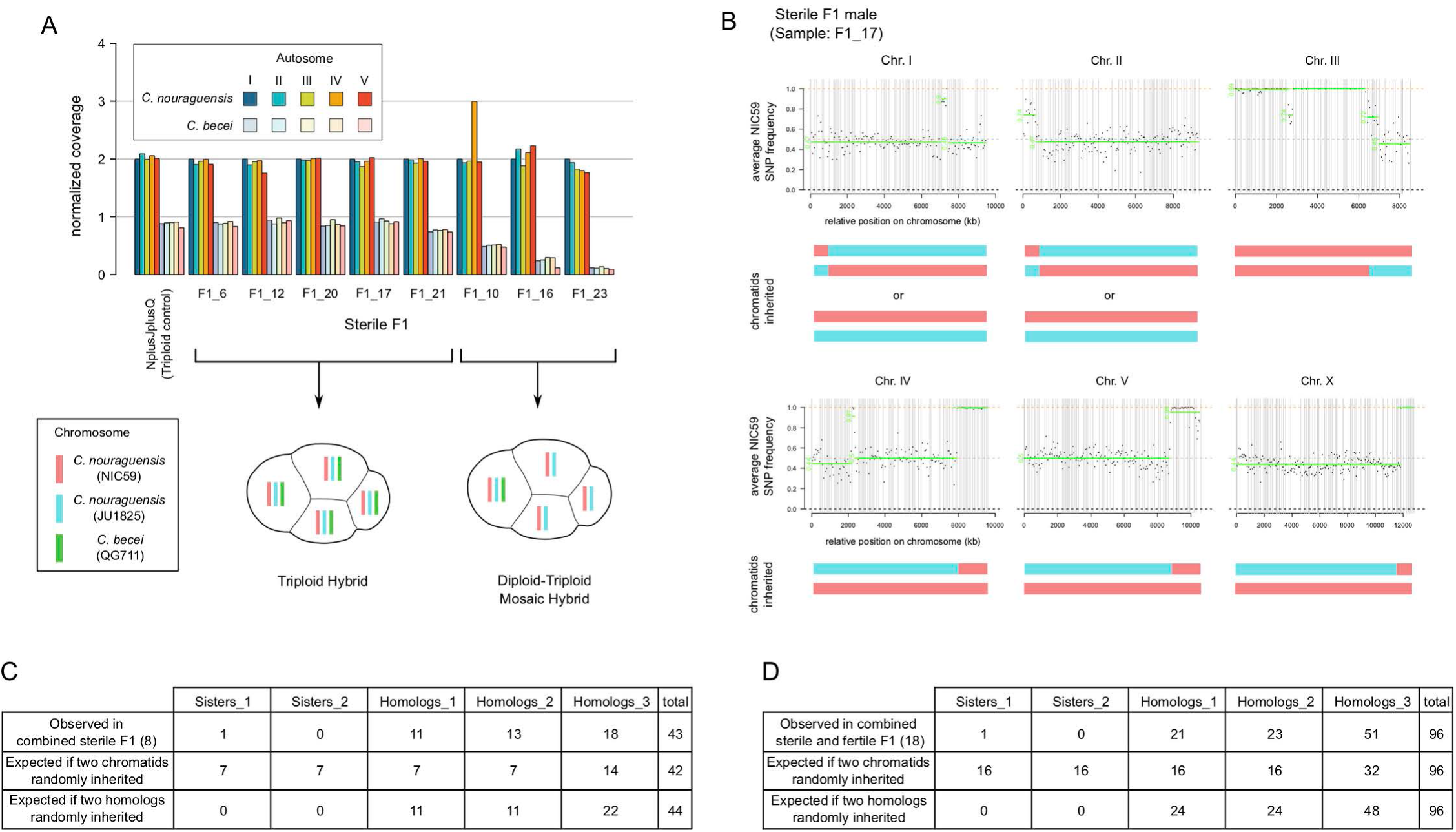
Sterile interspecific F1 inherit a diploid *C. nouraguensis* genome and a haploid *C. becei* genome. **(A)** The normalized read coverage for all autosomes in eight sterile F1 with a clear genotype. Coverage is normalized to the *C. nouraguensis* chromosome I coverage in each individual, which is set to two. Schematics below the graph show sterile F1 that are fully triploid hybrids (all cells of an F1 embryo have a diploid *C. nouraguensis* genome and a haploid *C. becei* genome) or diploid-triploid mosaic hybrids (all cells of an F1 embryo have a diploid *C. nouraguensis* genome, but only a subset of cells inherit the haploid *C. becei* genome). **(B)** An example of whole-genome *C. nouraguensis* genotyping data from a single sterile F1 male (F1_17). Each plot represents one of the six chromosomes. Each point represents the average NIC59 SNP frequency (after removing *C. becei* reads) in 50-kb windows along the physical length of the chromosome; haplotype change points and average allele frequency for each segment are shown by the green horizontal lines. The gray vertical lines represent breaks between scaffolds. The combination of two *C. nouraguensis* chromatids that best matches the genotyping data is shown underneath each plot. Genotyping and plots for all sterile F1 are shown in Figs S5 and S7. We excluded two sterile F1 from further analysis because they gave ambiguous genotypes (F1_18 and F1_26). **(C)** Frequency of the five distinguishable *C. nouraguensis* chromosome genotypes when combining all sterile F1 genotyping data. The table also shows the expected frequency of the five genotypes if any two chromatids were randomly inherited and if any two homologous chromatids were inherited. The frequencies observed in the sterile F1 are not different from those expected from random inheritance of two homologous chromatids (Fisher’s exact test, P=0.78). **(D)** Frequency of *C. nouraguensis* chromosome genotypes when combining all fertile and sterile F1 genotyping data. The frequencies observed are not different from those expected from random inheritance of two homologous chromatids (Fisher’s exact test, P=0.85).

Combining maternal chromosome genotypes from the sterile F1 indicates that two randomly selected homologous chromatids were inherited (Fig 4C and Fig S5C). Additionally, the combined data on maternal genotype frequencies from both sterile and fertile F1 also indicate random homologous chromatid inheritance (Fig 4D and Fig S5C). These results show that the fertile and sterile F1 share the same mechanism of diploid maternal inheritance and are distinguished by whether they inherit the haploid *C. becei* genome.

### The *C. becei* X-chromosome is toxic to F1 hybrids

In *Caenorhabditis*, males are hemizygous for the X-chromosome (XO) and produce haploid sperm that carry either a single X or no X. Therefore, roughly half of the F1 are expected to inherit the paternal *C. becei* X-chromosome. However, though the sterile F1 inherited *C. becei* autosomes, none inherited the *C. becei* X-chromosome (Fig S7). We collected additional viable F1 hybrids from the interspecies cross and determined whether any inherited the *C. becei* X-chromosome by PCR genotyping of two indel polymorphisms, one X-linked and one autosomal, that distinguish between *C. nouraguensis* and *C. becei* sequences (Fig 5A). We found that none of the F1 inherited the *C. becei* X-chromosome (Fig 5B), regardless of whether they were autosomal hybrids (22 females and 12 males) or had a maternal genotype (10 females).

**Fig 5.**
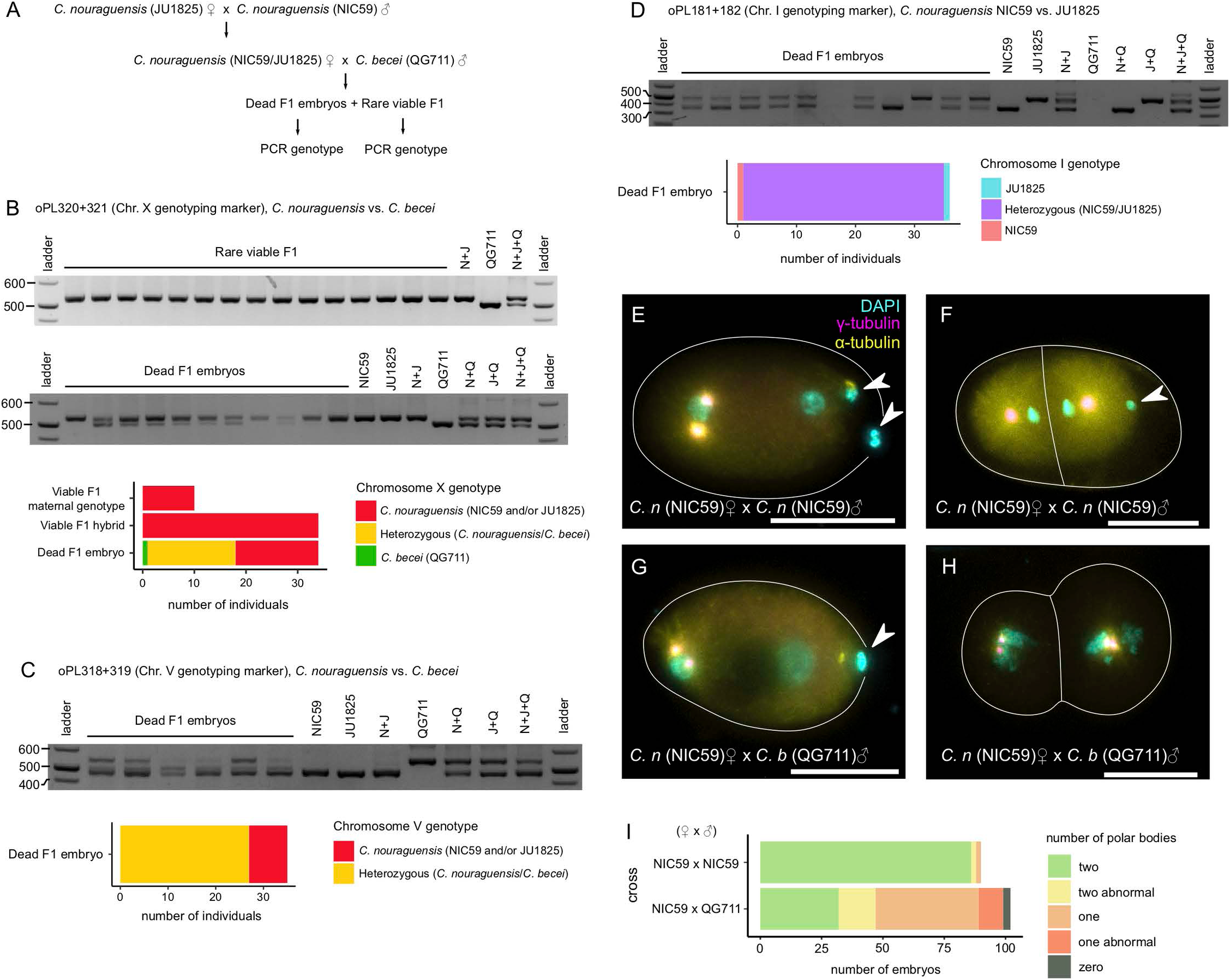
Dead interspecific F1 embryos inherit the *C. becei* X-chromosome and two maternal homologous chromatids. **(A)** Schematic illustrating how dead F1 embryos and viable F1 progeny were generated for PCR genotyping experiments. **(B)** An example of a DNA gel showing the chromosome X genotypes (oPL320+321) of rare viable F1 adults (top gel) and dead F1 embryos (bottom gel). The graph shows that no viable F1 animals inherited the *C. becei* X-chromosome, but half of the dead embryos inherited it. **(C)** An example of a DNA gel showing the chromosome V genotypes (oPL318+319) of dead F1 embryos. The graph shows that most dead F1 embryos inherited chromosome V from both parents but that some inherited only a maternal (*C. nouraguensis*) copy. **(D)** An example of a DNA gel showing the chromosome I genotypes (oPL181+182) of dead F1 embryos. Importantly, the primers used in this PCR reaction do not produce products when using a control *C. becei* QG711 embryo lysate as template, so any signal should be from *C. nouraguensis* templates. The graph shows that almost all dead F1 embryos have a heterozygous NIC59/JU1825 genotype. **(E)** A one-cell stage embryo derived from crossing *C. nouraguensis* (NIC59) females to *C. nouraguensis* (NIC59) males, fixed and stained for DNA (cyan), γ-tubulin (magenta), and α-tubulin (yellow). γ and α-tubulin staining shows the mitotic spindle, which helps determine how many cells there are in the embryo and at what point those cells are in the cell cycle. The white lines outline cells. The two polar bodies that remain associated with the embryo are indicated by arrowheads. **(F)** An intraspecies *C. nouraguensis* embryo with only one polar body. **(G)** A hybrid embryo with only one polar body, derived from crossing *C. nouraguensis* (NIC59) females to *C. becei* (QG711) males. **(H)** A hybrid embryo with zero polar bodies. Scale bars (E-H): 20 µm. **(I)** Roughly half the hybrid embryos have only one polar body. By contrast, embryos derived from *C. nouraguensis* intraspecies crosses almost always have two polar bodies. “Two abnormal” refers to embryos that have two polar bodies, but one or both have an abnormal structure. “One abnormal” refers to embryos that have a single polar body with an abnormal structure.

Our failure to find any viable F1 carrying a *C. becei* X-chromosome suggests that it is toxic to F1 hybrids. Alternatively, this pattern could reflect an unusual segregation pattern specifically involving the X chromosome in hybrids. To distinguish between these possibilities, we PCR genotyped dead F1 embryos from the same interspecies cross. Consistent with the expected inheritance of the X-chromosome from males, we found that 17 of 34 (50%) dead F1 embryos inherited both the *C. becei* paternal X-chromosome and the *C. nouraguensis* maternal X-chromosome, whereas 16 of 34 (47%) inherited only the maternal *C. nouraguensis* X-chromosome (Fig 5B). One dead F1 embryo possessed only a paternal *C. becei* genotype, suggesting that it lost the maternal *C. nouraguensis* X-chromosome. Because dead F1 embryos can inherit the *C. becei* X-chromosome but viable F1 animals do not, we conclude that the *C. becei* X-chromosome is toxic to F1 hybrids. Previous work has described the “large X-effect” on hybrid inviability and sterility [10]. We hypothesize that a dominant incompatibility involving one or more loci on the *C. becei* X chromosome underlies its toxic effect on hybrids with *C. nouraguensis*.

### Dead F1 embryos have unusual maternal and paternal inheritance

To determine whether the unusual patterns of maternal and paternal inheritance seen in viable F1 also occur in the dead F1 embryos in the interspecies cross, we PCR genotyped the same dead F1 embryos described above at an autosomal indel polymorphism that distinguishes the two species (Fig 5C). We found that 27 of the 35 dead F1 embryos (77%) had a heterozygous *C. nouraguensis*/*C. becei* genotype while eight (22%) had an only-maternal *C. nouraguensis* genotype (Fig 5C). These data suggest that although most embryos inherited both maternal and paternal DNA, a significant fraction failed to inherit paternal *C. becei* DNA for this marker. However, given that only a single marker was scored, it is possible that these embryos inherited paternal DNA for other chromosomes and were aneuploid. Aneuploidy caused by incomplete loss of the paternal genome may contribute to death of embryos.

To determine whether dead F1 embryos also inherit two homologous *C. nouraguensis* chromatids from each maternal bivalent, we PCR genotyped them at an autosomal indel polymorphism in the center of chromosome I that distinguishes NIC59 and JU1825 alleles. We found that 34 of 36 dead F1 embryos had a heterozygous NIC59/JU1825 genotype, whereas one embryo had a JU1825 genotype and one embryo had a NIC59 genotype (Fig 5D). These data suggest that very few embryos are derived from a canonical female meiosis in which a haploid maternal complement is inherited. Instead, most embryos appear to have inherited two homologous chromatids from an aberrant female meiosis.

To better understand how female meiosis is modified so that two homologous chromatids are inherited, we quantified the number of polar bodies found in F1 embryos, of which the vast majority are expected to die. If most *C. nouraguensis* oocytes fertilized by *C. becei* sperm have aberrant meiotic segregations that result in the inheritance of two homologous chromatids, we would predict that fewer than two polar bodies are formed. We DAPI stained F1 embryos derived from crossing *C. nouraguensis* (NIC59) females to *C. becei* (QG711) males. As a control, we stained embryos derived from intraspecies NIC59 crosses. As expected, NIC59 embryos almost always have two polar bodies whose morphology and size appear similar across individuals (Fig 5E and 5I). Interestingly, we found that 2/90 NIC59 embryos had only one polar body (Fig 5F and 5I), suggesting that abnormal meiosis can occur at a low frequency in intraspecies crosses. By contrast, approximately half of the hybrid embryos have only one polar body (52/102), suggesting that one of the two female meiotic divisions has failed (Fig 5G-I). The polar bodies in hybrid embryos can also appear abnormal in their morphology (Fig 5I). Together with our genotyping data, these observations indicate that meiotic segregations are frequently perturbed in *C. nouraguensis* females mated to *C. becei* males.

### Hybrid embryos inherit sperm centrioles and DNA, but exhibit mitotic defects

During fertilization in *Caenorhabditis,* the sperm deposits centrioles and a condensed paternal genome into the egg where it forms a pronucleus (Fig 6A). The paternal pronucleus then decondenses and migrates to join the maternal pronucleus for the first mitotic division (Fig 6B-E) [28]. In obligately gynogenetic species, the paternal genome can be lost by remaining condensed during the first few mitotic divisions of the zygote rather than being replicated and segregated properly [29, 30]. To determine how the paternal genome behaves in hybrid embryos, we stained *C. nouraguensis* x *C. becei* F1 hybrid embryos with DAPI (staining DNA) and antibodies to α-tubulin (staining spindles) and γ-tubulin (staining centrosomes). We found that hybrid embryos always have centrioles and a paternal genome derived from the sperm as is normal in *Caenorhabditis*, but in a small fraction of animals, the centrioles are abnormally dissociated from the male pronucleus (Fig 6M and 6Q). We consistently observed both paternal and maternal pronuclei decondensing and meeting (Fig 6I-L and 6Q). Thus, unlike obligate gynogenetic species, there is not frequent failure of the paternal genome to decondense in these hybrid embryos. However, it is possible that the paternal genome fails to decondense at a low frequency and that those embryos are the ones that exhibit clean paternal genome loss and become fertile adults.

**Fig 6.**
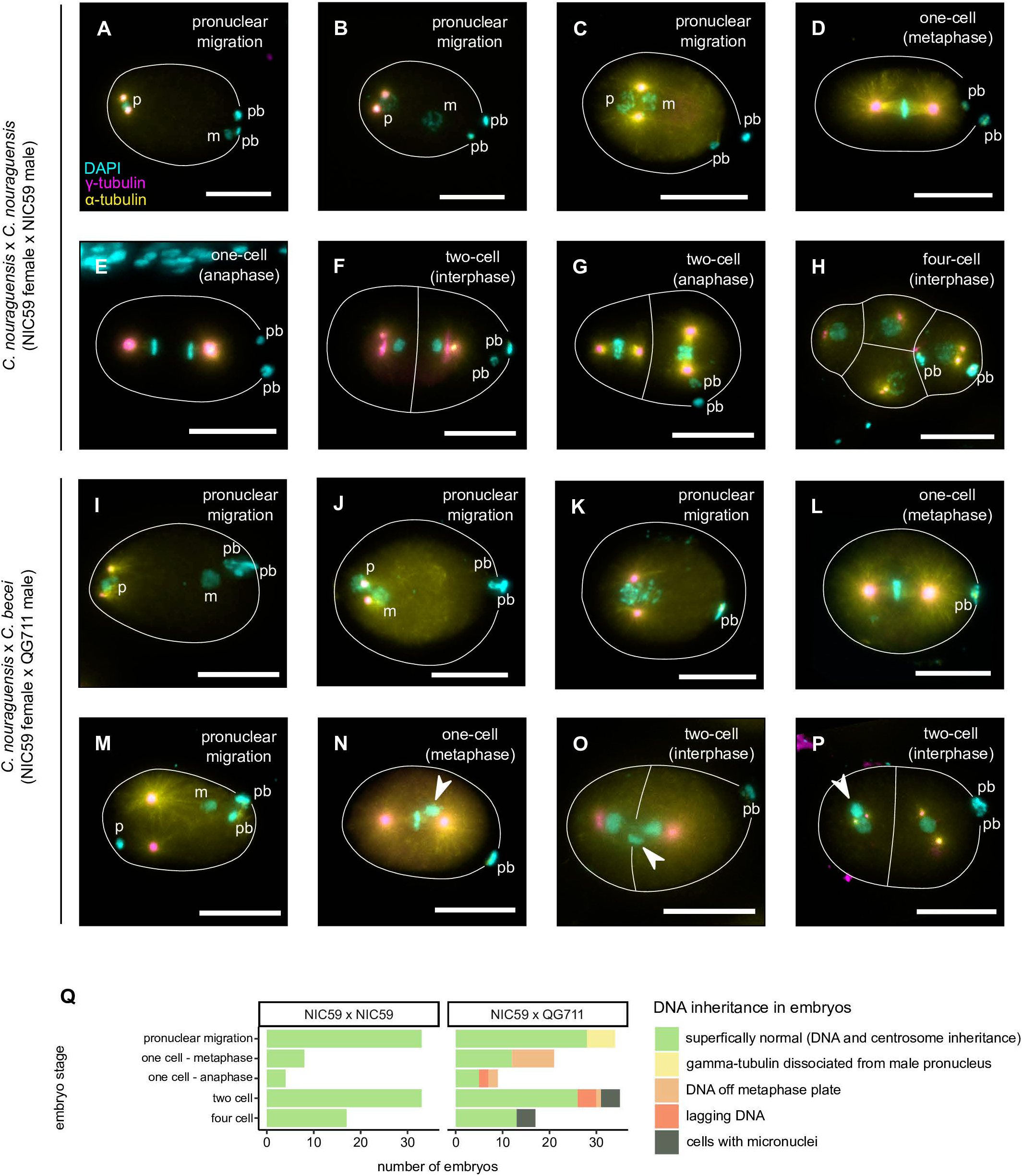
Cytological characterization of early embryonic development in interspecies hybrids. **(A-H)** A set of fixed embryos that summarize early embryonic development in *C. nouraguensis* intraspecies crosses (NIC59 female x NIC59 male). Embryos stained for DNA (cyan), α-tubulin (yellow) and γ-tubulin (magenta). Scale bars: 20 µm. **(A)** When the sperm fertilizes the oocyte, it deposits a condensed haploid paternal genome (“p”) and centrioles into the egg. The centrioles gather γ-tubulin, which nucleates microtubule polymerization (two magenta dots). Fertilization triggers the female meiotic divisions, resulting in two polar bodies (“pb”) and the formation of a haploid maternal pronucleus (“m”). **(B)** The two pronuclei decondense and migrate towards each other. Centrosomes remain associated with the paternal pronucleus. **(C)** The two pronuclei meet. **(D)** Maternal and paternal chromosomes align along the metaphase plate of the first mitotic spindle. **(E)** Sister chromatids segregate to opposite spindle poles during anaphase and **(F)** form interphase nuclei after cytokinesis. **(G)** A two-cell embryo with both cells undergoing anaphase. **(H)** A four-cell stage embryo with all cells in interphase. **(I-P)** Hybrid embryos derived from *C. nouraguensis* (NIC59) female x *C. becei* (QG711) crosses. **(I-L)** Hybrid embryos that do not exhibit abnormalities in centrosome inheritance or pronuclear decondensation and migration **(M-P)** Hybrid embryos that exhibit abnormalities during early embryogenesis. **(M)** Centrosomes in hybrid embryos can be dissociated from the male pronucleus during pronuclear migration. Both pronuclei are condensed. **(N)** A one-cell hybrid embryo with misaligned DNA at metaphase (arrowhead). **(O)** A two-cell hybrid embryo with lagging DNA between the two interphase cells (arrowhead). **(P)** A two-cell hybrid embryo during interphase. One cell has an extra smaller nucleus (micronucleus, arrowhead). **(Q)** Quantification of embryos with abnormalities like those shown in Fig 6M-P.

We also observed later stages of hybrid embryogenesis from metaphase of the first mitosis to the four-cell stage. We found that a fraction of hybrid embryos exhibit DNA that is not properly aligned on the mitotic spindle (Fig 6N and 6Q), lagging DNA during or after anaphase (Fig 6O and 6Q), and probable micronuclei during interphase (Fig 6P and 6Q). Although it is unclear whether the aberrant DNA is maternal or paternal, these data are consistent with some paternal genome loss at later embryonic stages.

### Diploid maternal inheritance occurs occasionally in *C. nouraguensis* intraspecies crosses

Because we observed a low frequency of embryos with a single polar body in normal intraspecies *C. nouraguensis* crosses (2/90, Fig 5F and 5I), we wondered whether diploid maternal inheritance could occur at low frequency within *C. nouraguensis* independently of interspecies hybridization. To more sensitively and directly assay for diploid maternal inheritance within *C. nouraguensis*, we designed an experiment in which *C. nouraguensis* sperm can fertilize *C. nouraguensis* oocytes but do not contribute the paternal genome. In this scenario, a canonical female meiosis would result in haploid maternal embryos that would fail to develop into viable adults [20]. However, if *C. nouraguensis* oocytes inherit a diploid maternal genome, they could complete embryogenesis and develop into viable adults.

We blocked the inheritance of the haploid sperm genome using UV irradiation to damage and inactivate the sperm genome while still allowing for fertilization and development of the maternal haploid embryo [31, 32]. We exposed NIC59 and JU1825 adult males to a high dose of shortwave UV radiation and then crossed them to non-irradiated JU1825 or NIC59 adult virgin females, respectively. We then screened for and PCR genotyped viable progeny at several autosomal loci to determine whether they inherited both maternal and paternal DNA (heterozygous NIC59/JU1825) or just maternal DNA (Fig 7A).

**Fig 7.**
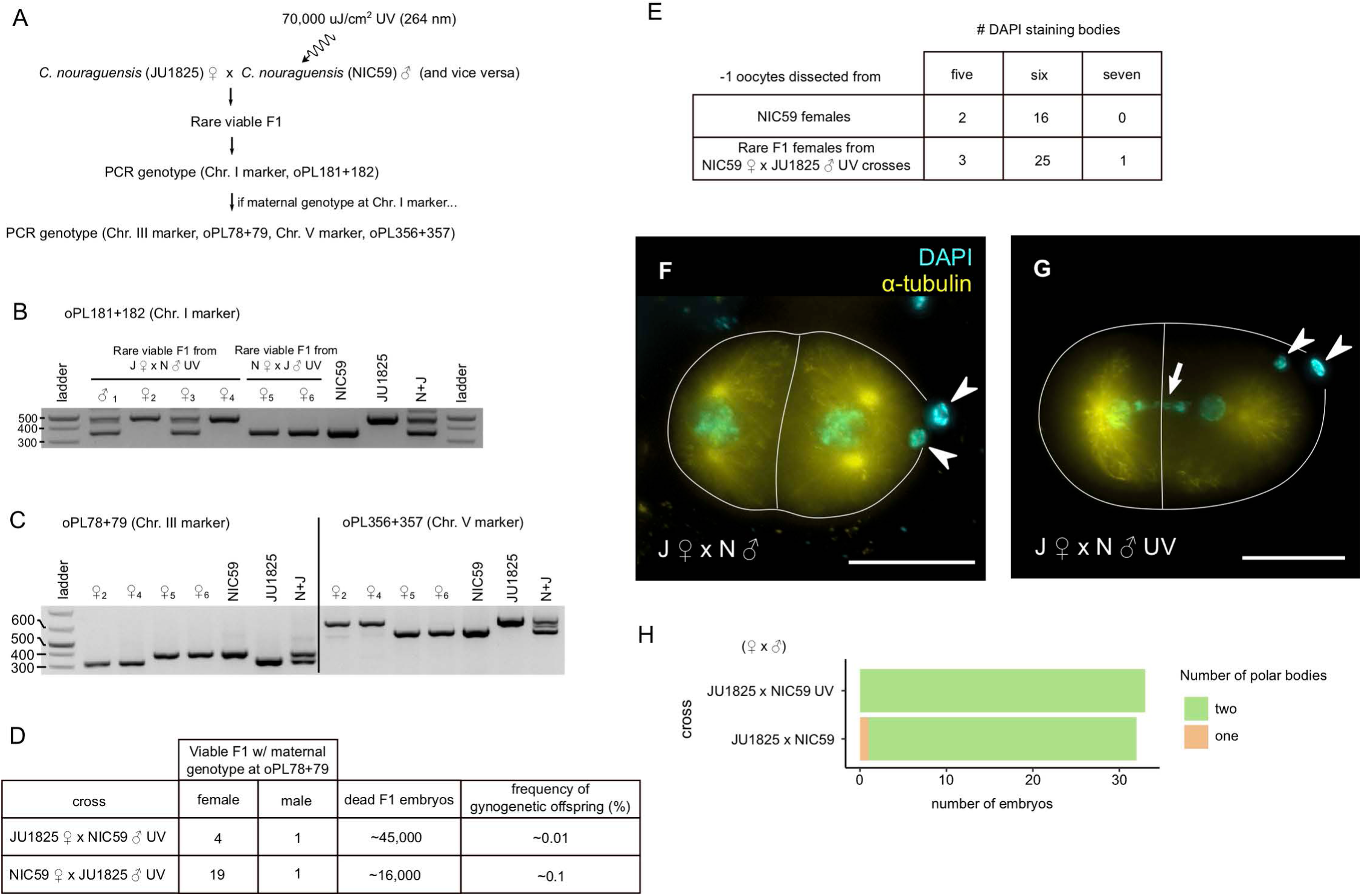
Diploid maternal inheritance can occur independently of interspecies hybridization. **(A)** Flowchart illustrating how the UV irradiation experiments were conducted. **(B)** A DNA gel showing the chromosome I genotypes (oPL181+182) of several rare viable F1 derived from crossing either *C. nouraguensis* JU1825 females to UV-irradiated *C. nouraguensis* NIC59 males (J♀ x N♂ UV) or NIC59 females to UV-irradiated JU1825 males (N♀ x J♂ UV). The sex of the rare viable F1 is depicted above each lane, with an identifying number as subscript. **(C)** A DNA gel showing the chromosome III and V genotypes (oPL78+79 and oPL356+357) of the rare F1 from Fig 7B that had a maternal genotype for chromosome I. All also had a maternal genotype at these two markers. **(D)** A table quantifying the frequency of gynogenetically-produced offspring from all *C. nouraguensis* intraspecies UV experiments (two for each cross). **(E)** A table quantifying the number of DAPI staining bodies in the −1 oocytes of gynogenetically-produced females from *C. nouraguensis* intraspecies UV experiments. **(F)** A two-cell embryo derived from an intraspecies *C. nouraguensis* cross (JU1825♀ x NIC59♂). Two polar bodies are indicated by white arrowheads. White lines outline cells. Scale bar: 20 µm. **(G)** A two-cell embryo derived from an intraspecies *C. nouraguensis* cross with irradiated males (JU1825♀ x NIC59♂ UV). Two polar bodies are indicated by white arrowheads. Lagging DNA (likely UV-irradiated paternal DNA) is indicated by the white arrow. **(H)** Quantification of polar bodies in *C. nouraguensis* intraspecies crosses.

From a cross of JU1825 females to NIC59 UV-irradiated males, only seven out of approximately 18,000 embryos survived to adulthood. Upon PCR genotyping a chromosome I indel polymorphism, we found that four of these seven individuals had a heterozygous NIC59/JU1825 genotype and therefore inherited both maternal and paternal alleles at this locus. We presume these animals represent rare cases in which the sperm genome was not destroyed. However, three females inherited only the maternal JU1825 allele (Fig 7B). We PCR genotyped these females at two other unlinked autosomal indel polymorphisms. All had a maternal JU1825 genotype at all three loci, suggesting that they inherited only the maternal genome and were asexually-produced (Fig 7C). We performed similar experiments with NIC59 females and UV-irradiated JU1825 males. Only two out of roughly 7,000 progeny survived to adulthood. These two females had only-maternal NIC59 genotypes at all three autosomal loci (Fig 7B and 7C). We performed additional UV-experiments to better quantify the frequency of gynogenetically-produced offspring. In total, we found that gynogenetically-produced offspring occur at a rate of ∼0.01% in JU1825 female x NIC59 male UV crosses and ∼0.1% in NIC59 female x JU1825 male UV crosses. Interestingly, each cross had a single gynogenetically-produced male (Fig 7D).

We then determined the ploidy of the gynogenetically-produced females from the UV experiments by counting the number of DAPI-staining bodies in their −1 oocytes. We found that almost all had six DAPI-staining bodies, indicating that they result from diploid maternal inheritance (Fig 7E). The genotyping and DAPI staining data together indicate that diploid maternal inheritance in *C. nouraguensis* oocytes does not require fertilization by *C. becei* sperm and can occur even in intraspecies crosses, at least when the males are sterilized by UV.

A caveat to the UV experiments is that they may not be indicative of what occurs during normal non-irradiated intraspecies crosses. For instance, diploid maternal inheritance could be artificially induced by fertilization with UV-damaged sperm. To determine whether female meiosis is perturbed when oocytes are fertilized by UV-irradiated sperm, we stained embryos derived from JU1825 female x NIC59 male UV crosses. We found that the embryos we scored all exhibited two superficially normal polar bodies, much like the non-UV control (Fig 7F-H). This result suggests that female meiosis is usually normal when an oocyte is fertilized by UV-damaged sperm. Furthermore, PCR genotyping of dead embryos derived from crossing heterozygous NIC59/JU1825 females to NIC59 UV-irradiated males is consistent with haploid maternal inheritance and destruction of the paternal NIC59 genome (Fig S8). Therefore, our data suggest that female meiosis is not generally perturbed by fertilization with UV-irradiated sperm, and that the diploid maternal inheritance seen in intraspecies crosses is not an artificial byproduct of a grossly disrupted meiosis. In support of this, we observe occasional embryos derived from normal intraspecies crosses with only a single polar body (Fig 5F and 5I, and Fig 7H), consistent with diploid maternal inheritance occurring in *C. nouraguensis* at a low frequency independently of UV-irradiation.

## DISCUSSION

In this study we investigated the hybridization of two sexual *Caenorhabditis* species, *C. nouraguensis* and *C. becei*. We found that rare viable offspring are produced in the cross of *C. nouraguensis* females to *C. becei* males. About one-third of the viable offspring are fertile and result from a combination of diploid maternal inheritance and paternal genome loss, two traits that define asexual gynogenetic reproduction. The remaining viable offspring are sterile and possess a diploid maternal *C. nouraguensis* genome and a haploid paternal *C. becei* genome. Furthermore, dead hybrid embryos also inherit two homologous maternal chromosomes, like the viable offspring, indicating that diploid maternal inheritance is a general feature of this cross. Finally, we found that diploid maternal inheritance can also occur at a low frequency in intraspecies *C. nouraguensis* crosses, as revealed by UV-irradiation of males. These data suggest that meiosis may be inherently fragile in *C. nouraguensis* oocytes and primed for diploid maternal inheritance. However, the frequency of diploid maternal inheritance is drastically increased when these oocytes are fertilized by *C. becei* sperm, suggesting that unknown factors present in *C. becei* sperm somehow alter *C. nouraguensis* female meiosis.

### Mechanism of diploid maternal inheritance

We found that diploid maternal inheritance in fertile and sterile offspring almost always results from inheriting two randomly selected homologous chromatids from each maternal bivalent in the oocyte. We hypothesize that this diploid maternal inheritance reflects a stereotypical segregation mechanism that is a slight modification of canonical meiotic divisions (Fig 8A).

**Fig 8.**
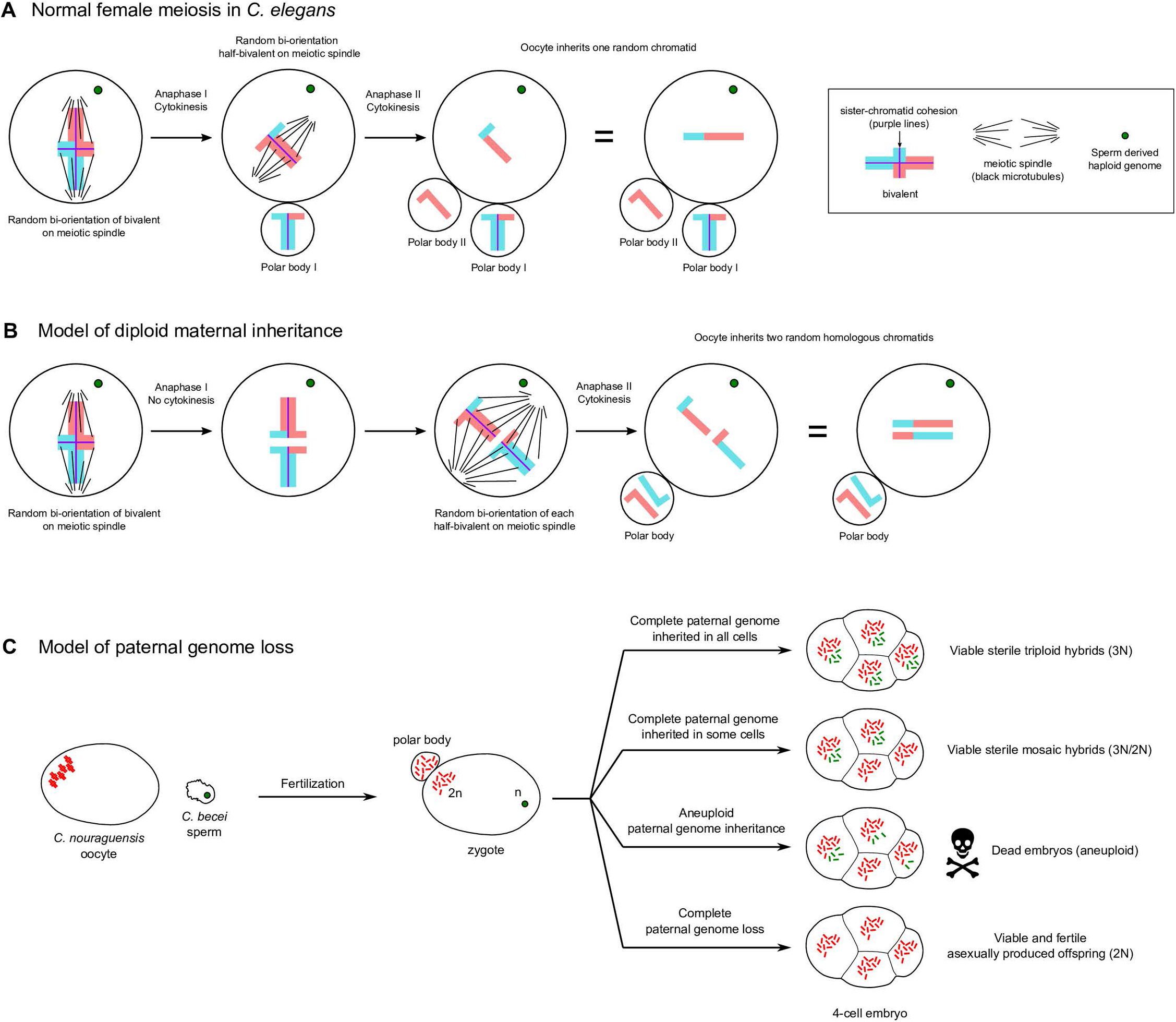
Models for diploid maternal inheritance and paternal genome loss in *C. nouraguensis* oocytes. **(A)** Schematic of canonical meiotic divisions in *C. elegans*. Upon fertilization, each maternal bivalent (only one shown here) randomly bi-orients its homologs on the meiotic spindle, cohesion is lost between homologous chromosomes and homologs segregate. One set of homologs segregates into the first polar body while the other is retained in the oocyte (Anaphase I and Cytokinesis). Then the half-bivalent in the oocyte randomly bi-orients on the meiotic spindle, sister chromatid cohesion is lost and sister chromatids segregate. One sister chromatid is segregated into the second polar body while the other is retained in the oocyte (Anaphase II and Cytokinesis). Thus, the oocyte inherits only one random chromatid from each bivalent. **(B)** One model for how female meiosis could be modified to inherit two random homologous chromatids from a bivalent. Upon fertilization, the bivalent randomly bi-orients its homologs on the meiotic spindle and cohesion is lost between homologous chromosomes as is normal. Homologs segregate but cytokinesis fails (Anaphase I) and both half-bivalents remain in the oocyte. Each half-bivalent then bi-orients on the meiotic spindle, sister chromatid cohesion is lost, and sister chromatids segregate. One chromatid from each half-bivalent segregates into the second polar body while the other is retained in the oocyte (Anaphase II and Cytokinesis). Thus, the oocyte inherits two random homologous chromatids from a bivalent. **(C)** Model of paternal genome loss in hybrids between *C. nouraguensis* females and *C. becei* males. In nearly all cases, a diploid egg is produced, but the fate of the embryo depends on the outcome of paternal genome elimination. In the rare viable offspring the entire haploid *C. becei* paternal genome except for the X chromosome is inherited in all cells (sterile triploid hybrids), only a subset of cells (sterile diploid-triploid mosaic hybrids), or in no cells (fertile asexually-produced offspring). The vast majority of the offspring are dead embryos that have partial losses of the paternal genome in at least some cells and are aneuploid. Approximately half of the dead embryos inherit the *C. becei* X chromosome that is toxic to hybrids.

We propose that gynogenetic development in *C. nouraguensis* results from an alteration of the first meiotic division followed by a normal second meiotic division. In our model (Fig 8B), homologous chromosomes segregate normally at anaphase I, but instead of segregating one set of homologs into the first polar body, both sets are retained in the oocyte. Both half-bivalents within the oocyte then randomly segregate sister chromatids at anaphase II. One chromatid segregates into a polar body while the other is retained in the oocyte. Thus, the oocyte inherits two randomly selected homologous chromatids from each bivalent. This model suggests that diploid maternal inheritance could result from a modification of female meiosis such as failure of cytokinesis after anaphase I. There is cytological evidence that incomplete anaphase I and a failure of cytokinesis generates diploid eggs in *Daphnia pulex*, which undergoes obligate parthenogenesis [33].

Importantly, our model predicts the formation of only one polar body (Fig 8B). Approximately half of the dead hybrid embryos have only one polar body (Fig 5G and 5I). Combined with the fact that dead hybrid embryos also inherit two maternal homologs (Fig 5D), our data suggest these embryos also undergo the same mechanism of diploid maternal inheritance as the viable progeny. But if almost all embryos have diploid maternal inheritance (Fig 5D), why do approximately half still have two polar bodies? One possibility is that the two half-bivalents formed by the incomplete first meiotic division are sometimes segregated on the same spindle at meiosis II, resulting in one polar body, but are sometimes segregated on separate spindles for meiosis II, resulting in two polar bodies.

How might meiosis I be disrupted to produce diploid maternal inheritance? Some loss-of-function mutations and gene knockdowns in *C. elegans* can possibly lead to diploid maternal inheritance [34, 35], suggesting plausible cellular mechanisms underlying this phenomenon. During normal meiosis I in *C. elegans*, the meiotic spindle migrates and orients perpendicularly to the oocyte cortex. In anaphase I, one set of homologs segregates towards the cortex into the first polar body while the other is retained in the oocyte. However, weak knockdown of dynein heavy chain (*dhc-1*) causes the meiotic spindle to be oriented parallel to the cortex rather than perpendicular during meiosis I; as a result, homologous chromosomes segregate parallel to the cortex in anaphase I and no polar body is formed [35]. The half-bivalents that remain in the oocyte then segregate relatively normally during anaphase II and form a polar body. The oocyte presumably inherits two randomly selected homologous chromatids from each bivalent. A similar misalignment of the meiotic spindle at metaphase I is also thought to be the mechanism underlying the generation of diploid eggs in the obligately parthenogenetic species *Drosophila mangabeirai* [36]. We hypothesize that a misalignment of the meiotic spindle or a failure of cytokinesis may similarly result in diploid maternal inheritance in *C. nouraguensis*.

What are the evolutionary consequences of this type of diploid maternal inheritance? We have shown that diploid maternal inheritance in *C. nouraguensis* oocytes is the result of inheriting two homologous chromatids that have undergone meiotic recombination, and therefore fits a model of automixis in which two homologous chromatids are combined (“central fusion”). Because there is only a single recombination event that is biased towards the chromosome ends, inheriting two homologous chromatids either maintains heterozygosity across the entire chromosome or most of it (Fig 3B). However, recombination does occur in the middle of chromosomes at a lower frequency [26], which should eventually result in genome-wide homozygosity and inbreeding depression after many generations of automixis. Most *Caenorhabditis* species are dioecious (females and males), with some exhibiting genetic hyperdiversity [37, 38]. Therefore, inbreeding depression could be a sizable hurdle to overcome for the persistence of this form of gynogenesis evolving from an obligately sexual *Caenorhabditis* ancestor.

### Females can produce males asexually

Given that two copies of each maternal autosome are almost always inherited in the interspecies *C. nouraguensis* x *C. becei* cross, we were surprised to observe that some diploid fertile F1 are males. Indeed, sequencing data show that the three fertile males have only a single X chromosome (Fig S5 and Fig S7). Thus, missegregation of the X-chromosome during the modified female meiosis can result in only one X chromatid being inherited by the oocyte. Interestingly, two of the three fertile males were also aneuploid (triploid) for a single *C. nouraguensis* autosome, indicating that widespread chromosome missegregation occurred while producing these individuals. Generating males at a low frequency can theoretically aid the spread and maintenance of gynogenesis by transmitting alleles required for diploid maternal inheritance into neighboring sexual populations [39]. The production of males even at a low frequency can also help propagate an intraspecific gynogenetic lineage by enabling the activation of the egg and inheritance of centrioles without relying on males from a related species. This is observed in the nematode *Mesorhabditis belari,* in which only 9% of offspring are males and 91% are gynogenetically reproducing females [30]. Thus, it is notable that the modified female meiotic segregations that facilitate asexual reproduction also generate males that could promote the long-term maintenance of an asexual lineage.

### Paternal genome loss

In addition to diploid maternal inheritance, gynogenesis also requires paternal genome loss. In viable F1 offspring from the *C. nouraguensis* x *C. becei* cross, we found that the entire haploid *C. becei* paternal genome can be inherited in all cells (triploid hybrids), only a subset of cells (diploid-triploid mosaic hybrids), or in no cells (fertile asexually-produced offspring) (Fig 8C). Given the low frequency of viable offspring, it appears that complete loss of the paternal genome (resulting in fertile offspring) or complete retention of the paternal genome (resulting in sterile triploid offspring) are low frequency events. Rather, the majority of the animals likely have partial losses of the paternal genome that result in aneuploidy and death of embryos.

Interestingly, the offspring of naturally gynogenetic fish species can have a similar spectrum of paternal inheritance. Specifically, most offspring completely fail to inherit the paternal genome (diploid maternal clones), but a small fraction of offspring can inherit it in a subset of cells (diploid-triploid mosaic) or all cells (triploid) [29,40,41]. Given these similarities, the mechanism of paternal genome loss in *C. nouraguensis* interspecies gynogenesis might be similar to that of some obligately gynogenetic fish species. However, paternal genome loss can occur in a variety of contexts and has different underlying causes, each of which could potentially explain the loss of the *C. becei* genome [42–47]. In the context of interspecies hybridization, detailed studies in *Hordeum* (barley) and *Xenopus* show that paternal chromosomes fail to recruit centromeric proteins, lag during embryonic divisions, and form micronuclei [12, 13]. It is noteworthy that we also find lagging DNA and micronuclei in a fraction of dead *C. nouraguensis*/*C. becei* hybrid embryos, potentially indicating that paternal *C. becei* chromosomes fail to load centromeric proteins required for chromosome segregation.

### Asexual reproduction via hybridization

In animals, fertilization by male sperm is required to reactivate the developmental program of oocyte meiosis, and sperm also provide centrioles to the zygote. Thus, the evolution of parthenogenetic asexual reproduction in a sexual female requires that the oocyte not rely on sperm for centrioles and reactivation of meiosis. This barrier can be overcome via gynogenesis in which sperm fertilize oocytes and provide meiotic reactivation and centrioles, but sperm DNA is not inherited. While solving one problem, gynogenesis introduces several new problems. First, in an asexual species, where do the males come from? This problem is solved in some obligately gynogenetic species that are composed entirely of females and require that their eggs be fertilized and activated by males of a closely related species [9]. Thus, hybridization can overcome one of the major barriers to the emergence of asexual reproduction. In the hybridization studied here, fertilization of *C. nouraguensis* oocytes by *C. becei* sperm leads to inheritance of paternal centrioles, an altered oocyte meiosis that produces diploid eggs, and unreliable inheritance of paternal DNA. Gynogenetic reproduction in this cross is rare because complete paternal genome elimination is rare (Fig 8C). We hypothesize that this rare gynogenetic reproduction could serve as a transitional state between sexual reproduction and more reliable interspecies gynogenetic reproduction.

As illustrated above, a second problem for gynogenesis to overcome is the need for the paternal genome to be completely and cleanly eliminated. Here again, hybridization may provide part of the solution if the oocyte cytoplasm of one species does not support the reliable inheritance of the paternal DNA of the other species. Such incompatibility between oocyte cytoplasm and sperm DNA is far more likely between species than within species, providing another possible reason that many asexual species are formed through hybridization. Hybridization may also facilitate diploid maternal inheritance if incompatibilities between sperm and egg disrupt oocyte meiosis and lead to the production of diploid maternal eggs, as occurs in the hybridization between *C. nouraguensis* and *C. becei*. The hybrid incompatibilities observed in this cross support the “balance hypothesis” that suggests how hybridization can facilitate asexual reproduction. On its own, complete paternal genome elimination in the context of normal sexual reproduction would be strongly deleterious as it would lead to the production of lethal haploid progeny. Likewise, diploid maternal inheritance induced by sperm in the context of normal sexual reproduction would lead to the production of triploid progeny that have reduced fitness (sterile in the cross we studied). But in combination, these two incompatibilities can lead to the production of viable asexually-produced progeny.

In conclusion, our study establishes the *C. nouraguensis* – *C. becei* hybridization as a new genetic model system to study how hybrid incompatibilities may facilitate the emergence of rare asexuality in animals. Further study of this system may provide insights into the mechanisms of diploid maternal inheritance and paternal genome loss, as well as the obstacles associated with a transition from sexual to asexual reproduction.

## MATERIALS AND METHODS

### Strain isolation and maintenance

All strains of *Caenorhabditis* used in this study were derived from single gravid females isolated from rotten fruit or flowers [48, 49]. Strains of *C. nouraguensis* were kindly provided by Marie-Anne Félix (“JU” prefix) and Christian Braendle (“NIC” prefix). Most strains are outbred, except JU2079, QG2082 and QG2083, which are inbred lines derived from JU1825, QG704 and QG711, respectively. These inbred lines were used to generate the *C. nouraguensis* and *C. becei* genome assemblies. Strain stocks were stored at −80°. Strains were maintained at 25° on standard NGM plates spread with a thin lawn of *E. coli* OP50 bacteria [50].

### Phylogenetic analysis

To construct a phylogeny, we used a subset of the coding sequences from Kiontke et al. [48]. The following genes were used: *ama-1, lin-44, par-6*, *pkc-3*, *W02B12.9*, *Y45G12B.2a*, *ZK686.3*, and *ZK795.3*. The *C. yunquensis* coding sequences were obtained from NCBI, while all other coding sequences were obtained from the caenorhabditis.org BLAST server (blast.caenorhabditis.org/#) using the *C. elegans* homolog as the query sequence. The coding sequences for each species were concatenated and aligned with MUSCLE [51] using default settings. A maximum likelihood phylogeny was made using PhyML 3.0 online (http://www.atgc-montpellier.fr/phyml/) [52] using a Generalized Time Reversible (GTR) substitution model with six substitution rate categories. Tree searching was performed using Nearest Neighbor Interchanges (NNIs). Branch support was determined by 100 bootstraps.

### Quantifying strain viability

Strain viability was quantified as in [53]. Briefly, we crossed 10 virgin L4 females and males of the same strain on a single plate coated with palmitic acid. The plates were placed at 25° overnight, allowing the worms to mature to adulthood and begin mating. The adult worms were then moved to a new plate rimmed with palmitic acid, allowed to mate and lay eggs for 1.5 hours at 25°, and then removed. The eggs laid within that time were immediately counted. Two days later, we placed the plates at 4° for 1 hr and counted the number of healthy L4 larvae and young adults per plate. We defined the percentage of viable progeny as the total number of L4 larvae and young adults divided by the total number of eggs laid.

### DIC imaging of embryogenesis

Twenty-four hours after the L4 stage, we picked 20-40 gravid females into a 30 µl pool of 1x Egg Buffer (25 mM HEPES pH 7.4, 118 mM NaCl, 48 mM KCl, 2 mM EDTA, 0.5 mM EGTA) on a glass coverslip and dissected out embryos by cutting the adults in half using two needles like scissors. The embryos were then transferred with a glass capillary tube into a fresh pool of Egg Buffer to dilute any contaminating bacteria, and then placed onto a 2% agarose pad on a glass slide. The mounted embryos were put into a humid chamber for 20 minutes before adding a coverslip. The edges of the coverslips were sealed with petroleum jelly to prevent evaporation of the agarose pad. DIC z-stack images were captured every 3 minutes for 10 hours using a Nikon Eclipse 80i compound microscope (60x oil lens, 1.40 NA). Images were processed using Image J [54].

### Calculating the frequency of rare interspecific F1

For each interspecies cross we set up 20-25 plates, each with one virgin L4 female crossed to a single virgin L4 male. The plates were placed at 25° overnight, allowing the worms to mature to adulthood and begin mating and laying eggs. To prevent overcrowding, the adult couples were moved to a new plate every day for three days. The number of eggs laid on each plate was counted the day the parents were removed. Each plate was monitored for two days to check for the presence of rare viable F1, which were counted and moved onto a new plate.

### Generating rare interspecific F1 and testing their fertility

To collect enough rare viable F1 animals for fertility testing and genotyping, we set up six cross plates, each with 30 *C. nouraguensis* L4 females and 30 *C. becei* L4 males and let them develop into adults overnight at 25°. We monitored the plates for the presence of viable F1 larvae each day for 5-6 days. Each viable F1 was moved to its own new palmitic acid-rimmed plate, and its fertility was tested as either an L4 or young adult by backcrossing it to one or two *C. nouraguensis* individuals of the opposite sex. F1 individuals were classified as sterile if they mated to *C. nouraguensis* but produced no F2 embryos, while F1 individuals were classified as fertile if they produced F2 embryos. Mating was inferred by the presence of a male-deposited copulatory plug on the female vulva [55]. After fertility testing and PCR genotyping, we found that hybrid males produced no embryos when crossed to *C. nouraguensis* females and were classified as sterile. However, they often failed to produce a copulatory plug, so we cannot tell whether sterility is due to abnormal germline development or defective mating.

To track the fate of the two maternal homologous chromosomes in subsequent crosses, we crossed two genetically distinct strains of *C. nouraguensis* (JU1825 and NIC59) to generate heterozygous *C. nouraguensis* NIC59/JU1825 females. Specifically, we crossed virgin L4 JU1825 females to NIC59 males and vice versa to generate female offspring that are heterozygous NIC59/JU1825 at all nuclear loci and carry either JU1825 or NIC59 mitochondria, respectively. We denote the genotype of these heterozygous females by the following nomenclature: “(mitochondrial genotype); nuclear genotype”. Therefore, the first cross produces *C. nouraguensis* (J); N/J females, while the second produces *C. nouraguensis* (N); N/J females (Table S1). We then crossed both types of *C. nouraguensis* N/J virgin females to *C. becei* QG711 males. We collected the rare viable F1 from these interspecies crosses and tested their fertility by backcrossing them to the *C. nouraguensis* strain matching their mitochondrial genotype in order to avoid known cytoplasmic-nuclear incompatibilities within this species [53].

### Generating worm and embryo lysates for PCR

Single worm PCR: Each adult worm was placed in a PCR tube with 10 µl of lysis buffer (1x Phusion HF buffer + 1.0% Proteinase K) and frozen at −80° for at least 15 minutes. The worms were then lysed by incubating the tubes at 60° for 60 mins and 95° for 15 mins. 1 µl of worm lysate was used in each 10 µl PCR reaction. Control lysates were generated by mixing 10-15 worms in lysis buffer.

Single embryo PCR: Dead embryos were picked into a 30 µl pool of 2 mg/ml chitinase on an 18×18 mm coverslip. OP50 bacteria were washed off embryos by pipetting the chitinase solution up and down. Using a glass capillary tube, embryos were transferred to a new pool of chitinase and washed again, and individual embryos were placed in a PCR tube with 5 µl of 2 mg/ml chitinase solution and incubated at room temperature for 2 hours. 5 µl of lysis buffer (2x Phusion HF Buffer + 2.0% Proteinase K) was added to each PCR tube and mixed by light vortexing. The samples were then frozen and lysed as in the single worm PCR protocol. 2.5 µl of embryo lysate was used in each 10 µl PCR reaction. Control lysates were generated by mixing 10-15 embryos in the same PCR tube.

### PCR primers

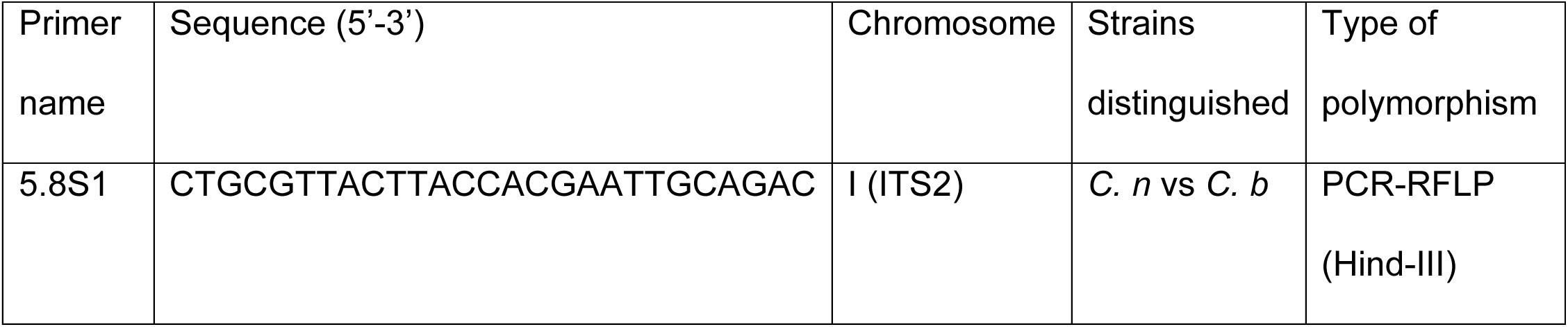
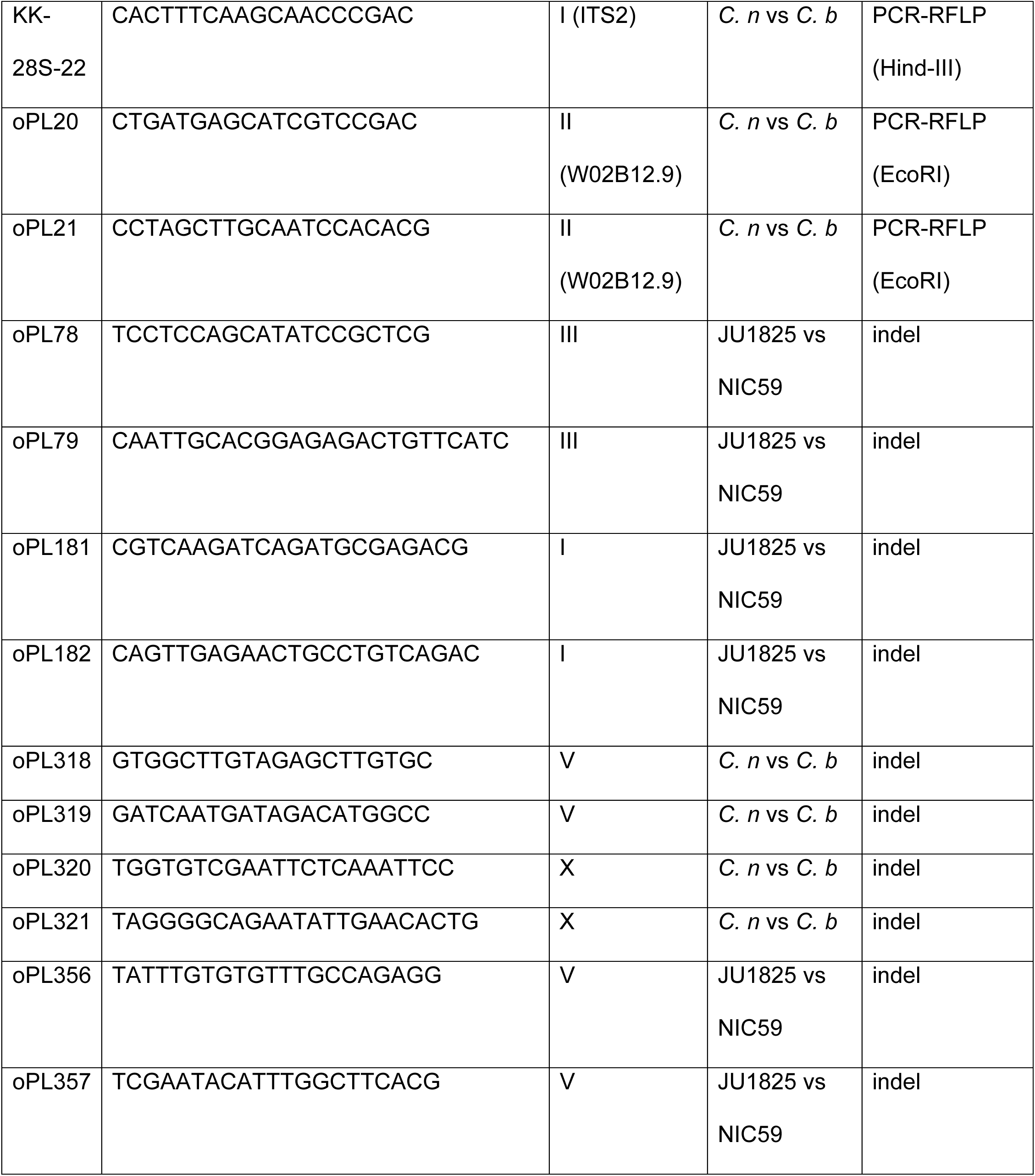

The primers used in the PCR-RFLP polymorphism assays lie in conserved coding sequences that are easily amplified from a range of *Caenorhabditis* species [48]. Primers 5.8S1 and KK-28S-22 were originally used in Kiontke et al [48]. The primers used in the PCR indel polymorphism assays were generated after the assembly of the *C. nouraguensis* genome. Indel polymorphisms between NIC59 and JU1825 were found by manually screening *C. nouraguensis* (JU2079) scaffolds in IGV [56] for regions where JU1825 had read coverage but NIC59 did not (and vice versa). Indel polymorphisms between *C. nouraguensis* and *C. becei* were manually found by aligning homologous sequences on scaffolds assigned to a chromosome of interest.

### Fixing and DAPI staining the female germline

Fixing and DAPI staining germlines was performed as in [57]. F1 L4 females were mated to *C. nouraguensis* L4 males. 24 hours later, we dissected the gonads from fertile F1 females and stained with DAPI. 30-40 females were picked into 15 µl of 1x Egg Buffer + 0.1% Tween-20 on a glass coverslip and their heads were cut off with needles to expel the gonad. After dissection, 15 µl of Fixation Solution (1x Egg Buffer + 7.4% formaldehyde) was added and mixed well by pipetting. Gonads were fixed for 5 minutes, then 15 µl of the mixture was removed and a Histobond slide was laid down gently on top of the coverslip. Excess liquid was removed with Whatman paper and samples were frozen by placing the slide on a metal block on dry ice for 10 minutes. Coverslips were then quickly flipped off with a razor blade and the slides were placed in a Coplin jar with −20° methanol for 1 minute. The slides were then moved to a Coplin jar with 1x PBS + 0.1% Tween-20 for 10 minutes, then a Coplin jar with 1x PBS + 0.1% Tween-20 + 5 µl of 5 mg/ml DAPI for 15 minutes, then a Coplin jar with 1x PBS + 0.1% Tween-20 for 20 minutes. Samples were then mounted with 11 µl Mounting Media (35 µl 2M Tris + 15 µl MilliQ dH_2_O + 450 µl glycerol) and sealed with nail polish. Images were taken using a Nikon Eclipse 80i compound microscope and processed using Image J.

### C. becei genome assembly and linkage map construction

The chromosomal reference assembly for *C. becei* is based on high-coverage paired-end and mate-pair short-read sequencing, low-coverage Pacific Biosciences long reads, and genetic linkage information from an experimental intercross family. We used two inbred lines, QG2082 and QG2083, derived from isofemale lines QG704 and QG711 respectively by sib-mating for 25 generations. QG704 and QG711 were isolated from independent samples of rotten flowers (QG704) or fruit (QG711) collected in Barro Colorado Island, Panama, in 2012 [49]. We generated conservative *de novo* assemblies for the inbred lines, identified SNPs that distinguish them, and then used a six-generation breeding design to generate recombinant populations (G_4_BC_2_) from which we generated a SNP-based linkage map. With the map-scaffolded conservative assembly, we were able to evaluate more contiguous, less conservative genome assemblies for consistency, and we selected one of these as our final *C. becei* genome. Each of these steps is described in detail below.

#### Sequencing

We extracted genomic DNA from QG2083 by proteinase K digestion and isopropanol precipitation [58] and prepared paired-end (PE) 600-bp insert and mate-pair (MP) 4-kb insert Illumina Nextera libraries, from which 100-bp reads were sequenced to approximately 30x and 60x expected coverage (see below) on an Illumina HiSeq 2000 at the NYU Center for Genomics and Systems Biology GenCore facility (New York, USA). PE reads for QG2082 were generated similarly, for 20x expected coverage. PE reads for G_4_BC_2_ mapping populations were generated from 250bp insert libraries sequenced to mean 4x coverage per line with 150-bp reads on a HiSeq 2500. FASTX Toolkit (v. 0.0.13; options ‘-n -l 25 -M 9 -Q 33’; http://hannonlab.cshl.edu/fastx_toolkit/) was used to trim low quality bases from PE reads, and NextClip (v. 1.3; [59]) was used to remove adaptors from MP reads and assess read orientation. Contaminating bacterial sequence was removed with Kraken (v. 1.0; [60]) using a database built from NCBI nt (downloaded 10/06/2014), and sequencing errors were corrected with Blue (v. 1.1.2; [61]). Raw data is available from the NCBI SRA under project PRJNA525787.

#### Genome assembly for map construction

We assembled contigs and conservative scaffolds for QG2083 with the string graph assembler SGA [62]. Scaffolding required contigs to be linked by at least 5 unambiguously mapped mate-pair reads and full path consistency between contigs and all mapped mate-pair reads (option ‘--strict’). PacBio long reads (12x coverage, P4-C2 chemistry, Duke University Center for Genomic and Computational Biology) were used for a single round of gap closing and scaffolding with PBJelly (v. 13.10, minimum evidence of two reads; [63]), followed by scaffold breakage at any regions of mate-pair read discordance with REAPR (v. 1.0.17; [64]). This yielded a fragmented assembly of 9152 scaffolds of length at least 500 bp spanning 95.9 Mb, with half the assembly in scaffolds of 56 kb or more.

#### Variant calling

QG2082 reads were aligned to the QG2083 reference scaffolds with bwa mem (v. 0.5.9; [65]), processed with samtools (v. 1.2; http://www.htslib.org/) and Picard (v. 1.111; http://broadinstitute.github.io/picard/) utilities, and SNPs were called using the GATK HaplotypeCaller (v. 3.1-1; hard filtering on ‘MQ < 40.0 || DP < 4 || FS > 60.0 || ReadPosRankSum < −8.0 || QD < 4.0 || DP > 80’; [66]). Fixed diallelic SNPs were supplemented with calls from reference mapping of the pooled G_4_BC_2_ data and, to mitigate potential mapping bias against highly divergent regions, SNP calls from whole genome alignment of a *de novo* assembly of G_4_BC_2_ data pooled with QG2083 data at 1:1 expected heterozygosity, using the heterozygous aware assembler Platanus [67] and Mummer [68]. This yielded a total of 1.63M diallelic SNPs across 6843 scaffolds spanning 89.2 Mb. These SNPs provide markers for genetic map construction.

#### Genetic map construction

We generated F_2_ animals from reciprocal crosses between QG2083 and QG2082. We then performed 160 single-pair random-mating intercrosses among F_2_s to generate G_3_s, and again among G_3_s to generate G_4_s. These intercrosses allow additional meioses to expand the genetic map. We backcrossed G_4_ females to QG2082 males, generating progeny carrying one recombinant version of the genome and one intact QG2082 genome. To increase the quantity of DNA representing these genomes, we crossed females to QG2082 males again and allowed the resulting G_4_BC_2_ populations to grow for a single generation before DNA extraction and library preparation. Data for 87 G_4_BC_2_ populations with at least 0.5x expected coverage were mapped to the QG2083 scaffolds and genotyped by HMM using a modified MSG pipeline (assuming six crossovers per assembled genome, genotyping error rates of 0.001, and a scaling parameter on transition probabilities of 1×10^-11^, taking the dominant parental assignment as a single genotype for each scaffold) [69]. Markers were filtered in r/qtl based on missing data (<50%) and segregation distortion (χ^2^ test *p* > 1×10^-10^), followed by removal of redundant markers [70]. Based on genotyping error rate estimates we further required a minimum of 70 SNP calls per scaffold and a minimum length of 1 kb, yielding 1337 non-redundant scaffold markers spanning 81.5 Mb. Six linkage groups were recovered over a wide range of minimum pairwise LOD scores, and within linkage group marker ordering was carried out by the minimum spanning tree method implemented in ASMap [71].

#### Assembly of the C. becei draft genome

Linkage group scaffolds were aligned against multiple Platanus *de novo* assemblies using the sequencing data from inbred lines and subsampling of the approximately 350x pooled data from G_4_BC_2_ lines, with the UCSC chain/net pipeline [72]. Genome evolution in the *Caenorhabditis* genus occurs largely through intrachromosomal rearrangement, and 84-94% of the net for each *C. becei* linkage group aligned to the homologous *C. elegans* chromosome. The assembly most concordant with genetic data was selected, based on the number and aligned length of (1) scaffolds mapping to multiple linkage groups and (2) outliers in the ratio of genetic to physical distance. The selected assembly contained six cases where scaffolds mapped to multiple linkage groups – usually terminal regions of large scaffolds, with these terminal alignments spanning just over 1 Mb in total – and three scaffolds where mapping within linkage groups was inconsistent with the genetic data (median absolute deviation 99th percentile for the ratio of genetic to physical span), spanning 179 kb.

The draft assembly improved contiguity and incorporated several Mb of sequence not represented by the genetic map scaffolds, from 2483 redundant marker scaffolds to 256 final scaffolds spanning 84.9 Mb. We estimate a genome size of around 95 Mb based on mapped read-depth [62], with 90 Mb of that unique sequence amenable to assembly and variant calling.

### C. nouraguensis genome assembly

The *C. nouraguensis* reference genome was generated from JU2079, an inbred strain derived from 28 generations of JU1825 sibling matings (Marie-Anne Félix, personal communication). Genomic DNA and RNA were purified from mixed-stage individuals and sent to Mark Blaxter at the University of Edinburgh for Illumina sequencing. 125-bp paired-end reads were generated from two libraries, one with inserts of ∼400 bp (∼55 million read-pairs), the other with inserts of ∼550 bp (∼54 million read-pairs) (SRA project accession PRJEB10884). After generating a preliminary assembly, we used BlobTools [73] for taxonomic classification. Some contigs matched *E. coli* (provided as food) and some matched an unsequenced bacterial species in the Firmicutes phylum. We extracted the Firmicutes contigs and used GSNAP [74] to remove matching read-pairs, and to remove any read-pairs matching the *E. coli* REL606 genome (Genbank accession NC_012967), a strain closely related to OP50. We also removed reads failing Illumina’s ‘chastity filter’ and used cutadapt to trim poor quality sequences (phred score < 10) and adapter sequences from the 3’ ends of reads. We then performed error-correction using Musket [75] with a k-mer size of 28, and performed *de novo* genome assembly, scaffolding and gap closure using Platanus [67] with an initial k-mer size of 21. We then used REAPR [64] to break any misassembled scaffolds. The resulting assembly is approximately 73 Mb in size. We used BUSCO [76] to measure its completeness using a set of conserved genes, and find that our assembly contains 81.5% of conserved single-copy orthologs as a single copy, 9.8% as duplicates, and 3.6% as fragments, very similar to what is seen in the finished *C. elegans* genome assembly [76]. Using MUMMER [68], we ordered and oriented our *C. nouraguensis* scaffolds based on synteny to the *C. becei* genome assembly. Using this approach, some scaffolds remain unplaced and others may be misplaced, either because of true differences between the two species’ genomes, or because MUMMER’s alignments are noisy.

We also generated assemblies of the mitochondrial genomes of *C. becei* strain QG2083, and *C. nouraguensis* strain JU2079 using reads generated by Mark Blaxter’s lab (EMBL ENA accession ERR1018617), filtered as above to remove *E. coli* reads, and down-sampled to 10 million read-pairs for *C. nouraguensis*. We then used the Assembly by Reduced Complexity (ARC) [77] pipeline to assemble mitochondrial genomes, using the *C. elegans* mitochondrial genome sequence as a starting point.

### Whole-genome amplification and library preparation

After testing the fertility of rare viable offspring from crosses of *C. nouraguensis* JU1825/NIC59 females to *C. becei* males (see above), we prepared Illumina whole-genome sequencing libraries as follows. We first transferred each worm individually to a blank NGM plate with no OP50 lawn for 30 minutes to reduce the amount of contaminating bacterial DNA. We then performed whole-genome amplification using the Qiagen REPLI-g Single Cell kit [78]. We picked individual worms into 4 µl of PBS sc and froze them at −20° for at least an hour. We thawed the samples and added 3 µl of Buffer D2, mixed well, and incubated at 65° for 20 minutes rather than the recommended 10 minutes to aid in worm lysis. The rest of the protocol was performed according to the manufacturer’s guidelines. We purified the resulting amplified DNA using the Zymo Research Genomic DNA Clean & Concentrator kit (gDDC-10). We submitted approximately 0.5-0.8 ng of each sample to the Fred Hutchinson Cancer Research Center Genomics Core Facility. A uniquely barcoded sequencing library for each sample was made using the Illumina Nextera library prep kit and sequenced on an Illumina Hiseq2500 to generate 50-bp paired-end reads. We sequenced eleven fertile and ten sterile F1 individuals. We also sequenced a set of control samples so that we could determine how coverage varies across the genome for pure *C. nouraguensis* or *C. becei* populations, as well as to allow us to determine fixed SNPs between the NIC59 and JU1825 *C. nouraguensis* strains. Specifically, we sequenced genomic DNA derived from large populations of NIC59, JU1825 and QG711 (sample names: NIC59_bulk, JU1825_bulk and QG711_bulk). In order to determine whether SNP frequencies and coverage metrics behave as expected in heterozygous or hybrid triploid genomes, we sequenced three additional controls: (a) a single heterozygous NIC59/JU1825 female (sample name: F1_NIC59_JU1825); (b) a sample where we placed one NIC59 female and one JU1825 female in the same tube (sample name: NIC59plusJU1825); and (c) a sample where we placed one NIC59 female, one JU1825 female and one QG711 female in the same tube (sample name: NIC59plusJU1825plusQG711).

### SNP calling, genotyping, and coverage calculations

Using GSNAP [74], we mapped the reads from each sample to a combined reference genome containing the *C. nouraguensis* and *C. becei* nuclear and mitochondrial genome assemblies, the *E. coli* REL606 genome sequence and the Firmicutes contigs described above. We allowed GSNAP to report only a single map position for each read (--npaths=1). We further filtered read mappings, requiring mapq value of at least 20 to select uniquely mapping reads.

After preprocessing bam files by marking duplicates (using picard’s MarkDuplicates, http://broadinstitute.github.io/picard/) and realigning indels (using GATK’s RealignerTargetCreator and IndelRealigner tools [66]), we called SNPs in *C. nouraguensis* scaffolds using samtools mpileup [79] (ignoring indels, disabling the per-base alignment quality option and increasing the depth down-sampling parameter to 6660). We counted reads matching each allele using GATK’s VariantAnnotator tool [66], disabling the down-sampling option and using the “ALLOW_N_CIGAR_READS” option. We noticed that the distribution of allele frequencies in our ‘heterozygous’ control samples was skewed towards the reference allele, likely because those reads are easier to map to the reference assembly. We therefore used this first round of SNP calls to remap all reads to the combined reference genome using GSNAPs “SNP-tolerant” mode that allows reads to map to either haplotype. SNPs were called a second time, after one additional level of read mapping filtering, where we used the R/Bioconductor Rsamtools package [80] to select only reads where the full length of the read could be aligned to the reference genome.

We then used the R/Bioconductor VariantAnnotation package [80] to select high-quality fixed differences between the NIC59 and JU1825 strains as follows: we required a SNP quality score of at least 100; that opposite alleles be fixed (non-reference frequencies of <5% and >95%) in the two strains; that read depth in some *C. nouraguensis* control samples be within typical range for that sample (5-50 for JU1825_bulk and NIC59_bulk, and 100-450 for the combined JU2079 samples); and that read depth be 0 in our *C. becei* control samples (QG711_bulk and the two QG2083 libraries). 337,493 SNPs passed these filters (an average of one SNP every 217 bp). To estimate mean allele frequencies in 50-kb windows across the genome, we counted the total number of reads matching NIC59 and JU1825 alleles across the window and divided the NIC59 count by the total count. We then used a circular binary segmentation algorithm, implemented in the Bioconductor package DNAcopy [81], to estimate the locations of breaks between different haplotypes as well as the average allele frequency in each segment.

Using the same filtered bam files, we determined coverage at each base position using the samtools depth tool [79]. Examining coverage and aligned bases in our control samples (JU1825_bulk, NIC59_bulk and QG711_bulk) reveals very high within-species sequence diversity, as seen in some other nematodes [37,38,82]. This high diversity means that in some genomic regions, especially on the chromosome arms, short reads are unalignable to the reference genomes, resulting in many regions of no coverage. Other genomic regions show more aligned base-pairs, but still harbor many SNPs. Therefore, before calculating mean coverage metrics, we filtered the base positions under consideration to include only those that had coverage of at least 8 reads in the relevant control samples (JU1825_bulk and NIC59_bulk for the *C. nouraguensis* assembly and QG711_bulk for the *C. becei* assembly), and then determined mean values in 50-kb regions of each scaffold. This filtering mitigated diversity-related coverage variation a little, but even in control samples coverage is still lower on chromosome arms than centers. To generate per-chromosome copy number estimates (Fig. 4A), we manually defined 1-2 regions in the center of each chromosome that are well-covered in each of the control samples and took the mean of 50-kb window coverages in each region.

### Fixing and staining embryos

60-80 gravid females were picked into 30 µl of 1x Egg Buffer + 0.1% Tween-20 on a glass coverslip and dissected in half with needles to release embryos. After the females were dissected, we waited for five minutes to allow the mostly 1-cell stage embryos to develop. A Histobond slide was then gently laid on top of the coverslip. Excess liquid was removed with Whatman paper and the samples were frozen by placing the slide on a metal block on dry ice for 10 minutes. Coverslips were then quickly flipped off with a razor blade and the slides were immediately placed in a Coplin jar with −20° methanol for 10 min. The slides were then moved to a Coplin jar with −20° acetone for 10 min. The slides were washed three times for 10 mins each in Coplin jars with 1x PBS + 0.1% Tween-20, then blocked for 30 mins in a Coplin jar with 1x PBS + 0.1% Tween-20 + 0.5% BSA + 0.02% sodium azide (blocking buffer). Each slide was then incubated overnight at 4° with a 1:100 dilution of both mouse monoclonal anti-α-tubulin (clone DM1A, Sigma) and rabbit polyclonal anti-γ-tubulin (ab50721, abcam) primary antibodies in blocking buffer. The next day, the slides were washed three times for 10 mins each in Coplin jars with 1x PBS + 0.1% Tween-20. Each slide was then incubated in the dark for 2 hours at room temperature with a 1:1000 dilution of both Alexa-Fluor 488 goat anti-mouse and Rhodamine Affinipure goat anti-rabbit secondary antibodies (Jackson ImmunoResearch) diluted in 1x PBS + 0.1% Tween-20. The slides were moved to a Coplin jar with 1x PBS + 0.1% Tween-20 for 10 minutes, then a Coplin jar with 1x PBS + 0.1% Tween-20 + 5 µl of 5 mg/ml DAPI for 15 minutes, then a Coplin jar with 1x PBS + 0.1% Tween-20 for 20 minutes. Samples were mounted with 12 µl VectaShield (H-1000, Vector Laboratories) and sealed with nail polish. Samples were imaged using a Nikon Eclipse 80i compound microscope with a 100x oil objective (1.45 NA). Images were processed using Image J.

We only analyzed embryos at early stages of embryogenesis (1-4 cell stage). In normal embryos at this stage, the DNA in the two polar bodies appears as two distinct clumps at the periphery of the embryo that are easily distinguishable from other embryonic nuclei. Polar bodies in hybrid embryos were considered to have abnormal morphologies if they were unlike those observed in the JU1825 control. This encompasses a range of abnormalities such as not being round, being larger, or having easily distinguishable chromatids.

### Male UV irradiation

Virgin *C. nouraguensis* L4 males and L4 females were collected and separately placed onto two new NGM plates seeded with OP50 and rimmed with palmitic acid. We added a few females to the male plate to coax the males to stay on the surface of the plate. The next day, now young adult males were picked onto a blank NGM plate rimmed with palmitic acid. This plate was placed in a CL-1000 UV Crosslinker (Ultra-Violet Products) without a lid and the males were exposed to 70,000 µJ/cm^2^ 254-nm UV radiation. Immediately afterward, the males were picked onto new NGM plates seeded with OP50 and rimmed with palmitic acid. Each plate had 30-90 irradiated males and 30-90 young adult virgin females. The next day, rare viable larvae were picked from the cross plates onto a new NGM plate seeded with OP50 and rimmed with palmitic acid. When these progeny reached adulthood, they were individually frozen, lysed and PCR genotyped.

We tested the following UV exposures: 15,000, 40,000, 50,000, 60,000, 70,000, 80,000, 150,000 and 250,000 µJ/cm^2^. Males treated with lower exposures (15,000 and 40,000 µJ/cm^2^) were able to produce many viable progeny, indicating that their sperm DNA was not damaged enough to induce paternal genome loss. Males treated with the highest exposures (150,000 and 250,000 µJ/cm^2^) mostly stopped moving and never recovered. We found that males treated with 70,000 µJ/cm^2^ were healthy enough to mate and produce many dead fertilized embryos, but relatively few viable progeny.

Each 30 female x 30 male or 90 female x 90 male plate eventually became overcrowded with dead embryos, making it difficult to manually quantify their exact number. Instead of counting each embryo on these plates, we performed a five female x five male cross in parallel and counted their dead embryos. We then determined the average number of dead embryos per female on the 5 female x 5 male plates and multiplied that by the number of total females to get a rough estimate of the total number of dead embryos.

## ACKNOWLEDGMENTS

We thank Marie-Anne Félix and Christian Braendle for providing the strains of *C. nouraguensis* used in this study; Max Bernstein, Vicky Cattani, Taniya Kaur, Jasmine Nicodemus and Annalise Paaby for help performing *C. becei* mapping crosses and preparing sequencing libraries; Mark Blaxter, Lewis Stevens and members of the *Caenorhabditis* Genomes Project for sequencing *C. nouraguensis* (JU2079) as well as providing a public BLAST server of newly sequenced *Caenorhabditis* species (http://blast.caenorhabditis.org/); Maulik Patel for help in preparing DNA for *C. nouraguensis* genome sequencing (JU1825 and NIC59); Lews Caro and Sam Hart for additional experiments; Jihong Bai for sharing his UV-irradiator; Lamia Wahba for advice on single-worm sequencing; Needhi Bhalla and Mara Schvarzstein for discussions of *C. elegans* female meiosis; Jim Priess for discussions of *C. elegans* development; and Hannah Seidel for advice on time-lapse imaging of *Caenorhabditis* embryos. We also thank Barbara Wakimoto, Diane Shakes, Lisa Kursel, Irini Topalidou, Ofer Rog and members of his lab, and members of the Ailion lab for comments on the manuscript. We thank the Fred Hutchinson Genomics core for Illumina library preparation and sequencing.

## SUPPORTING INFORMATION LEGENDS

**Fig S1. Crosses measuring intra-strain viability and intraspecies and interspecies fecundity. (A)** Wild isolates of *C. becei* (QG704 and QG711) and *C. nouraguensis* (NIC59 and JU1825) have high levels of intra-strain viability. All strains have a sex ratio skewed towards females, some of which show a statistically significant difference from a 50:50 sex ratio (Fisher’s exact test with Bonferroni correction, JU1825 p=1.0, NIC59 p=0.06, QG711 p=0.03, QG704 p=0.03). The total number of offspring quantified for each cross is shown to the right of each bar graph. Data from both graphs are derived from the same crosses. **(B)** A graph showing the number of embryos laid for intraspecies *C. nouraguensis* crosses (10 NIC59 females x 10 NIC59 males) and interspecies *C. nouraguensis* female x *C. becei* male crosses (10 NIC59 females x 10 QG711 males) in a one-hour window on each of the first three days after the crosses were set. There are three replicates for each type of cross. Each point represents the number of embryos laid for a replicate in a one-hour window that day and the bar graph shows the average of those replicates. The *C. nouraguensis* female x *C. becei* male interspecies hybridization had significantly less embryos on days 2 and 3 of egg-laying as compared to the intraspecies *C. nouraguensis* crosses (*, day 2 p=0.04, day 3 p=0.04, Kruskal-Wallis test).

**Fig S2. Summary of interspecies crosses.** Rows show the females of each cross and males are shown in columns. The wild isolate strains used for each species are indicated. Black boxes are intraspecies crosses. Grey boxes are untested interspecies hybridizations. Rare viable F1 adults are present only when crossing *C. nouraguensis* females to *C. becei* males. Rare viable but sick F1 larvae are present in both directions of *C. becei* x *C. yunquensis* crosses. Worms mate but do not produce F1 embryos in *C. panamensis* female x *C. nouraguensis* male and *C. becei* female x *C. panamensis* male crosses. At least 12,000 dead F1 were screened for each cross that gave embryos.

**Fig S3. Fertile progeny with a maternal genotype and sterile progeny with a hybrid genotype are produced when *C. nouraguensis* females are crossed to males of a different *C. becei strain*.** (A) We tested the fertility of rare viable F1 derived from *C. nouraguensis* JU1825 x *C. becei* QG704 crosses by backcrossing to JU1825 individuals of the opposite sex. F1 were then genotyped at the ITS2 locus using a PCR-RFLP assay. **(B)** A gel showing the sex, fertility and genotype at the ITS2 locus of viable adult F1. All fertile F1 have a maternal genotype, with one exception: one hybrid F1 female laid inviable F2 embryos (F*, still considered fertile). All sterile F1 had a hybrid genotype. **(C)** A table summarizing the genotyping data in (B).

**Fig S4. Fertile F1 females are diploid.** The −1 oocytes from *C. nouraguensis* JU1825 and *C. becei* females primarily have six DAPI-staining bodies (two examples shown here). Most fertile F1 females derived from JU1825 female x QG711 male crosses have six DAPI-staining bodies (Example fertile F1 female #1). A minority have eight DAPI-staining bodies. In three of these cases, there appear to be seven relatively normal sized DAPI bodies plus a very small one (Example fertile F1 female #2, small DAPI body highlighted by white arrowhead). In the other two cases, all eight DAPI bodies appear roughly equal in size (Example fertile F1 female #3). This higher number of DAPI-staining bodies is not the chance observation of a low frequency meiotic defect in a nucleus that happens to be in the −1 oocyte position (for example, homologs fail to recombine and increase the number of univalents) because we observed the same number of DAPI-staining bodies in both germlines of the same fertile F1 female. We hypothesize that these extra DAPI-staining bodies represent extra DNA (either maternal or paternal) in addition to the two chromatids inherited from each maternal bivalent. Scale bar: 5 µm.

**Fig S5. Fertile F1 inherit two randomly selected homologous chromatids from each maternal bivalent. (A)** The six possible ways of combining two of the four genetically distinct chromosomes in a bivalent are illustrated, with their expected genotypic signature below them. There are five distinct genotypic signatures. Two result from combining sister-chromatids and are called “Sisters_1” and “Sisters_2”. Three result from combining homologous chromosomes and are called “Homologs_1”, “Homologs_2” and “Homologs_3”. (**B)** A table summarizing the genotype of each maternal chromosome for each sequenced F1 individual. The F1’s sex and fertility are noted. Each individual’s ploidy is inferred from the genotyping data. The genotype “Homologs_ambiguous” refers to chromosomes that are heterozygous in their centers, but one end of the chromosome is not obviously heterozygous (N/J) or homozygous for either NIC59 or JU1825. The genotype “Hemizygous X” refers to hemizygous X-chromosomes in males that have half the read depth of the autosomes. The genotype “Triploid” refers to chromosomes that have three copies instead of two based on relative read depth and whole-chromosome genotype. The “Sisters_3” genotype refers to chromosomes that have inherited two non-recombinant JU1825 chromatids. Three fertile females (F1_8, F1_11 and F1_39) have NIC59 and JU1825 alleles in the center of their chromosomes, but exhibit a slight skew from the expected 0.50 NIC59 allele frequency (Fig S7). We hypothesize that this skew is due to contaminating DNA derived from backcrossing each female when testing her fertility. For example, a viable female that was backcrossed to a JU1825 male would carry JU1825 sperm and therefore JU1825 DNA in her spermatheca. In this case, the contaminating DNA would skew the female’s entire genome to a lower NIC59 SNP frequency. Consistent with this, in all three females the more abundant allele matches the genotype of the male she was backcrossed to (Table S1). Correcting for this potential backcrossing contamination, the genotypes of all the chromosomes in these three females are consistent with the inheritance of two homologous chromatids. One fertile female (F1_25) and two sterile females (F1_18 and F1_26) have a very low proportion of NIC59 reads across their entire genome (Fig S7). We hypothesize this skew is due to contaminating DNA from JU1825 males during fertility testing and poor lysis of the female (Table S1). The low proportion of NIC59 reads made it difficult to infer actual maternal chromosome genotypes and are therefore categorized as “ambiguous”. **(C)** Tables summarizing the frequency of genotypes per chromosome in all fertile F1, all sterile F1, or a combination of all fertile and sterile F1.

**Fig S6. Genotypic signatures of multiple potential mechanisms of diploid maternal inheritance. (A)** A schematic of the cross used to determine how diploidy is restored in gynogenetically produced offspring in the interspecies cross. Heterozygous (NIC59/JU1825) *C. nouraguensis* females are crossed to *C. becei* (QG711) males. Individual viable offspring resulting from the hybridization undergo whole genome sequencing. **(B)** If diploid maternal inheritance results from mitotic (apomixis) rather than meiotic divisions, then viable F1 are clones of their mother and will be heterozygous NIC59/JU1825 across their entire genome (0.5 NIC59 allele frequency). **(C)** Diploid maternal inheritance can occur through endomitosis, which is when a haploid maternal genome replicates without a cell division. This results in two exact copies of each chromosome and therefore homozygosity for either NIC59 (1.0 NIC59 allele frequency) or JU1825 alleles (0.0 NIC59 allele frequency) across the entire genome. **(D)** Diploid maternal inheritance can occur through automixis (combining two of the four meiotic products). Combining two homologous chromatids will result in heterozygosity in the center of a chromosome, and heterozygosity or homozygosity at the chromosome ends. Combining two sister chromatids will result in homozygosity in the center of a chromosome and heterozygosity at one of the ends. Inheriting two sister chromatids for one chromosome and two homologous chromatids for another chromosome in the same genome is theoretically possible, but not depicted here.

**Fig S7. Fertile F1 inherit two randomly selected homologous chromatids from each maternal bivalent.** Each of the following pages contains plots describing whole-genome sequencing data of either a rare viable F1 individual or a control DNA sample. The sample name is at the top of each page, along with the individual’s sex, fertility and strain it was backcrossed to for fertility testing (if applicable). Each page has five rows of plots. The first row shows the genotypes of the sample’s *C. nouraguensis* maternal chromosomes (i.e. average NIC59 allele frequency in 50-kb windows across the *C. nouraguensis* assembly). Haplotype change points and average allele frequency for each segment are shown by the green horizontal lines. The second and third rows show the sample’s average read coverage of the *C. becei* and *C. nouraguensis* assemblies in 50-kb windows. The fourth and fifth rows show the average GC content of the *C. becei* and *C. nouraguensis* assemblies in 50-kb windows. The gray vertical lines represent breaks between scaffolds.

**Fig S8. Fertilization with UV-irradiated sperm does not cause frequent diploid maternal inheritance in intraspecies crosses. (A)** Fertilization with UV-irradiated sperm does not significantly increase diploid maternal inheritance in *C. nouraguensis* intraspecies crosses. Heterozygous *C. nouraguensis* (NIC59/JU1825) females were mated to UV-irradiated *C. nouraguensis* (NIC59) males. Dead embryos produced by this cross were PCR genotyped at a single autosomal locus (oPL78+79). The gel shows the genotypes of a fraction of the dead embryos assayed. Most embryos had either a JU1825 (11/31) or NIC59 genotype (17/31), consistent with haploid maternal inheritance and destruction of the paternal NIC59 genome. Although 3/31 embryos had a heterozygous genotype (red stars), we found that UV-irradiation can fail to completely destroy the paternal genome (see Fig S8B), suggesting that these three heterozygotes may result from haploid maternal inheritance of a JU1825 allele and haploid paternal inheritance of a NIC59 allele due to incomplete destruction of paternal DNA. **(B)** The paternal genome is not always eliminated by UV irradiation. *C. nouraguensis* (JU1825) females were mated to UV-irradiated *C. nouraguensis* (NIC59) males. Dead embryos produced by this cross were PCR genotyped at a single autosomal locus (oPL78+79). The gel shows the genotypes of a fraction of the dead embryos assayed. Red stars indicate cases in which the UV-irradiated paternal NIC59 DNA can be detected.

**Table S1. Characteristics of the samples submitted for whole-genome sequencing.** Whole-genome amplified F1 individuals derived from the interspecies cross are labeled as “F1_” followed by a barcode number. Sample “F1_NIC59_JU1825” is one heterozygous NIC59/JU1825 female that was whole-genome amplified and serves as a control for genome-wide heterozygosity. Sample “NIC59plusJU1825” comprises one NIC59 and one JU1825 female placed in the same tube and whole-genome amplified. This also serves as a control for genome-wide heterozygosity. Sample “NIC59plusJU1825plusQG711” comprises one NIC59, one JU1825 and one QG711 female placed into the same tube and whole-genome amplified. This serves as a triploid (diploid *C. nouraguensis* and haploid *C. becei*) control. The “NIC59_bulk”, “JU1825_bulk” and “QG711_bulk” samples are genomic preps made from large populations of each strain containing mixed developmental stages and sexes: these libraries were prepared without whole-genome amplification. The reads from the “NIC59_bulk” and “JU1825_bulk” samples were used to identify fixed SNPs between the two strains. The second column shows the interspecies cross that produced each viable F1 animal. In each case, heterozygous N/J *C. nouraguensis* females were crossed to *C. becei* QG711 males. However, the heterozygous *C. nouraguensis* females could have either a NIC59 or JU1825 mitochondrial genotype (denoted in parentheses). A fraction of each sample’s reads (Illumina 50-bp paired end) derive from *E.coli* (their food). The *E. coli* fraction is quite high in some samples, likely due to inefficient cleaning of single worms (see Materials and Methods). Of the reads that map to *Caenorhabditis* nuclear genomes, the approximate coverage and percent of all worm reads that map to each *Caenorhabditis* nuclear genome are shown.

**Video S1. *C. nouraguensis* (JU1825) embryonic development.** DIC time-lapse imaging of embryos derived from JU1825 female x JU1825 male crosses. Most develop into larvae, but two embryos fail to develop. Dissection and mounting of embryos causes some embryos to arrest at approximately the one-cell stage.

**Video S2. *C. becei* (QG711) embryonic development.** DIC time-lapse imaging of embryos derived from QG711 female x QG711 male crosses. Most develop into larvae, but one embryo fails to develop. Dissection and mounting of embryos causes some embryos to arrest at approximately the one-cell stage.

**Video S3. *C. becei* female x *C. nouraguensis* male embryonic death.** DIC time-lapse imaging of embryos derived from *C. becei* QG711 female x *C. nouraguensis* JU1825 male crosses. These embryos undergo cell divisions until approximately the 32-cell stage at which point they stereotypically arrest and fail to undergo further divisions. The embryos come out of focus at the end of the video.

**Video S4. *C. nouraguensis* female x *C. becei* male embryonic death.** DIC time-lapse imaging of embryos derived from *C. nouraguensis* JU1825 female x *C. becei* QG711 male crosses. These embryos arrest either at early or later stages of development. The early-arresting embryos generally have abnormal cell membrane morphologies. Specifically, some arrested embryos have faint cell membranes that appear to distend and take up more space than the normally more rounded cells seen in the early embryos of either parental species. Also, some early-arresting embryos have many nuclei but no discernible cell membranes separating them. The embryos that arrest at later stages have superficially normal cell and nuclear morphologies and undergo cell divisions until approximately the 44-87 cell stage. After this point, the cells in these embryos largely halt their divisions, with many appearing to be dissociated from one another, rounded, and of variable size. One common feature of the early and late-arresting embryos is that both can have multinucleate cells.

