## Supplementary information for "Hybridization promotes asexual reproduction in *Caenorhabditis* nematodes"

### Figure S1

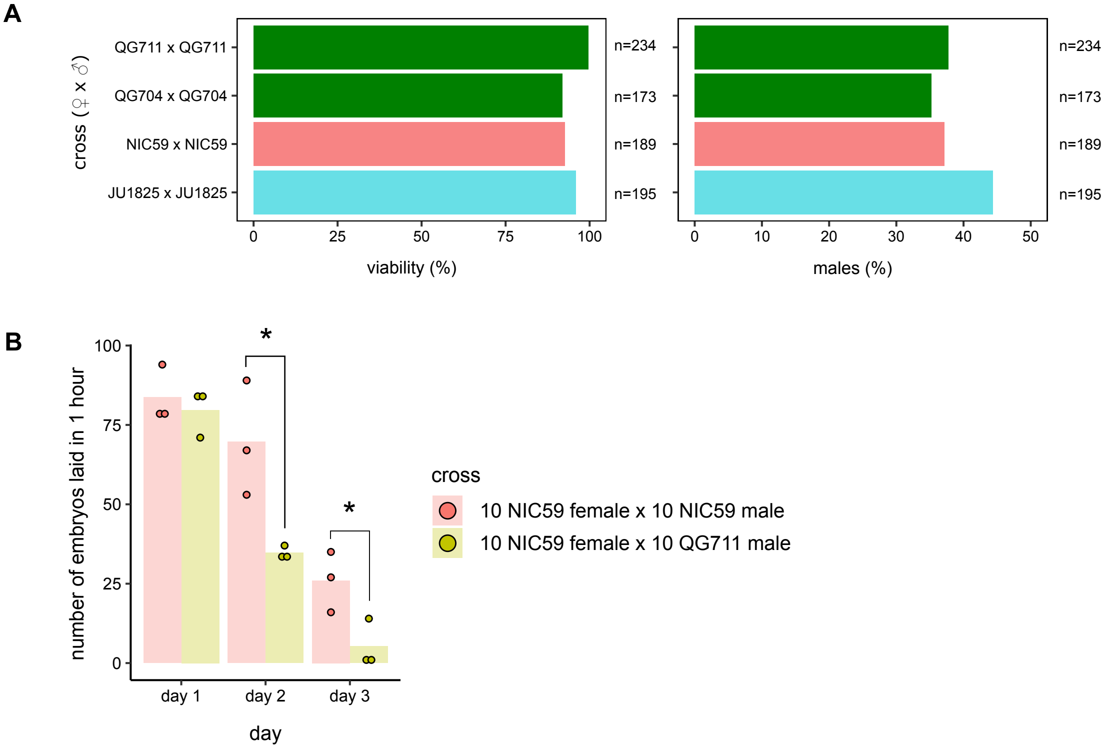


**Figure S1. Crosses measuring intra-strain viability and intraspecies and interspecies fecundity. (A)** Wild isolates of *C. becei* (QG704 and QG711) and *C. nouraguensis* (NIC59 and JU1825) have high levels of intra-strain viability. All strains have a sex ratio skewed towards females, some of which show a statistically significant difference from a 50:50 sex ratio (Fisher's exact test with Bonferroni correction, JU1825 p=1.0, NIC59 p=0.06, QG711 p=0.03, QG704 p=0.03). The total number of offspring quantified for each cross is shown to the right of each bar graph. Data from both graphs are derived from the same crosses. **(B)** A graph showing the number of embryos laid for intraspecies *C. nouraguensis* crosses (10 NIC59 females x 10 NIC59 males) and interspecies *C. nouraguensis* female x *C. becei* male crosses (10 NIC59 females x 10 QG711 males) in a one-hour window on each of the first three days after the crosses were set. There are three replicates for each type of cross. Each point represents the number of embryos laid for a replicate in a one-hour window that day and the bar graph shows the average of those replicates. The *C. nouraguensis* female x *C. becei* male interspecies hybridization had significantly less embryos on days 2 and 3 of egg-laying as compared to the intraspecies *C. nouraguensis* crosses (*, day 2 p=0.04, day 3 p=0.04, Kruskal-Wallis test).

**Figure S2**

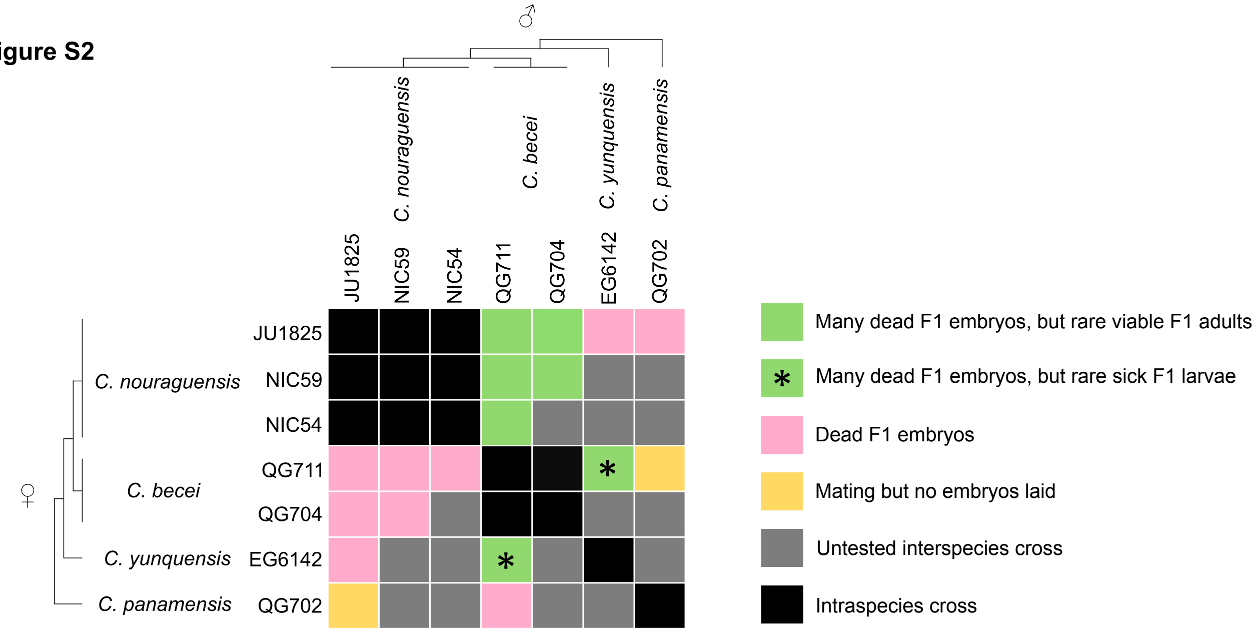


**Figure S2. Summary of interspecies crosses.** Rows show the females of each cross while males are shown in columns. The wild isolate strains used for each species are indicated. Black boxes are intraspecies crosses. Grey boxes are untested interspecies hybridizations. Rare viable F1 adults are present only when crossing *C. nouraguensis* females to *C. becei* males. Rare viable but sick F1 larvae are present in both directions of *C. becei* x *C. yunquensis* crosses. Worms mate but do not produce F1 embryos in *C. panamensis* female x *C. nouraguensis* male and *C. becei* female x *C. panamensis* male crosses. At least 12,000 dead F1 were screened for each cross that gave embryos.

### Figure S3

A

*C. nouraguensis* (JU1825) ♀ x *C. becei* (QG704) ♂

↓

Rare viable F1

1) Backcross F1 to JU1825 to test fertility (F= fertile, F*=fertile but dead F2 progeny, S=sterile).
2) PCR genotype F1 at ITS2 locus (Primers= 5.8S-1 + 28S-22, HindIII-HF digest).

B

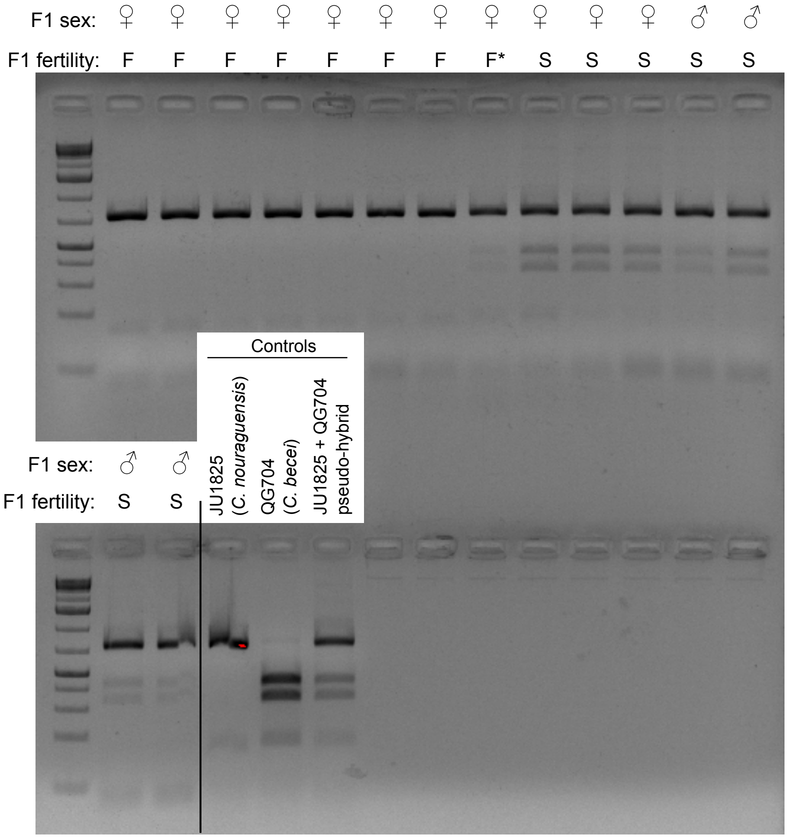


C

| Fertility of F1 adult | Hybrid (*C. nouraguensis*/*C. becei* ) | | Maternal (*C. nouraguensis*) | | |
|---|---|---|---|---|---|
| | female | male | female | male | total |
| Fertile | 1 | 0 | 7 | 0 | 8 |
| Sterile | 3 | 4 | 0 | 0 | 7 |
| total | 4 | 4 | 7 | 0 | 15 |

**Figure S3. Fertile progeny with a maternal genotype and sterile progeny with a hybrid genotype are produced when *C. nouraguensis* females are crossed to males of a different *C. becei* strain. (A)** We tested the fertility of rare viable F1 derived from *C. nouraguensis* JU1825 x *C. becei* QG704 crosses by backcrossing to JU1825 individuals of the opposite sex. F1 were then genotyped at the ITS2 locus using a PCR-RFLP assay. **(B)** A gel showing the sex, fertility and genotype at the ITS2 locus of viable adult F1. All fertile F1 have a maternal genotype, with one exception: one hybrid F1 female laid inviable F2 embryos (F*, still considered fertile). All sterile F1 had a hybrid genotype. **(C)** A table summarizing the genotyping data in (B).

**Figure S4**

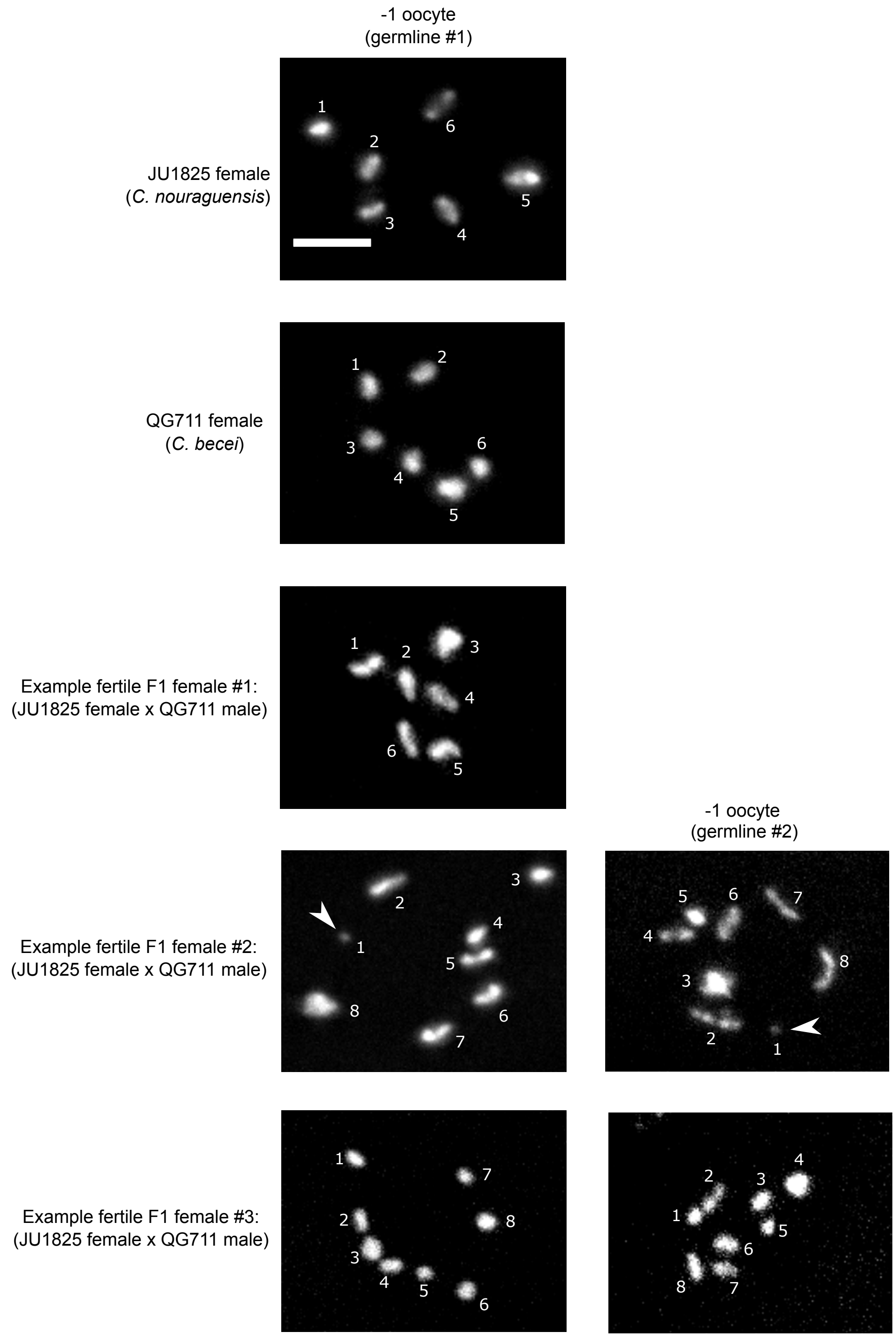


**Figure S4. Fertile F1 females are diploid.** The -1 oocytes from *C. nouraguensis* (JU1825) and *C. becei* (QG711) females primarily have six DAPI-staining bodies (two examples shown here). Most fertile F1 females derived from JU1825 female x QG711 male crosses have six DAPI-staining bodies (Example fertile F1 female #1). A minority have eight DAPI-staining bodies. In three of these cases, there appear to be seven relatively normal sized DAPI bodies plus a very small one (Example fertile F1 female #2, small DAPI body highlighted by white arrowhead). In the other two cases, all eight DAPI bodies appear roughly equal in size (Example fertile F1 female #3). This higher number of DAPI-staining bodies is not the chance observation of a low frequency meiotic defect in a nucleus that happens to be in the -1 oocyte position (for example, homologs fail to recombine and increase the number of univalents) because we observed the same number of DAPI-staining bodies in both germlines of the same fertile F1 female. We hypothesize that these extra DAPI-staining bodies represent extra DNA (either maternal or paternal) in addition to the two chromatids inherited from each maternal bivalent. Scale bar: 5 µm.

### Figure S5

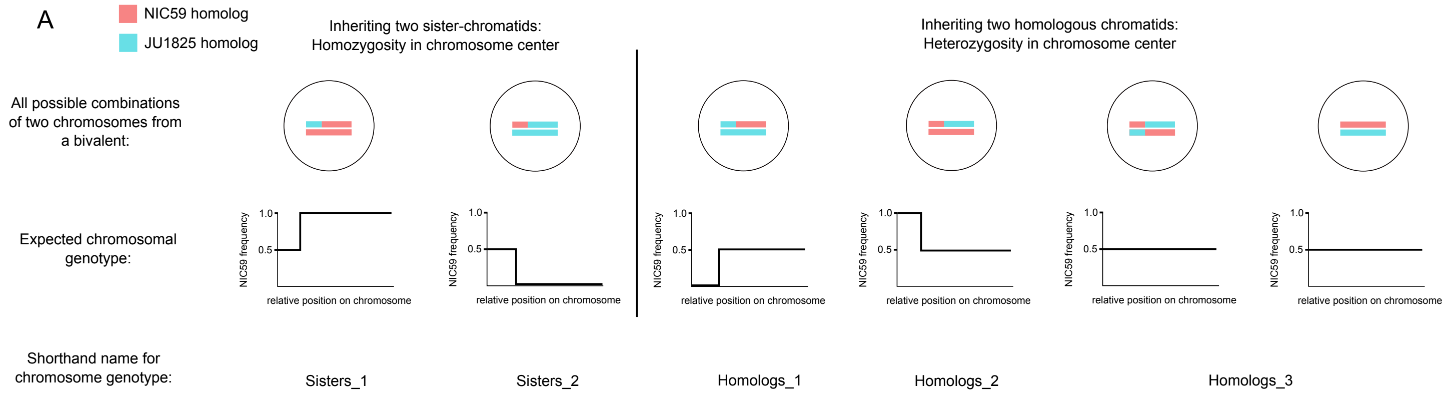


B

*C. nouraguensis* chromosome genotypes

| F1 sample name | fertility | sex | inferred ploidy | Chr. I | Chr. II | Chr. III | Chr. IV | Chr. V | Chr. X |
|---|---|---|---|---|---|---|---|---|---|
| F1_1 | fertile | female | diploid | Homologs_3 | Homologs_3 | Homologs_3 | Homologs_2 | Homologs_2 | Homologs_2 |
| F1_5 | fertile | female | diploid | Homologs_2 | Homologs_3 | Homologs_3 | Homologs_1 | Homologs_2 | Homologs_1 |
| F1_29 | fertile | female | diploid | Homologs_3 | Homologs_ambiguous | Homologs_3 | Homologs_1 | Homologs_3 | Homologs_3 |
| F1_41 | fertile | female | diploid | Homologs_2 | Homologs_3 | Homologs_3 | Homologs_3 | Homologs_3 | Homologs_3 |
| F1_8 | fertile | female | likely diploid, backcross contamination | Homologs_3 | Homologs_3 | Homologs_3 | Homologs_3 | Homologs_3 | Homologs_2 |
| F1_11 | fertile | female | likely diploid, backcross contamination | Homologs_3 | Homologs_3 | Homologs_3 | Homologs_1 | Homologs_1 | Homologs_3 |
| F1_39 | fertile | female | likely diploid, backcross contamination | Homologs_3 | Homologs_1 | Homologs_3 | Homologs_1 | Homologs_3 | Homologs_1 |
| F1_25 | fertile | female | ambiguous, backcross contamination | ambiguous | ambiguous | ambiguous | ambiguous | ambiguous | ambiguous |
| F1_46 | fertile | male | diploid | Homologs_ambiguous | Homologs_3 | Homologs_2 | Homologs_1 | Homologs_3 | Hemizygous X |
| F1_4 | fertile | male | diploid | Homologs_2 | Homologs_3 | Homologs_3 | Triploid | Homologs_3 | Hemizygous X |
| F1_48 | fertile | male | diploid | Homologs_2 | Homologs_3 | Triploid | Homologs_1 | Homologs_3 | Hemizygous X |
| F1_26 | sterile | female | ambiguous, backcross contamination | ambiguous | ambiguous | ambiguous | ambiguous | ambiguous | ambiguous |
| F1_18 | sterile | female | ambiguous, backcross contamination | ambiguous | ambiguous | ambiguous | ambiguous | ambiguous | ambiguous |
| F1_6 | sterile | female | triploid hybrid | Homologs_2 | Homologs_3 | Homologs_3 | Homologs_3 | Homologs_1 | Homologs_1 |
| F1_12 | sterile | female | triploid hybrid | Homologs_1 | Homologs_ambiguous | Homologs_3 | Homologs_3 | Homologs_3 | Homologs_3 |
| F1_20 | sterile | female | triploid hybrid | Homologs_2 | Homologs_2 | Homologs_3 | Homologs_3 | Homologs_3 | Homologs_1 |
| F1_16 | sterile | male | diploid-triploid hybrid | Homologs_1 | Homologs_1 | Homologs_1 | Homologs_3 | Homologs_3 | Hemizygous X |
| F1_23 | sterile | male | diploid-triploid hybrid | Homologs_3 | Homologs_2 | Homologs_1 | Homologs_1 | Sisters_3 | Hemizygous X |
| F1_10 | sterile | male | diploid-triploid hybrid | Homologs_2 | Homologs_2 | Homologs_3 | Triploid | Homologs_3 | Homologs_2 |
| F1_17 | sterile | male | triploid hybrid | Homologs_3 | Homologs_3 | Sisters_1 | Homologs_2 | Homologs_2 | Homologs_2 |
| F1_21 | sterile | male | triploid hybrid | Homologs_3 | Homologs_1 | Homologs_1 | Homologs_2 | Homologs_2 | Homologs_2 |

C

Fertile F1

| Chromosome genotype | Chr. I | Chr. II | Chr. III | Chr. IV | Chr. V | Chr. X | total |
|---|---|---|---|---|---|---|---|
| Homologs_1 | 0 | 1 | 0 | 6 | 1 | 2 | 10 |
| Homologs_2 | 4 | 0 | 1 | 1 | 2 | 2 | 10 |
| Homologs_3 | 5 | 8 | 8 | 2 | 7 | 3 | 33 |
| Homologs_ambiguous | 1 | 1 | 0 | 0 | 0 | 0 | 2 |
| Sisters_1 | 0 | 0 | 0 | 0 | 0 | 0 | 0 |
| Sisters_2 | 0 | 0 | 0 | 0 | 0 | 0 | 0 |
| Sisters_3 | 0 | 0 | 0 | 0 | 0 | 0 | 0 |
| Hemizygous X | 0 | 0 | 0 | 0 | 0 | 3 | 3 |
| Triploid | 0 | 0 | 1 | 1 | 0 | 0 | 2 |
| ambiguous | 1 | 1 | 1 | 1 | 1 | 1 | 6 |
| total | 11 | 11 | 11 | 11 | 11 | 11 | 66 |

Sterile F1

| Chromosome genotype | Chr. I | Chr. II | Chr. III | Chr. IV | Chr. V | Chr. X | total |
|---|---|---|---|---|---|---|---|
| Homologs_1 | 2 | 2 | 3 | 1 | 1 | 2 | 11 |
| Homologs_2 | 3 | 3 | 0 | 2 | 2 | 3 | 13 |
| Homologs_3 | 3 | 2 | 4 | 4 | 4 | 1 | 18 |
| Homologs_ambiguous | 0 | 1 | 0 | 0 | 0 | 0 | 1 |
| Sisters_1 | 0 | 0 | 1 | 0 | 0 | 0 | 1 |
| Sisters_2 | 0 | 0 | 0 | 0 | 0 | 0 | 0 |
| Sisters_3 | 0 | 0 | 0 | 0 | 1 | 0 | 1 |
| Hemizygous X | 0 | 0 | 0 | 0 | 0 | 2 | 2 |
| Triploid | 0 | 0 | 0 | 1 | 0 | 0 | 1 |
| ambiguous | 2 | 2 | 2 | 2 | 2 | 2 | 12 |
| total | 10 | 10 | 10 | 10 | 10 | 10 | 60 |

Fertile and Sterile F1

| Chromosome genotype | Chr. I | Chr. II | Chr. III | Chr. IV | Chr. V | Chr. X | total |
|---|---|---|---|---|---|---|---|
| Homologs_1 | 2 | 3 | 3 | 7 | 2 | 4 | 21 |
| Homologs_2 | 7 | 3 | 1 | 3 | 4 | 5 | 23 |
| Homologs_3 | 8 | 10 | 12 | 6 | 11 | 4 | 51 |
| Homologs_ambiguous | 1 | 2 | 0 | 0 | 0 | 0 | 3 |
| Sisters_1 | 0 | 0 | 1 | 0 | 0 | 0 | 1 |
| Sisters_2 | 0 | 0 | 0 | 0 | 0 | 0 | 0 |
| Sisters_3 | 0 | 0 | 0 | 0 | 1 | 0 | 1 |
| Hemizygous X | 0 | 0 | 0 | 0 | 0 | 5 | 5 |
| Triploid | 0 | 0 | 1 | 2 | 0 | 0 | 3 |
| ambiguous | 3 | 3 | 3 | 3 | 3 | 3 | 18 |
| total | 21 | 21 | 21 | 21 | 21 | 21 | 126 |

**Figure S5. Fertile F1 inherit two randomly selected homologous chromatids from each maternal bivalent. (A)** The six possible ways of combining two of the four genetically distinct chromosomes in a bivalent are illustrated, with their expected genotypic signature below them. There are five distinct genotypic signatures. Two result from combining sister-chromatids and are called "Sisters_1" and "Sisters_2". Three result from combining homologous chromosomes and are called "Homologs_1", "Homologs_2" and "Homologs_3". **(B)** A table summarizing the genotype of each maternal chromosome for each sequenced F1 individual. The F1's sex and fertility are noted. Each individual's ploidy is inferred from the genotyping data. The genotype "Homologs_ambiguous" refers to chromosomes that are heterozygous in their centers, but one end of the chromosome is not obviously heterozygous (N/J) or homozygous for either NIC59 or JU1825. The genotype "Hemizygous X" refers to hemizygous X-chromosomes in males that have half the read depth of the autosomes. The genotype "Triploid" refers to chromosomes that have three copies instead of two based on relative read depth and whole chromosome genotype. The "Sisters_3" genotype refers to chromosomes that have inherited two non-recombinant JU1825 chromatids. Three fertile females (F1_8, F1_11 and F1_39) have NIC59 and JU1825 alleles in the center of their chromosomes, but exhibit a slight skew from the expected 0.50 NIC59 allele frequency (Figure S7). We hypothesize that this skew is due to contaminating DNA derived from backcrossing each female when testing her fertility. For example, a viable female that was backcrossed to a JU1825 male would carry JU1825 sperm and therefore JU1825 DNA in her spermatheca. In this case, the contaminating DNA would skew the female's entire genome to a lower NIC59 SNP frequency. Consistent with this, in all three females the more abundant allele matches the genotype of the male she was backcrossed to (Table S1). Correcting for this potential backcrossing contamination, the genotypes of all the chromosomes in these three females are consistent with the inheritance of two homologous chromatids. One fertile female (F1_25) and two sterile females (F1_18 and F1_26) have a very low proportion of NIC59 reads across their entire genome (Figure S7). We hypothesize this skew is due to contaminating DNA from JU1825 males during fertility testing and poor lysis of the female (Table S1). The low proportion of NIC59 reads made it difficult to infer actual maternal chromosome genotypes and are therefore categorized as "ambiguous". **(C)** Tables summarizing the frequency of genotypes per chromosome in all fertile F1, all sterile F1, or a combination of all fertile and sterile F1.

### Figure S6

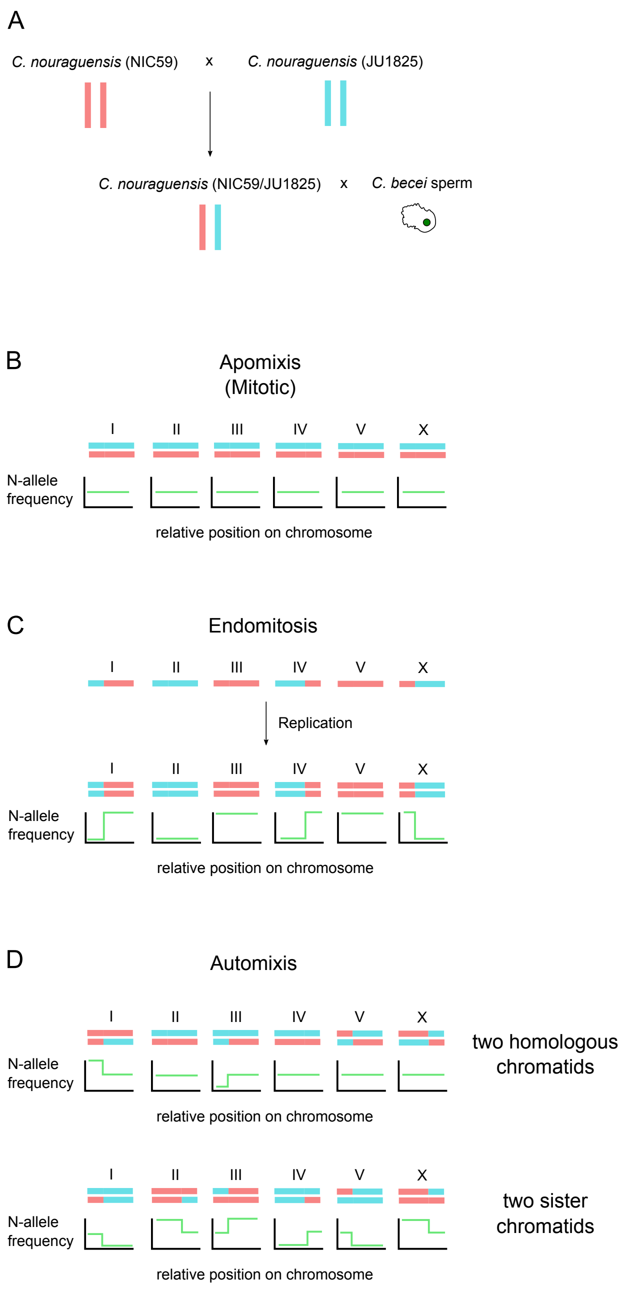


**Figure S6. Genotypic signatures of multiple potential mechanisms of diploid maternal inheritance. (A)** A schematic of the cross used to determine how diploidy is restored in gynogenetically produced offspring in the interspecies cross. Heterozygous (NIC59/JU1825) *C. nouraguensis* females are crossed to *C. becei* (QG711) males. Individual viable offspring resulting from the hybridization undergo whole genome sequencing. **(B)** If diploid maternal inheritance results from mitotic (apomixis) rather than meiotic divisions, then viable F1 are clones of their mother and will be heterozygous NIC59/JU1825 across their entire genome (0.5 NIC59 allele frequency). **(C)** Diploid maternal inheritance can occur through endomitosis, which is when a haploid maternal genome replicates without a cell-division. This results in two exact copies of each chromosome and therefore homozygosity for either NIC59 (1.0 NIC59 allele frequency) or JU1825 alleles (0.0 NIC59 allele frequency) across the entire genome. **(D)** Diploid maternal inheritance can occur through automixis (combining two of the four meiotic products). Combining two homologous chromatids will result in heterozygosity in the center of a chromosome, and heterozygosity or homozygosity at the chromosome ends. Combining two sister chromatids will result in homozygosity in the center of a chromosome and heterozygosity at one of their ends. Inheriting two sister chromatids for one chromosome and two homologous chromatids for another chromosome in the same genome is theoretically possible, but not depicted here.

**Figure S7. Fertile F1 inherit two randomly selected homologous chromatids from each maternal bivalent.** Each of the following pages contains plots describing whole-genome sequencing data of either a rare viable F1 individual or a control DNA sample. The sample name is at the top of each page, along with the individual's sex, fertility and strain it was backcrossed to for fertility testing (if applicable). Each page has five rows of plots. The first row shows the genotypes of the sample's *C. nouraguensis* maternal chromosomes (i.e. average NIC59 allele frequency in 50-kb windows across the *C. nouraguensis* assembly). Haplotype change points and average allele frequency for each segment are shown by the green horizontal lines. The second and third rows show the sample's average read coverage of the *C. becei* and *C. nouraguensis* assemblies in 50-kb windows. The fourth and fifth rows show the average GC content of the *C. becei* and *C. nouraguensis* assemblies in 50-kb windows. The gray vertical lines represent breaks between scaffolds.

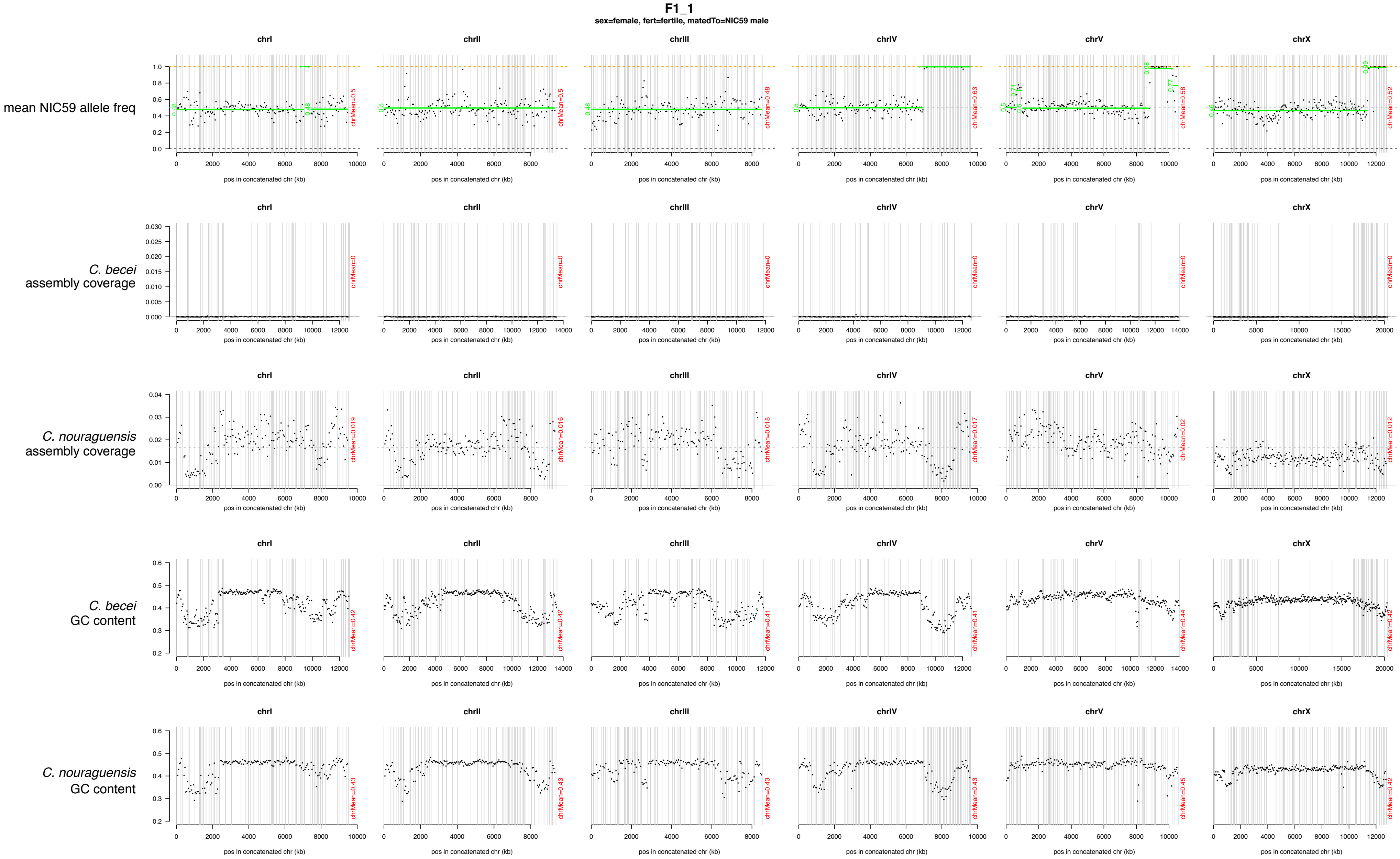
F1_1
sex=female, fert=fertile, matedTo=NIC59 male
mean NIC59 allele freq
C. becei assembly coverage
C. nouraguensis assembly coverage
C. becei GC content
C. nouraguensis GC content
chrI
chrII
chrIII
chrIV
chrV
chrX
pos in concatenated chr (kb)
chrMean=0.5
chrMean=0.5
chrMean=0.48
chrMean=0.63
chrMean=0.58
chrMean=0.52
chrMean=0
chrMean=0.019
chrMean=0.016
chrMean=0.018
chrMean=0.017
chrMean=0.02
chrMean=0.012
chrMean=0.42
chrMean=0.42
chrMean=0.41
chrMean=0.41
chrMean=0.44
chrMean=0.42
chrMean=0.43
chrMean=0.43
chrMean=0.43
chrMean=0.43
chrMean=0.45
chrMean=0.42

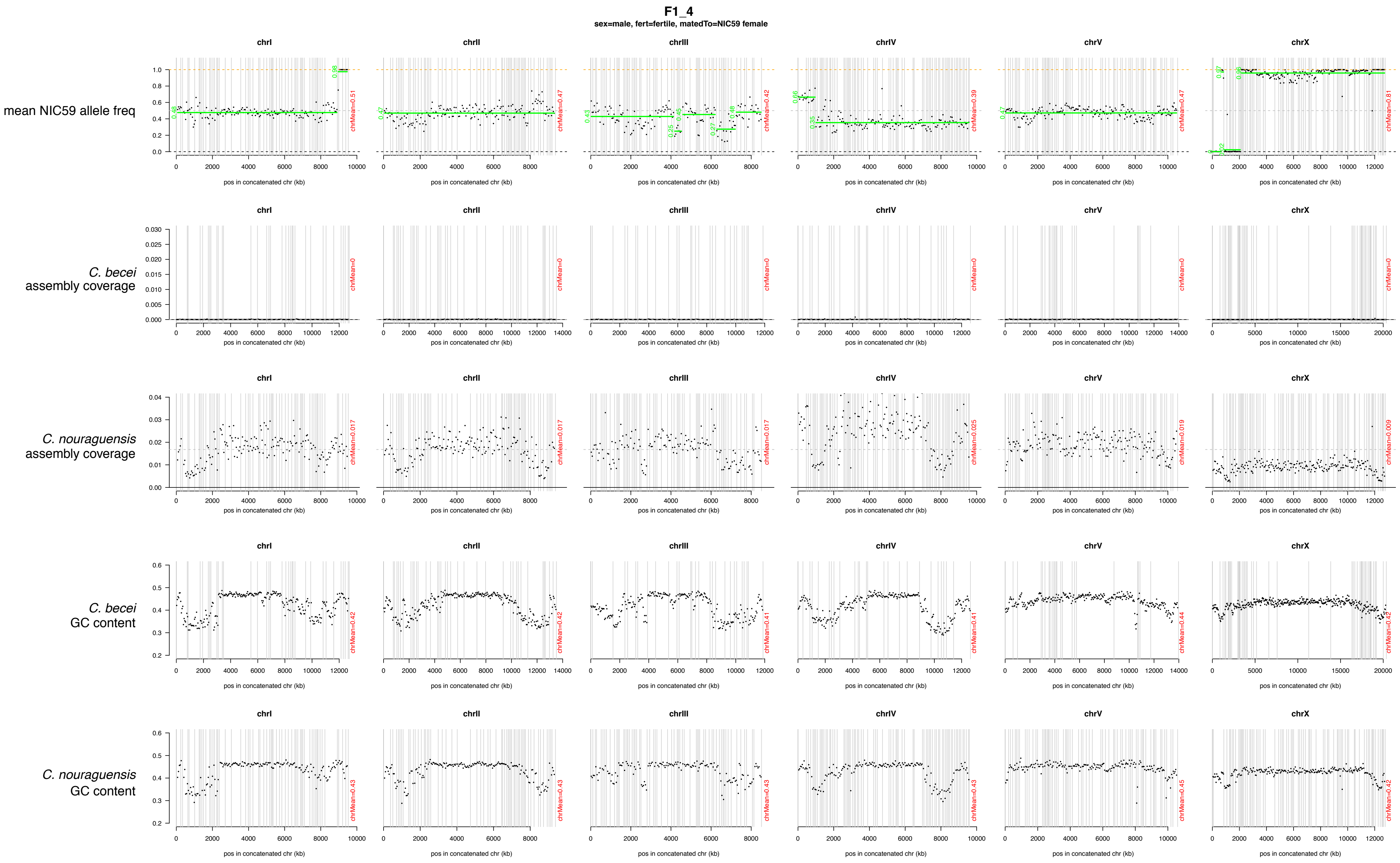
F1_4
sex=male, fert=fertile, matedTo=NIC59 female
mean NIC59 allele freq
C. becei assembly coverage
C. nouraguensis assembly coverage
C. becei GC content
C. nouraguensis GC content
chrI
chrII
chrIII
chrIV
chrV
chrX
pos in concatenated chr (kb)

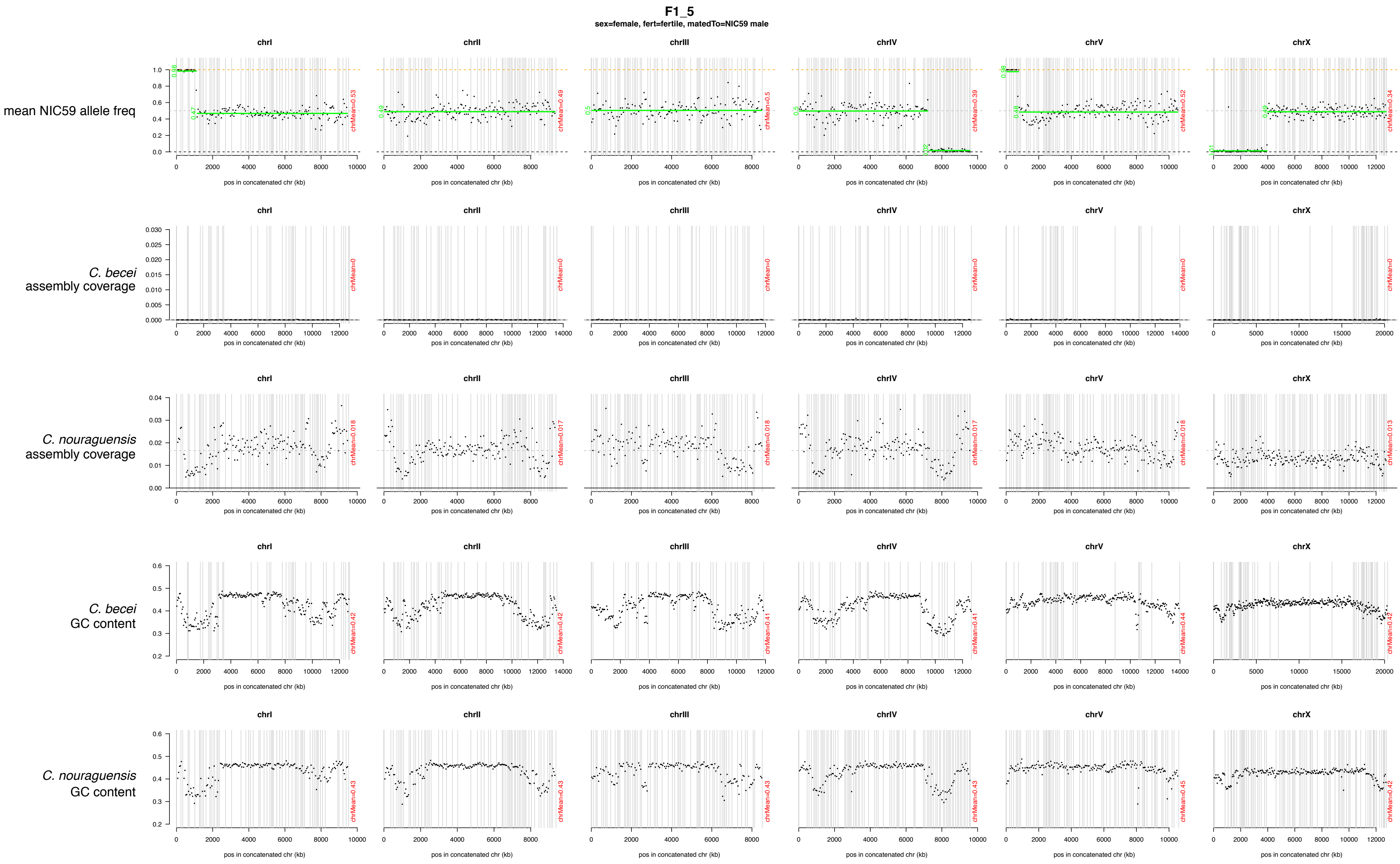

F1_5
sex=female, fert=fertile, matedTo=NIC59 male
mean NIC59 allele freq
C. becei assembly coverage
C. nouraguensis assembly coverage
C. becei GC content
C. nouraguensis GC content
chrI
chrII
chrIII
chrIV
chrV
chrX
pos in concatenated chr (kb)

**F1_6**

sex=female, fert=sterile, matedTo=NIC59 male

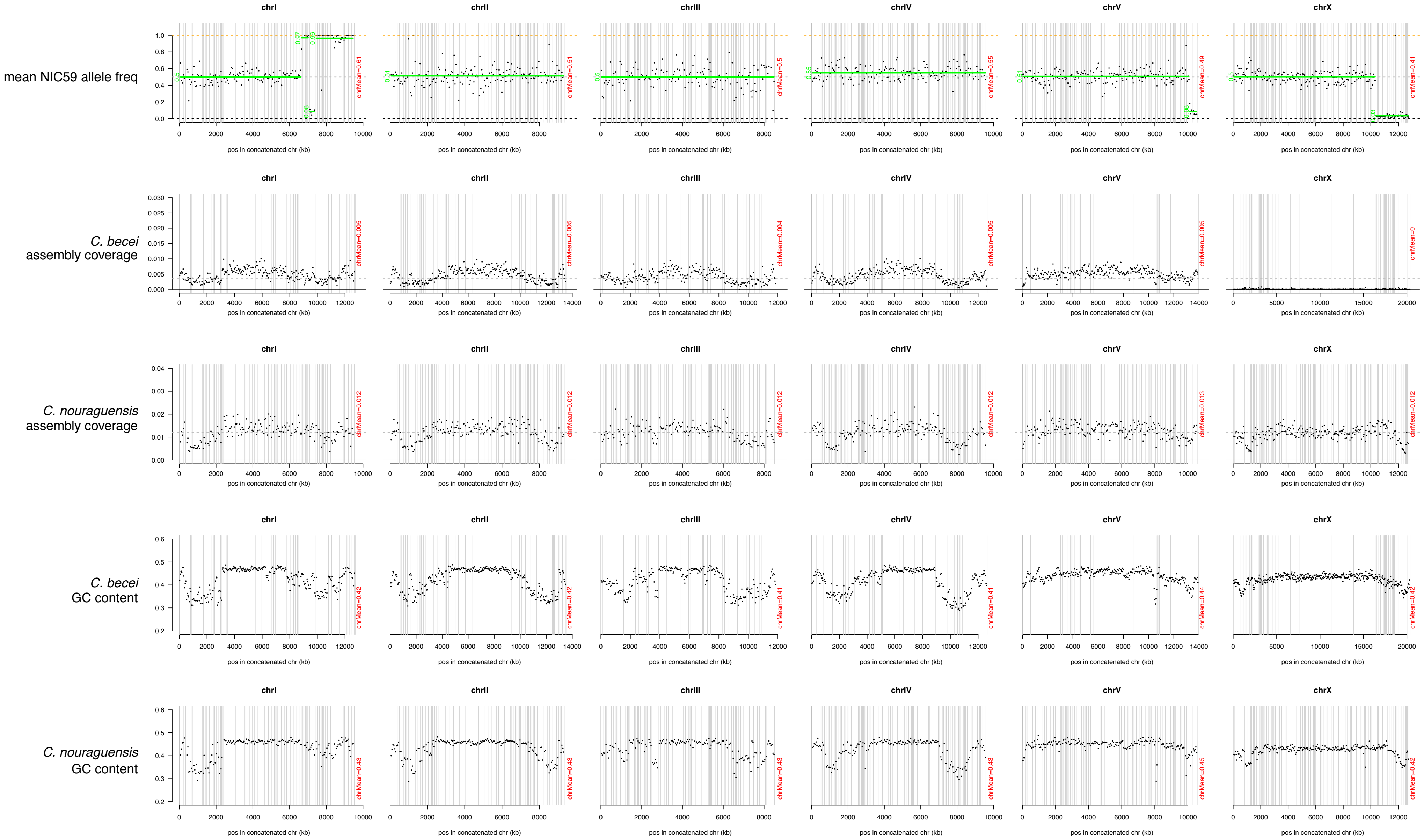

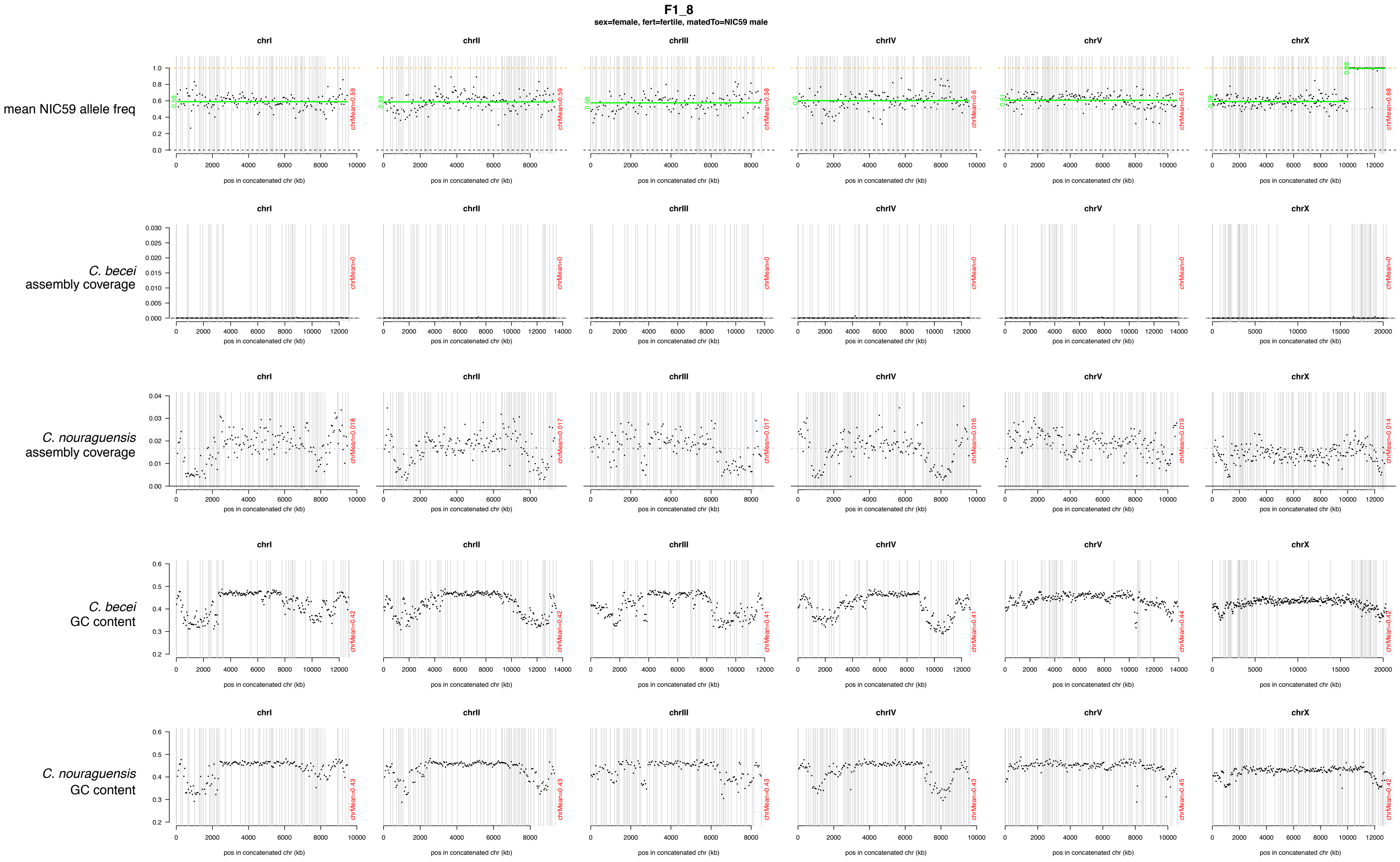
F1_8
sex=female, fert=fertile, matedTo=NIC59 male
mean NIC59 allele freq
C. becei assembly coverage
C. nouraguensis assembly coverage
C. becei GC content
C. nouraguensis GC content
chrI
chrII
chrIII
chrIV
chrV
chrX
pos in concatenated chr (kb)
chrMean=0.59
chrMean=0.59
chrMean=0.58
chrMean=0.6
chrMean=0.61
chrMean=0.68
chrMean=0
chrMean=0.018
chrMean=0.017
chrMean=0.017
chrMean=0.016
chrMean=0.019
chrMean=0.014
chrMean=0.42
chrMean=0.41
chrMean=0.44
chrMean=0.43
chrMean=0.45

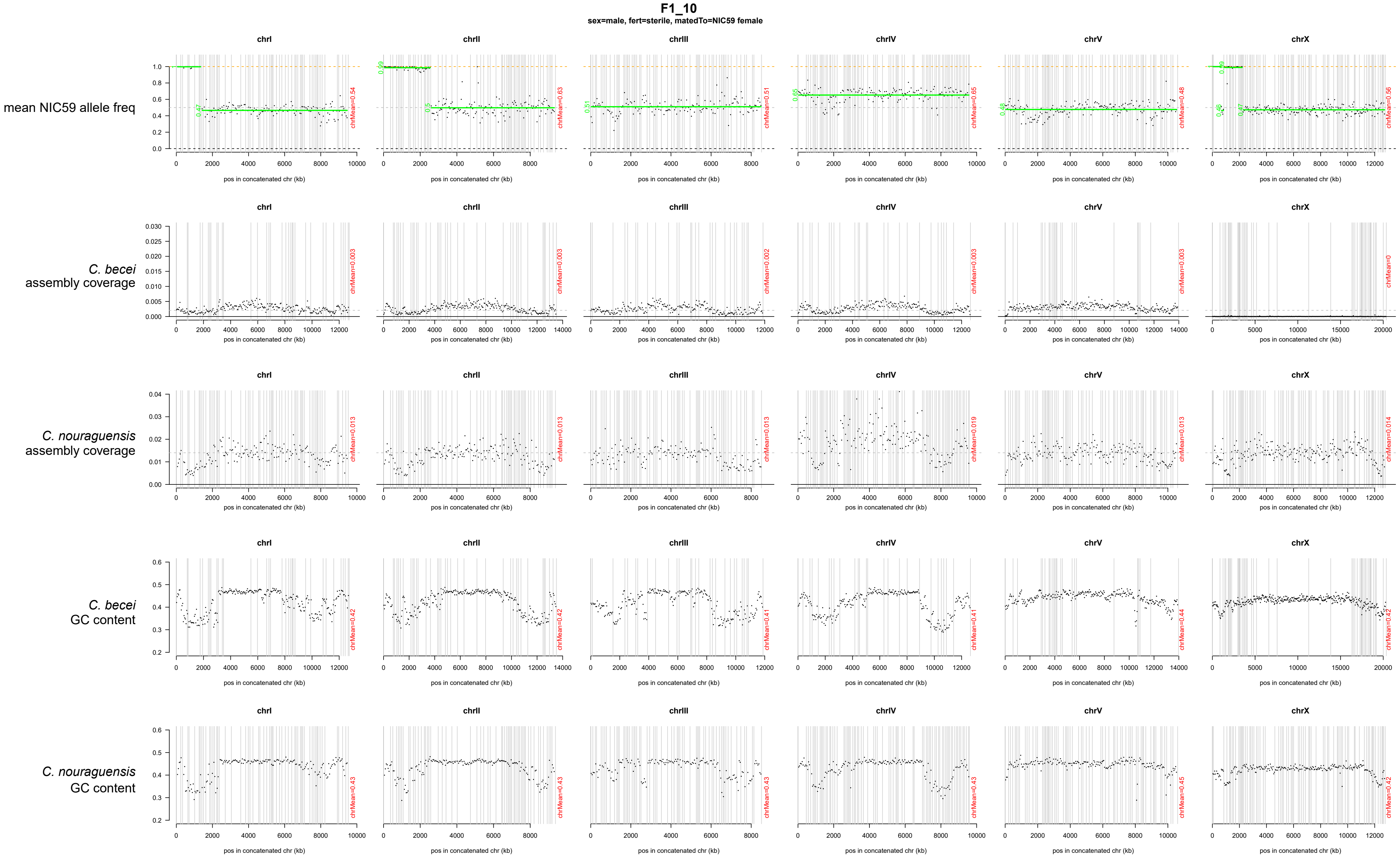
F1_10
sex=male, fert=sterile, matedTo=NIC59 female
mean NIC59 allele freq
C. becei assembly coverage
C. nouraguensis assembly coverage
C. becei GC content
C. nouraguensis GC content
chrI
chrII
chrIII
chrIV
chrV
chrX
pos in concatenated chr (kb)
chrMean=0.54
chrMean=0.63
chrMean=0.51
chrMean=0.65
chrMean=0.48
chrMean=0.56
chrMean=0.003
chrMean=0.003
chrMean=0.002
chrMean=0.003
chrMean=0.003
chrMean=0
chrMean=0.013
chrMean=0.013
chrMean=0.013
chrMean=0.019
chrMean=0.013
chrMean=0.014
chrMean=0.42
chrMean=0.42
chrMean=0.41
chrMean=0.41
chrMean=0.44
chrMean=0.42
chrMean=0.43
chrMean=0.43
chrMean=0.43
chrMean=0.43
chrMean=0.45
chrMean=0.42

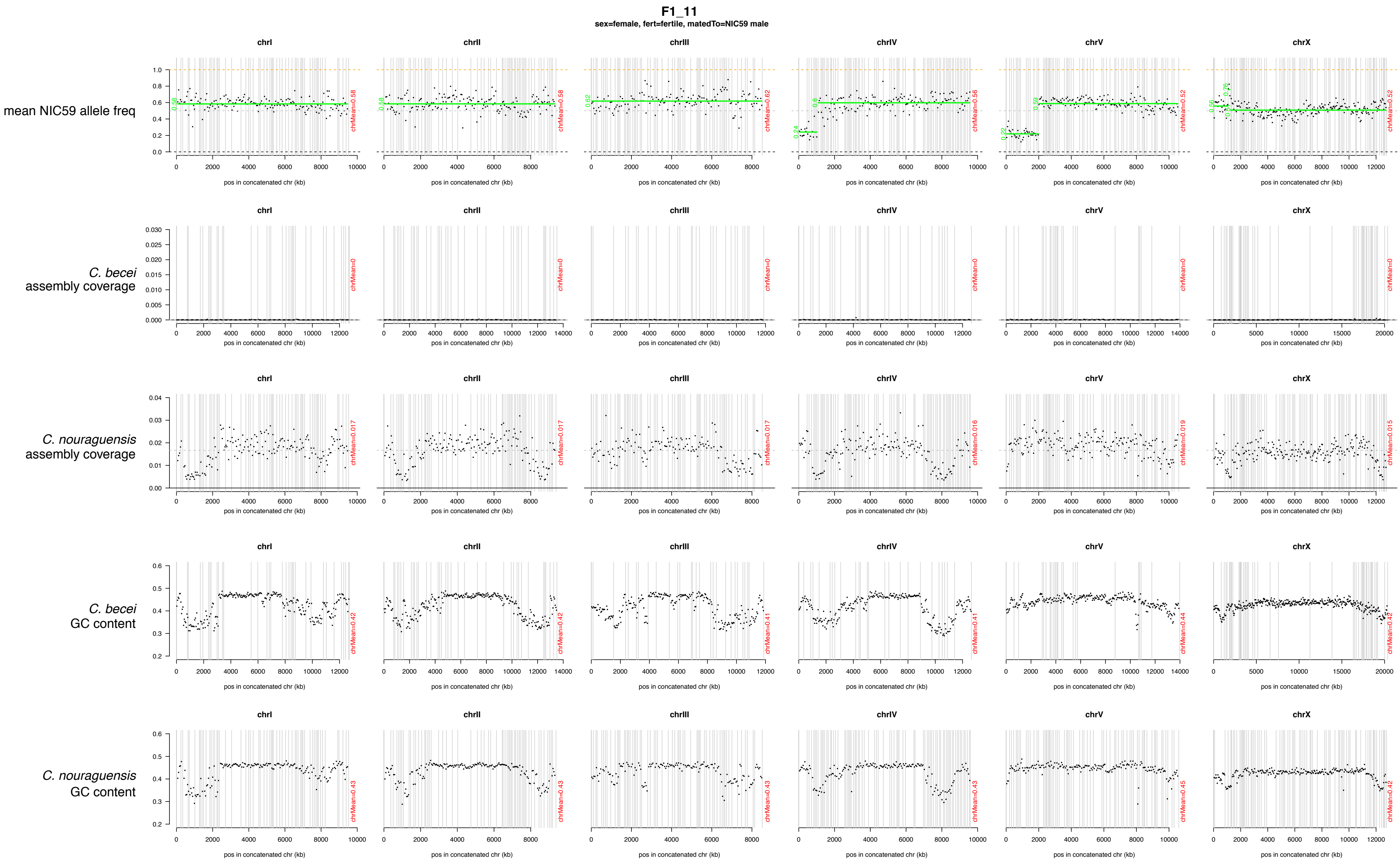
F1_11
sex=female, fert=fertile, matedTo=NIC59 male
chrI
chrII
chrIII
chrIV
chrV
chrX
mean NIC59 allele freq
C. becei assembly coverage
C. nouraguensis assembly coverage
C. becei GC content
C. nouraguensis GC content
pos in concatenated chr (kb)

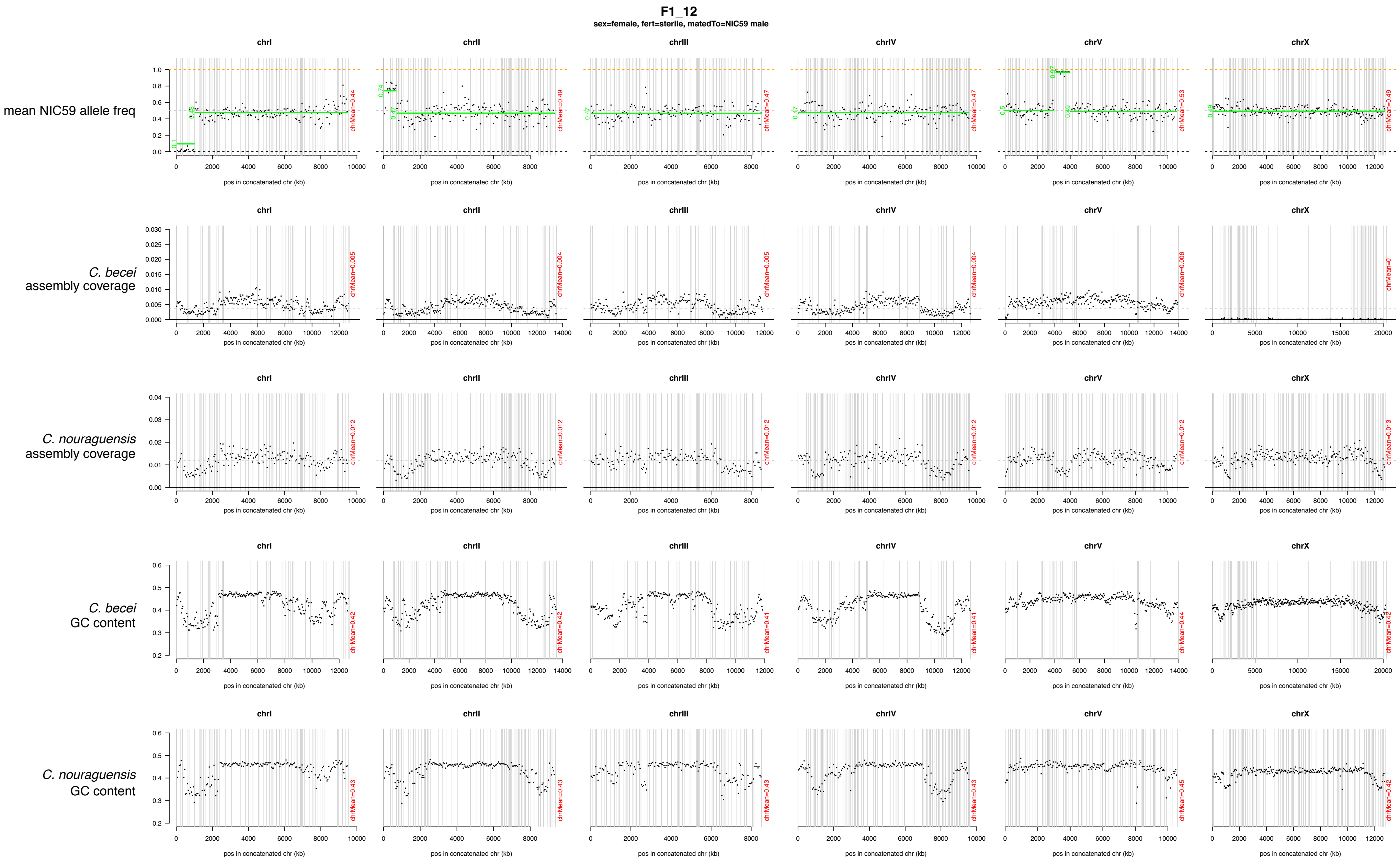
F1_12
sex=female, fert=sterile, matedTo=NIC59 male
mean NIC59 allele freq
C. becei assembly coverage
C. nouraguensis assembly coverage
C. becei GC content
C. nouraguensis GC content
chrI
chrII
chrIII
chrIV
chrV
chrX
pos in concatenated chr (kb)
chrMean=0.44
chrMean=0.49
chrMean=0.47
chrMean=0.47
chrMean=0.53
chrMean=0.49
chrMean=0.005
chrMean=0.004
chrMean=0.005
chrMean=0.004
chrMean=0.006
chrMean=0
chrMean=0.012
chrMean=0.012
chrMean=0.012
chrMean=0.012
chrMean=0.012
chrMean=0.013
chrMean=0.42
chrMean=0.42
chrMean=0.41
chrMean=0.41
chrMean=0.44
chrMean=0.42
chrMean=0.43
chrMean=0.43
chrMean=0.43
chrMean=0.43
chrMean=0.45
chrMean=0.42

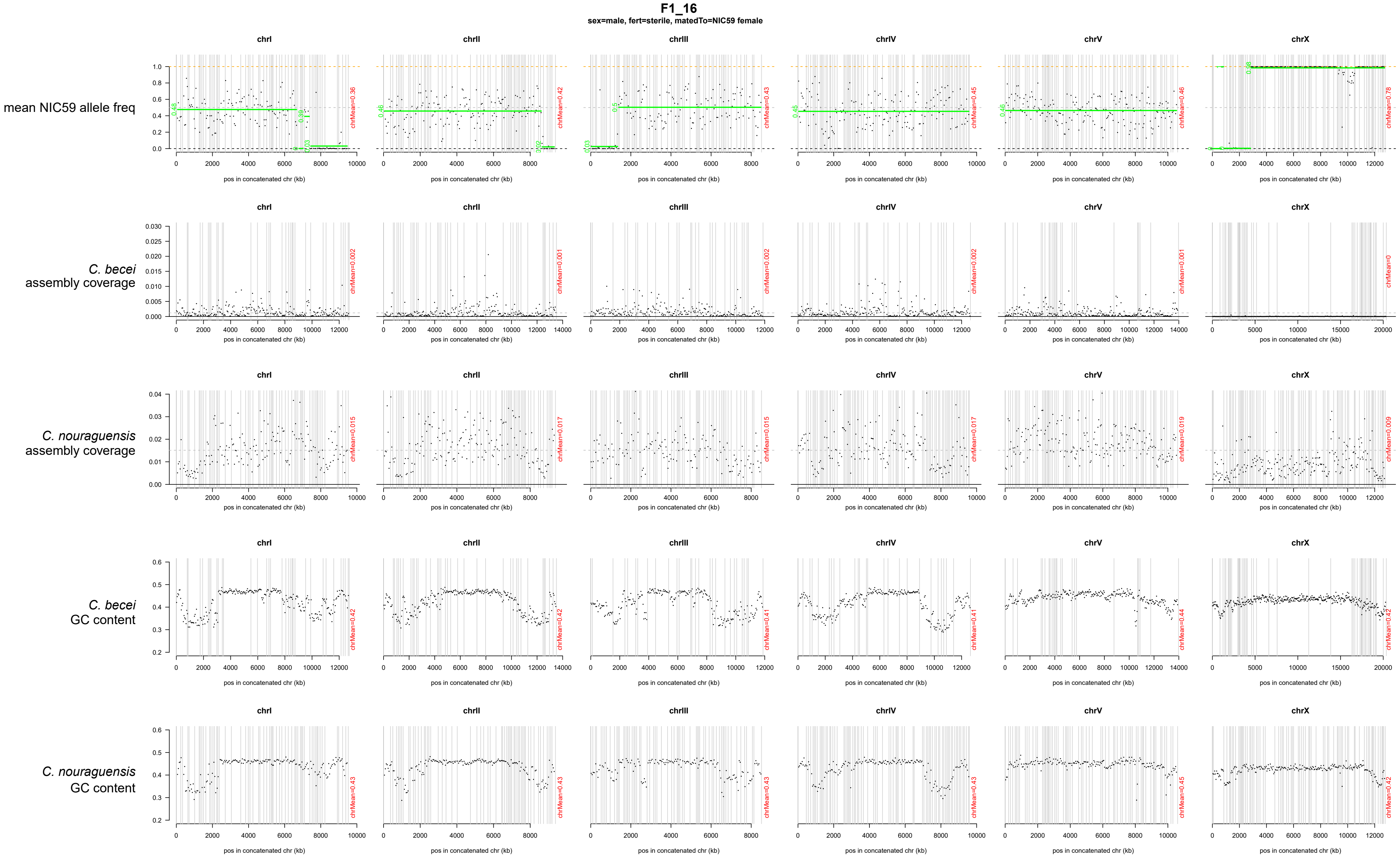

F1_16
sex=male, fert=sterile, matedTo=NIC59 female
chrI
chrII
chrIII
chrIV
chrV
chrX
mean NIC59 allele freq
C. becei assembly coverage
C. nouraguensis assembly coverage
C. becei GC content
C. nouraguensis GC content
pos in concatenated chr (kb)
chrMean=0.36
chrMean=0.42
chrMean=0.43
chrMean=0.45
chrMean=0.46
chrMean=0.78
chrMean=0.002
chrMean=0.001
chrMean=0.002
chrMean=0.002
chrMean=0.001
chrMean=0
chrMean=0.015
chrMean=0.017
chrMean=0.015
chrMean=0.017
chrMean=0.019
chrMean=0.009
chrMean=0.42
chrMean=0.42
chrMean=0.41
chrMean=0.41
chrMean=0.44
chrMean=0.42
chrMean=0.43
chrMean=0.43
chrMean=0.43
chrMean=0.43
chrMean=0.45
chrMean=0.42

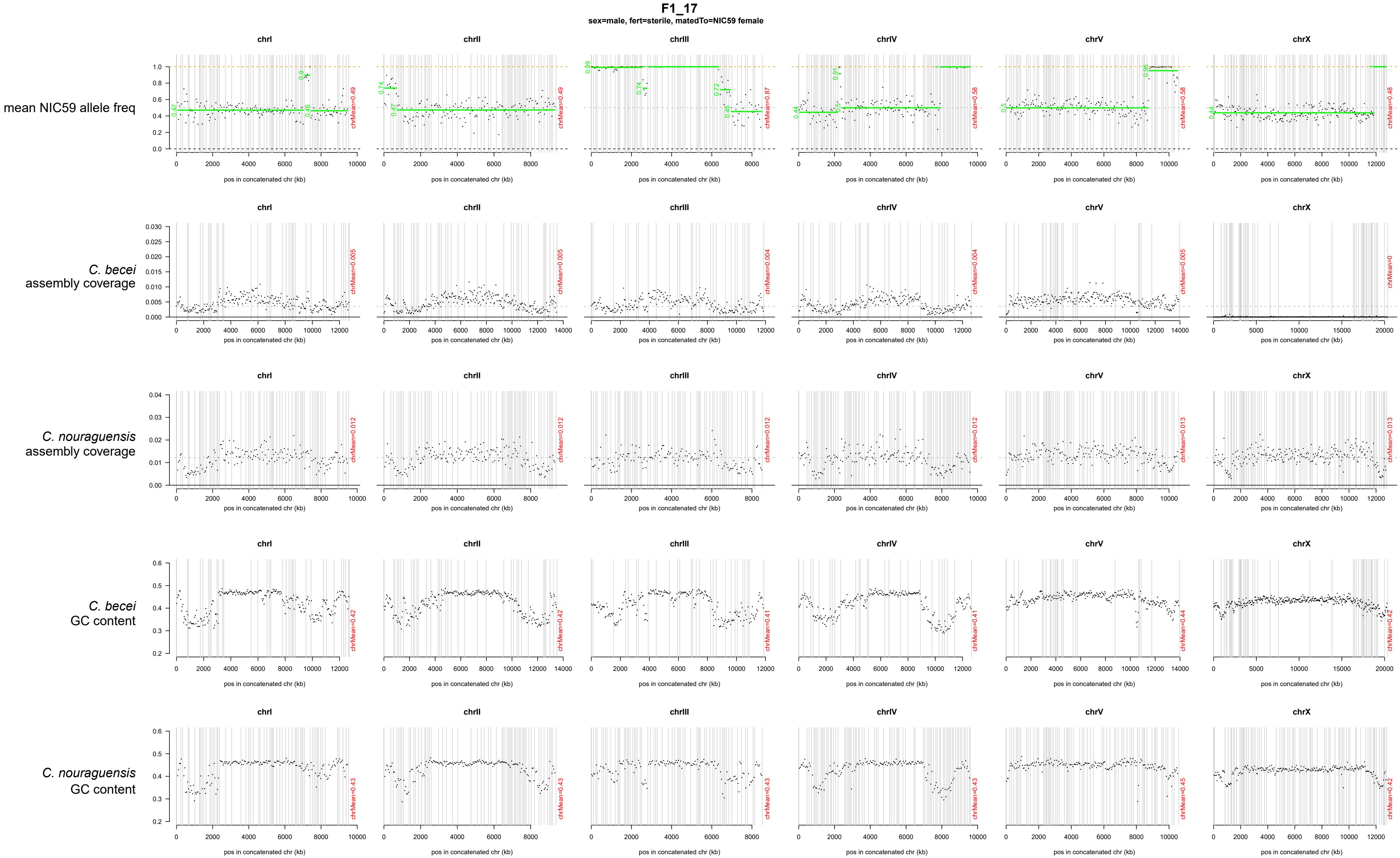
F1_17
sex=male, fert=sterile, matedTo=NIC59 female
chrI
chrII
chrIII
chrIV
chrV
chrX
mean NIC59 allele freq
C. becei assembly coverage
C. nouraguensis assembly coverage
C. becei GC content
C. nouraguensis GC content
pos in concatenated chr (kb)

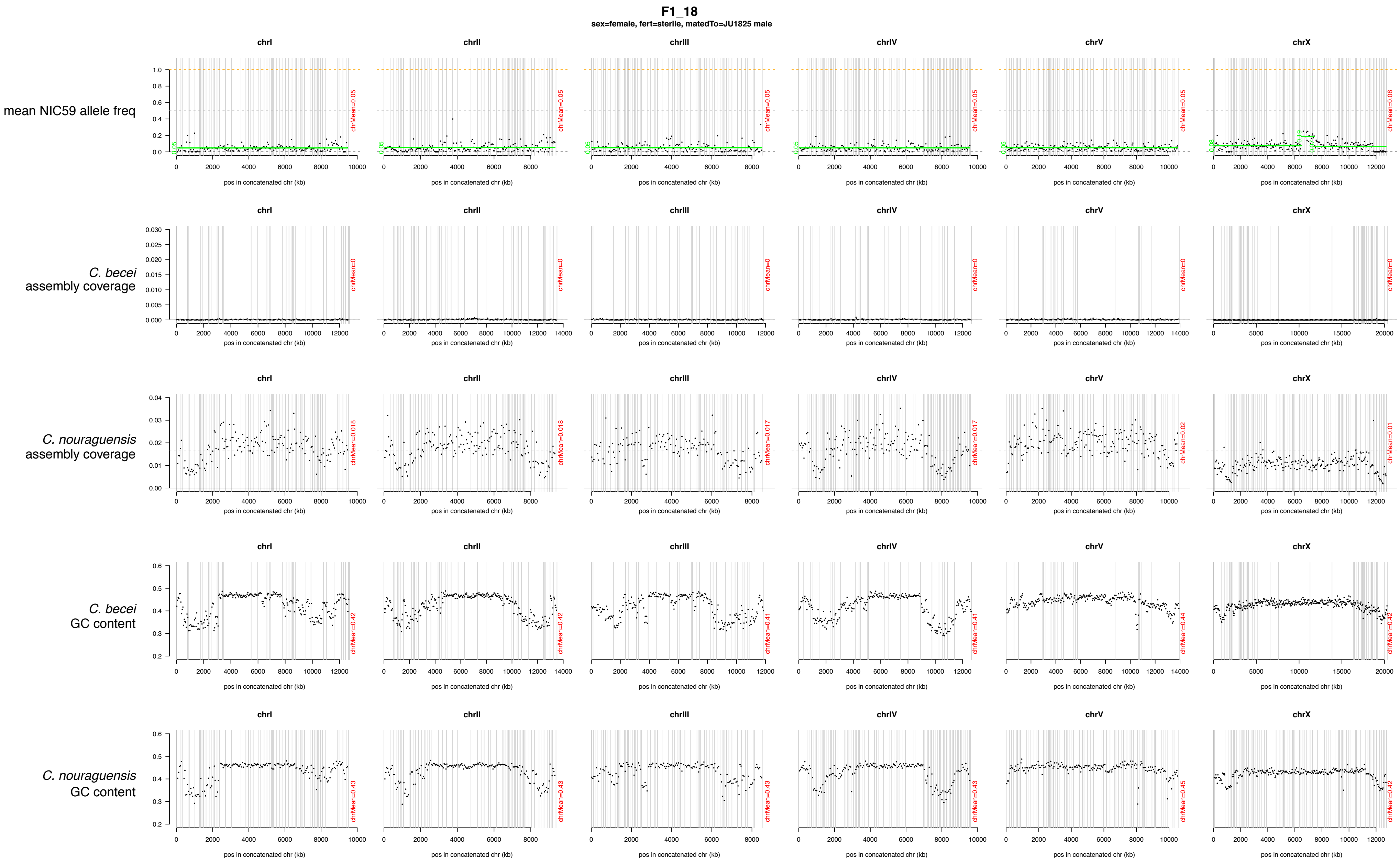

F1_18
sex=female, fert=sterile, matedTo=JU1825 male
mean NIC59 allele freq
C. becei assembly coverage
C. nouraguensis assembly coverage
C. becei GC content
C. nouraguensis GC content
chrI
chrII
chrIII
chrIV
chrV
chrX
pos in concatenated chr (kb)

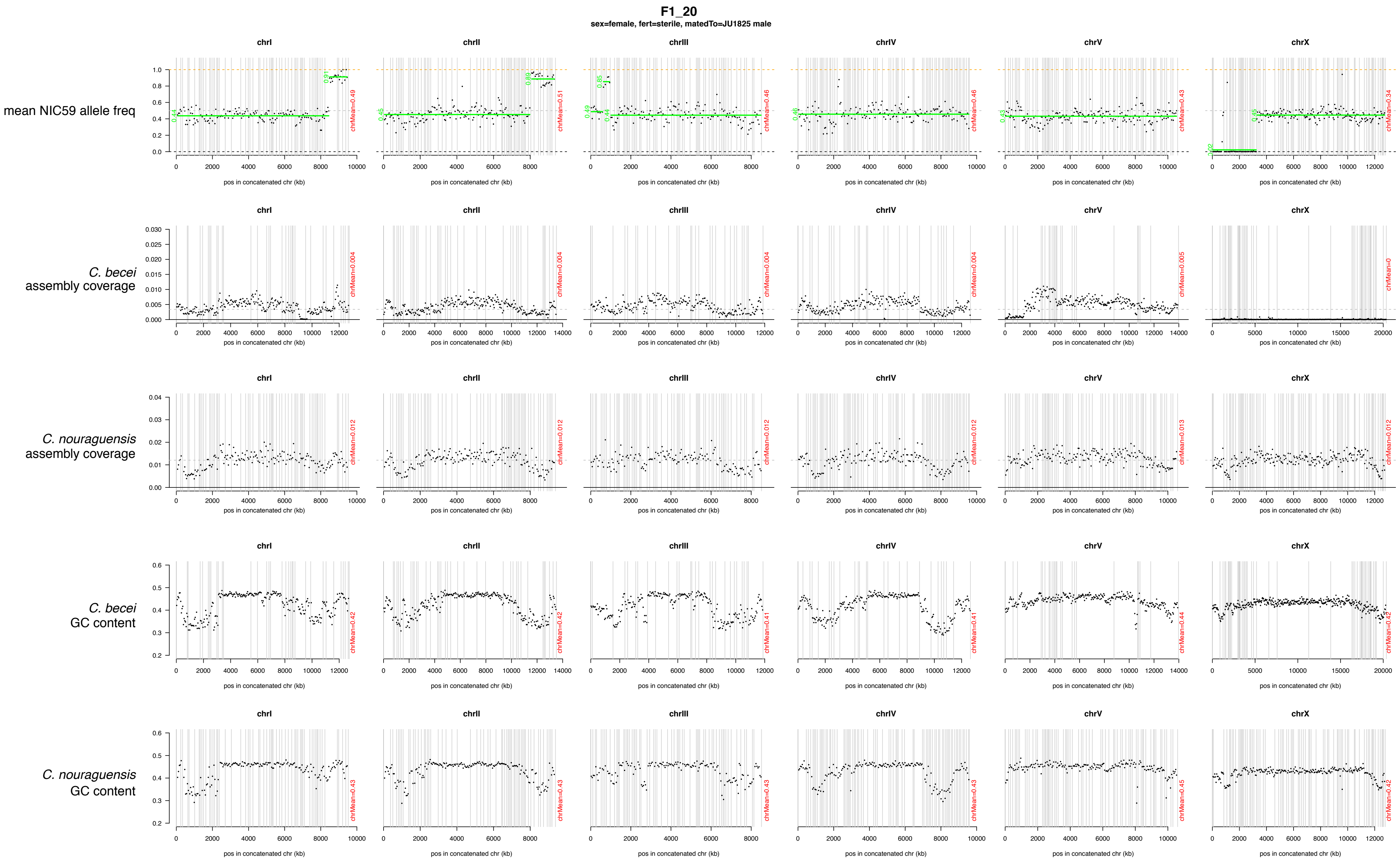
F1_20
sex=female, fert=sterile, matedTo=JU1825 male
chrI
chrII
chrIII
chrIV
chrV
chrX
mean NIC59 allele freq
C. becei
assembly coverage
C. nouraguensis
assembly coverage
C. becei
GC content
C. nouraguensis
GC content
pos in concatenated chr (kb)
chrMean=0.49
chrMean=0.51
chrMean=0.46
chrMean=0.46
chrMean=0.43
chrMean=0.34
chrMean=0.004
chrMean=0.004
chrMean=0.004
chrMean=0.004
chrMean=0.005
chrMean=0
chrMean=0.012
chrMean=0.012
chrMean=0.012
chrMean=0.012
chrMean=0.013
chrMean=0.012
chrMean=0.42
chrMean=0.42
chrMean=0.41
chrMean=0.41
chrMean=0.44
chrMean=0.42
chrMean=0.43
chrMean=0.43
chrMean=0.43
chrMean=0.43
chrMean=0.45
chrMean=0.42

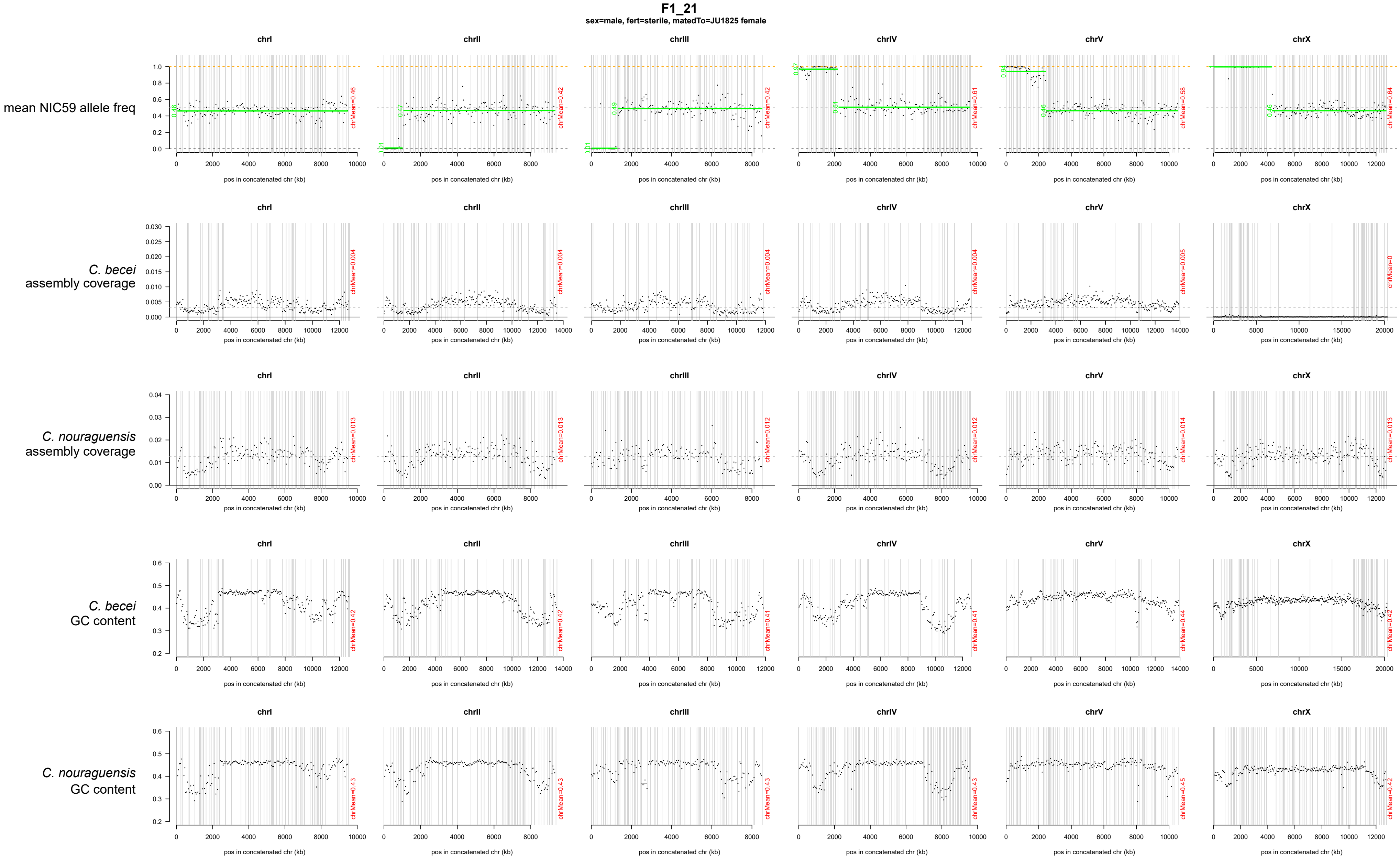
F1_21
sex=male, fert=sterile, matedTo=JU1825 female
mean NIC59 allele freq
C. becei assembly coverage
C. nouraguensis assembly coverage
C. becei GC content
C. nouraguensis GC content
chrI
chrII
chrIII
chrIV
chrV
chrX
pos in concatenated chr (kb)

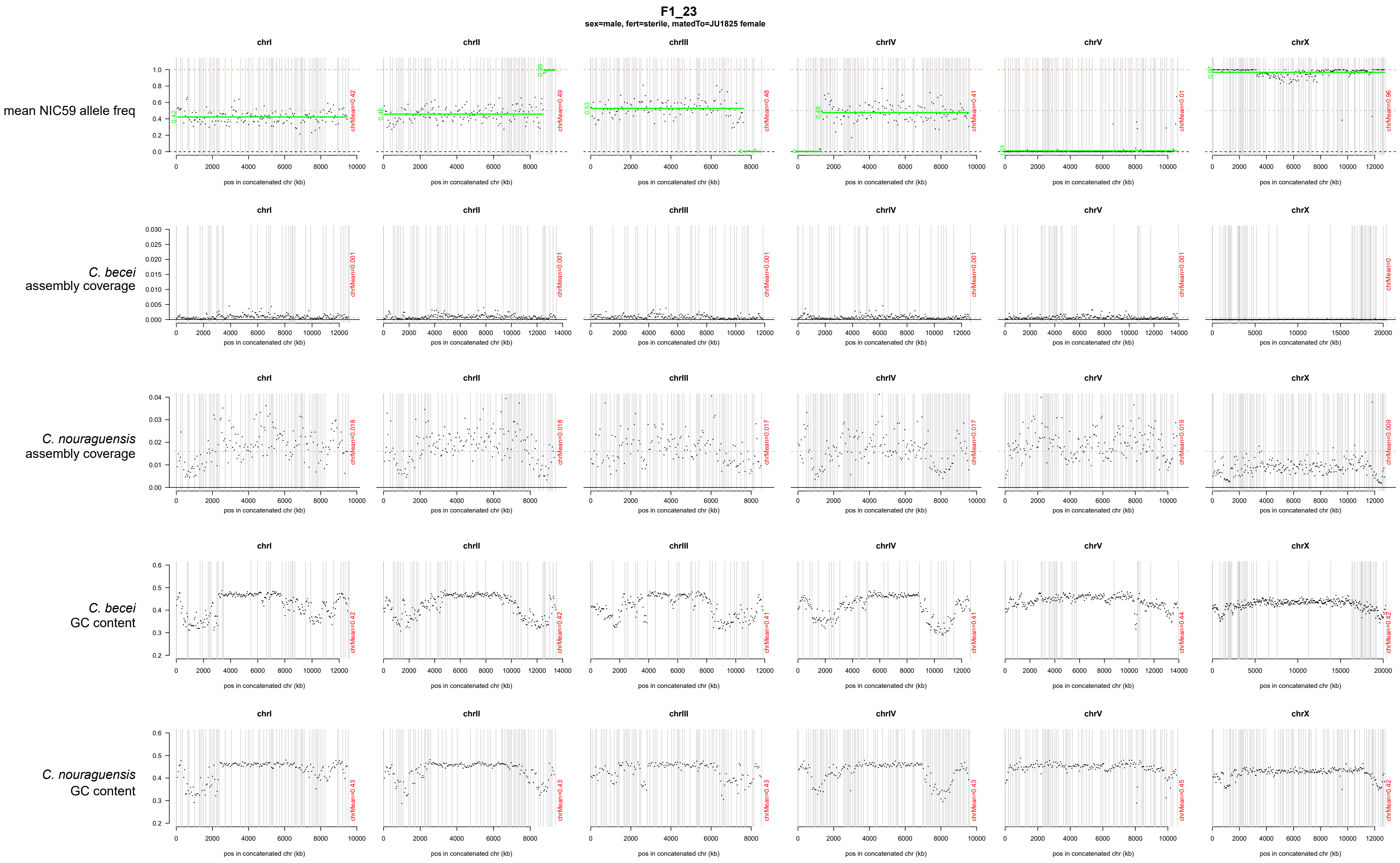
F1_23
sex=male, fert=sterile, matedTo=JU1825 female
mean NIC59 allele freq
C. becei assembly coverage
C. nouraguensis assembly coverage
C. becei GC content
C. nouraguensis GC content
chrI
chrII
chrIII
chrIV
chrV
chrX
pos in concatenated chr (kb)
chrMean=0.42
chrMean=0.49
chrMean=0.48
chrMean=0.41
chrMean=0.01
chrMean=0.96
chrMean=0.001
chrMean=0
chrMean=0.018
chrMean=0.017
chrMean=0.019
chrMean=0.009
chrMean=0.44
chrMean=0.43
chrMean=0.45

**F1_25**

**sex=female, fert=fertile, matedTo=JU1825 male**

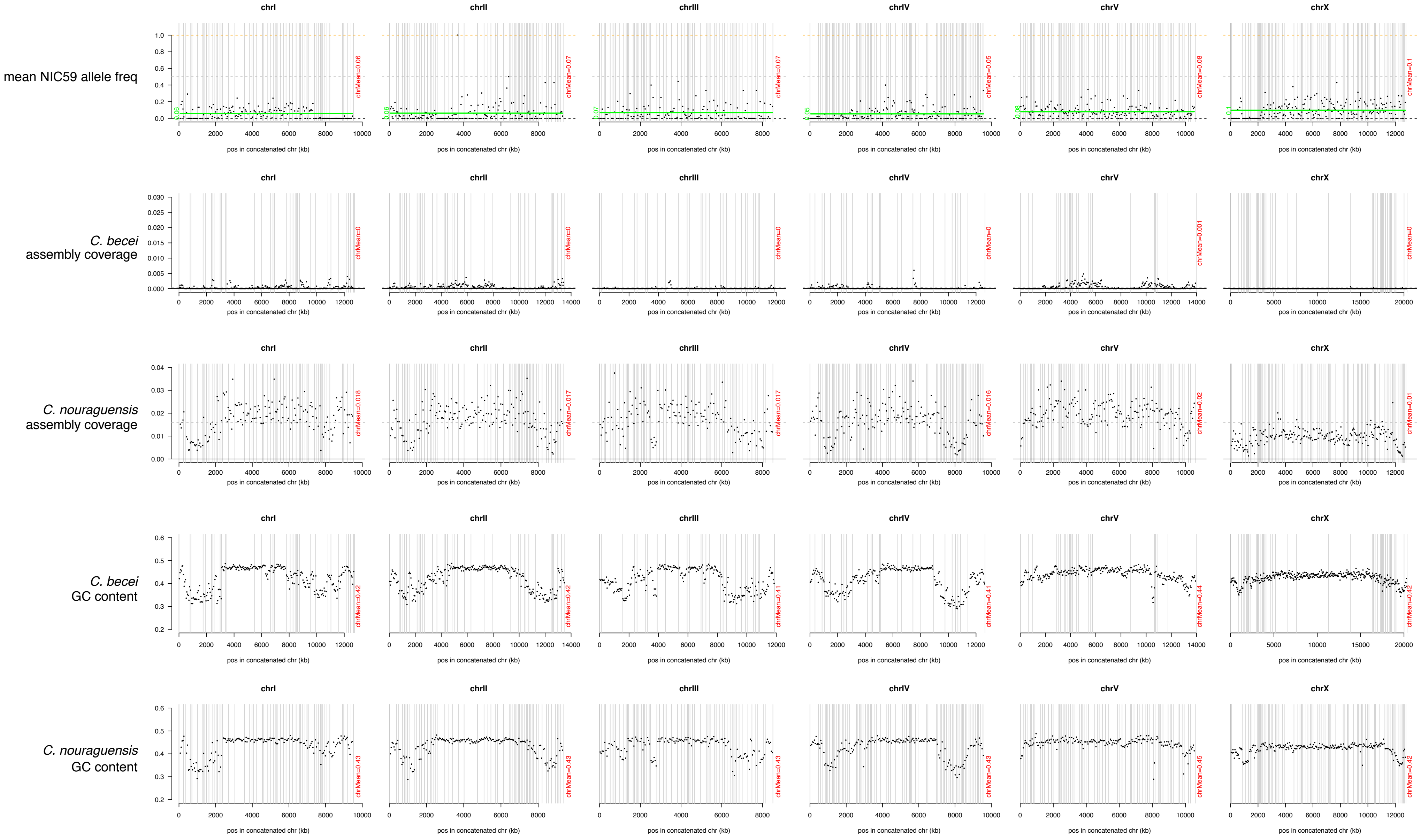

**F1_26**

**sex=female, fert=sterile, matedTo=JU1825 male**

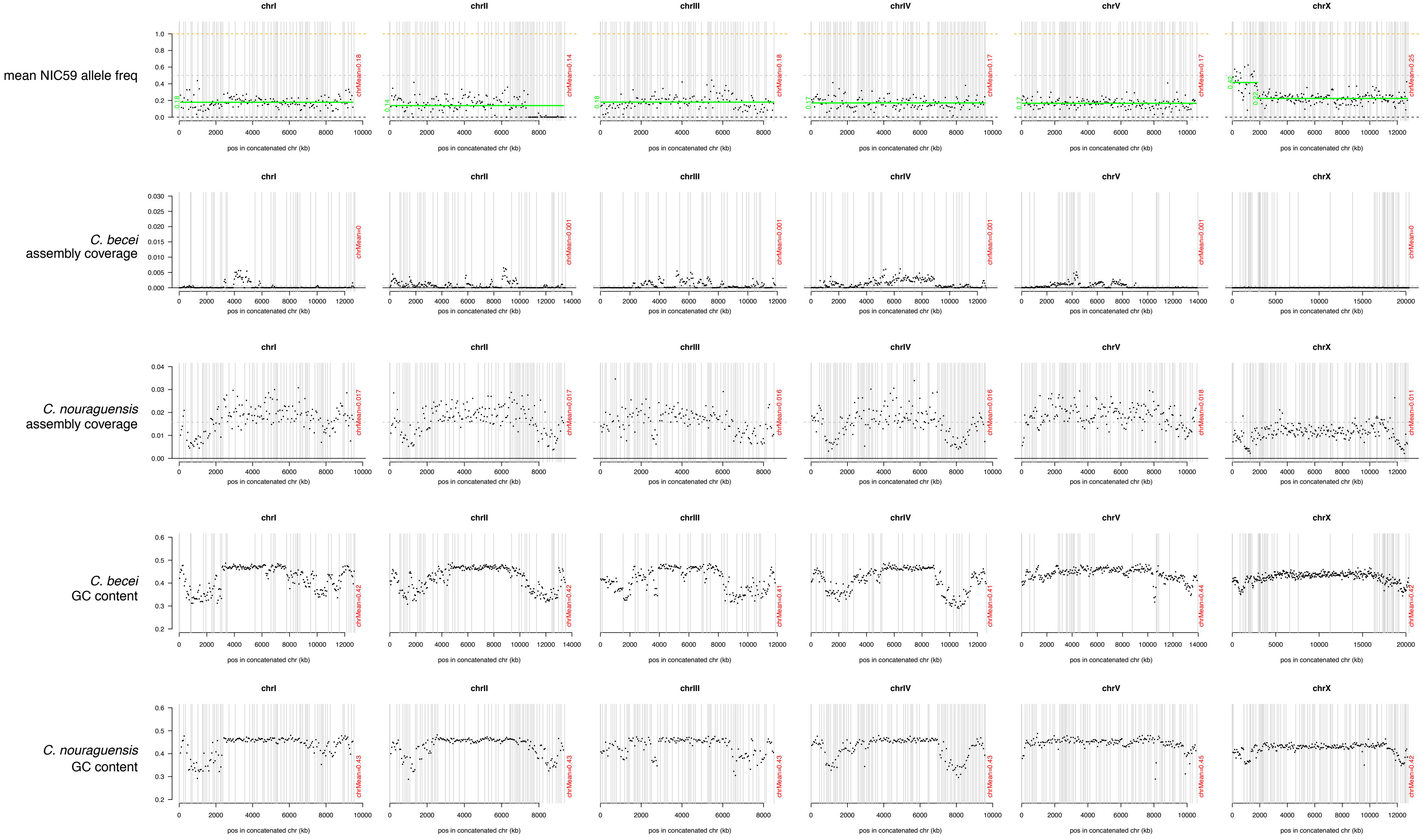

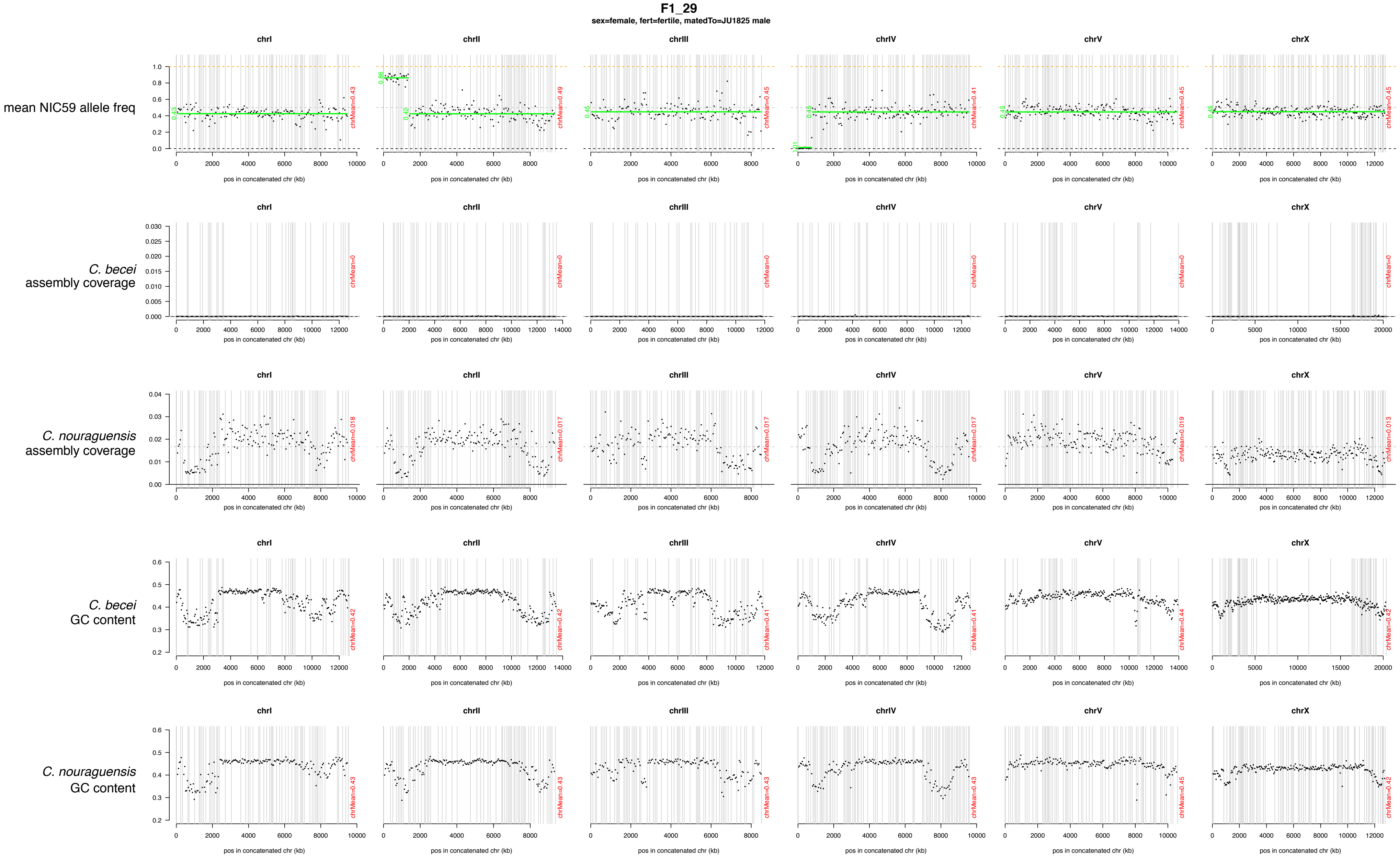
F1_29
sex=female, fert=fertile, matedTo=JU1825 male
chrI
chrII
chrIII
chrIV
chrV
chrX
mean NIC59 allele freq
C. becei assembly coverage
C. nouraguensis assembly coverage
C. becei GC content
C. nouraguensis GC content
pos in concatenated chr (kb)
chrMean=0.43
chrMean=0.49
chrMean=0.45
chrMean=0.41
chrMean=0.45
chrMean=0.45
chrMean=0
chrMean=0.018
chrMean=0.017
chrMean=0.017
chrMean=0.017
chrMean=0.019
chrMean=0.013
chrMean=0.42
chrMean=0.42
chrMean=0.41
chrMean=0.41
chrMean=0.44
chrMean=0.42
chrMean=0.43
chrMean=0.43
chrMean=0.43
chrMean=0.43
chrMean=0.45
chrMean=0.42

**F1_39**

sex=female, fert=fertile, matedTo=JU1825 male

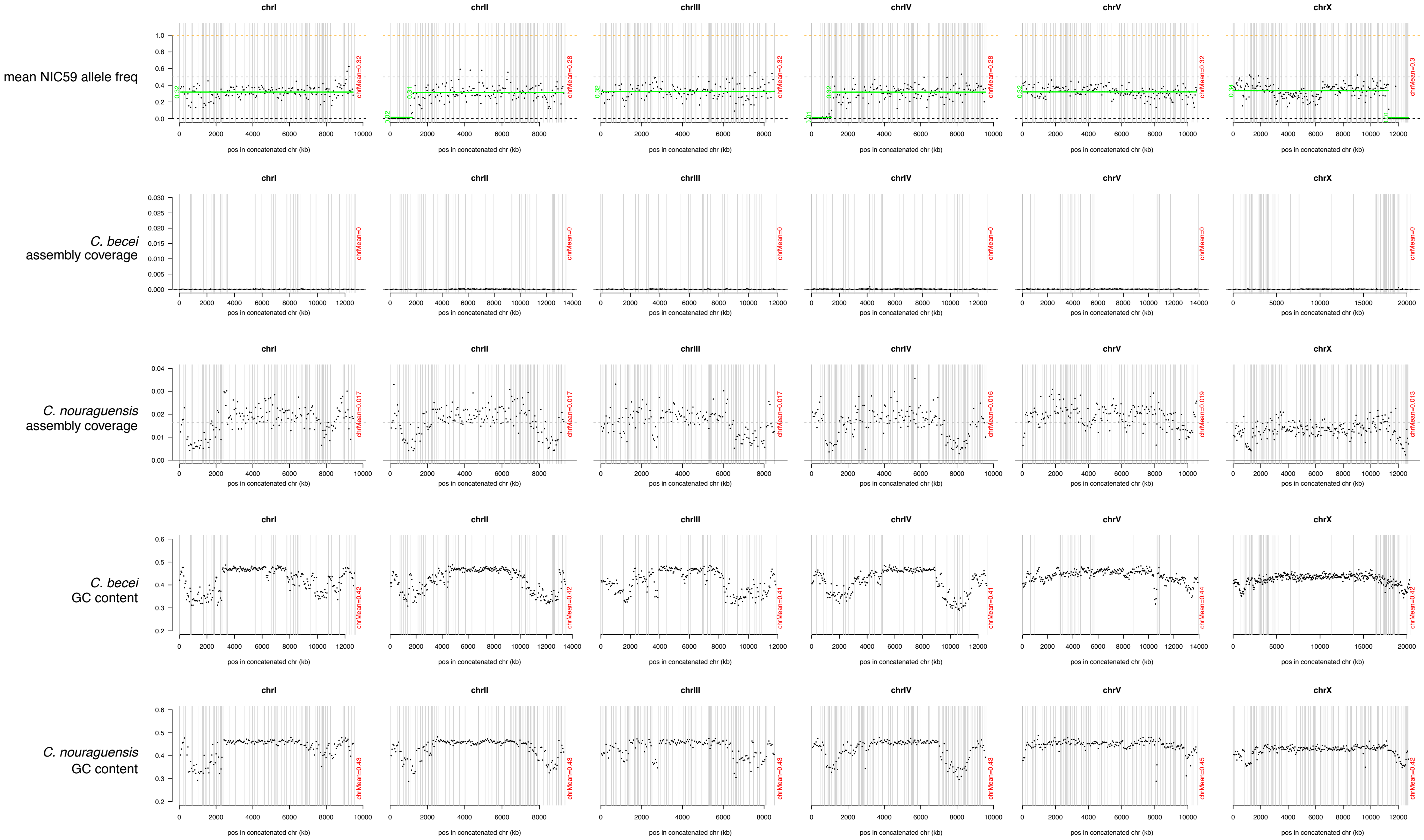

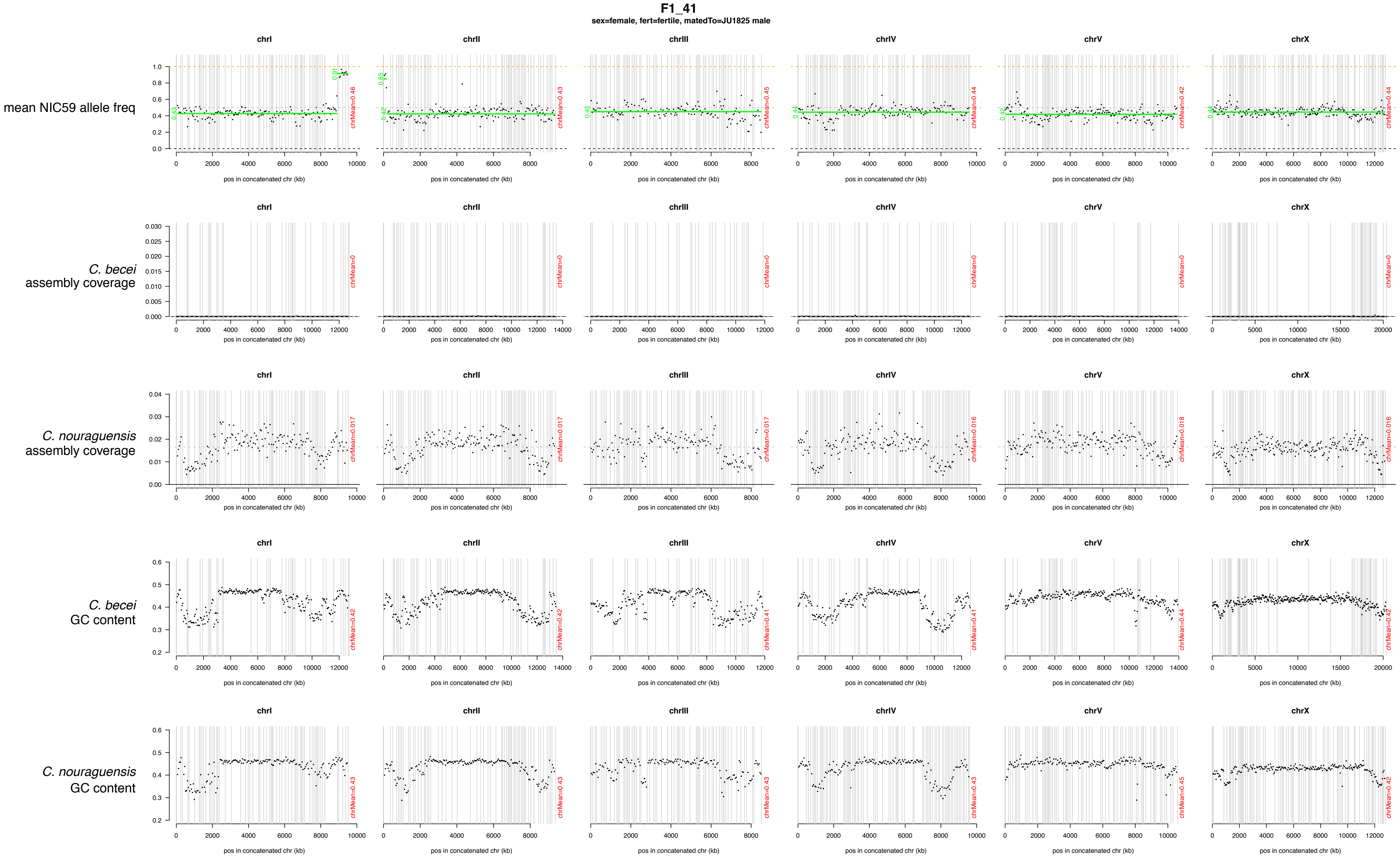
F1_41
sex=female, fert=fertile, matedTo=JU1825 male
mean NIC59 allele freq
C. becei assembly coverage
C. nouraguensis assembly coverage
C. becei GC content
C. nouraguensis GC content
chrI
chrII
chrIII
chrIV
chrV
chrX
pos in concatenated chr (kb)
chrMean=0.46
chrMean=0.43
chrMean=0.45
chrMean=0.44
chrMean=0.42
chrMean=0.44
chrMean=0
chrMean=0.017
chrMean=0.016
chrMean=0.018
chrMean=0.42
chrMean=0.41
chrMean=0.44
chrMean=0.43
chrMean=0.45

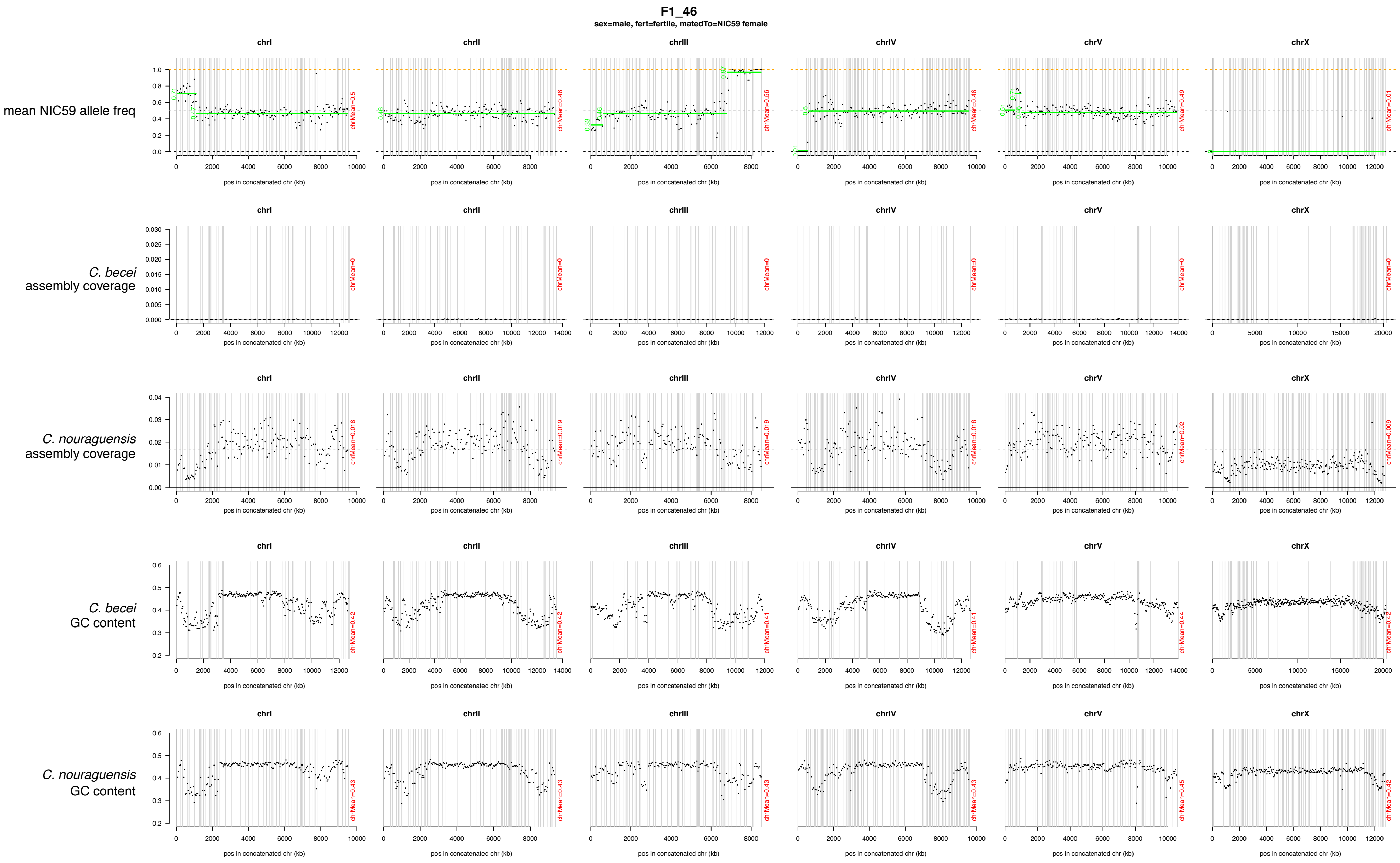

F1_46
sex=male, fert=fertile, matedTo=NIC59 female
chrI
chrII
chrIII
chrIV
chrV
chrX
mean NIC59 allele freq
chrMean=0.5
chrMean=0.46
chrMean=0.56
chrMean=0.46
chrMean=0.49
chrMean=0.01
C. becei assembly coverage
chrMean=0
C. nouraguensis assembly coverage
chrMean=0.018
chrMean=0.019
chrMean=0.019
chrMean=0.018
chrMean=0.02
chrMean=0.009
C. becei GC content
chrMean=0.42
chrMean=0.42
chrMean=0.41
chrMean=0.41
chrMean=0.44
chrMean=0.42
C. nouraguensis GC content
chrMean=0.43
chrMean=0.43
chrMean=0.43
chrMean=0.43
chrMean=0.45
chrMean=0.42
pos in concatenated chr (kb)

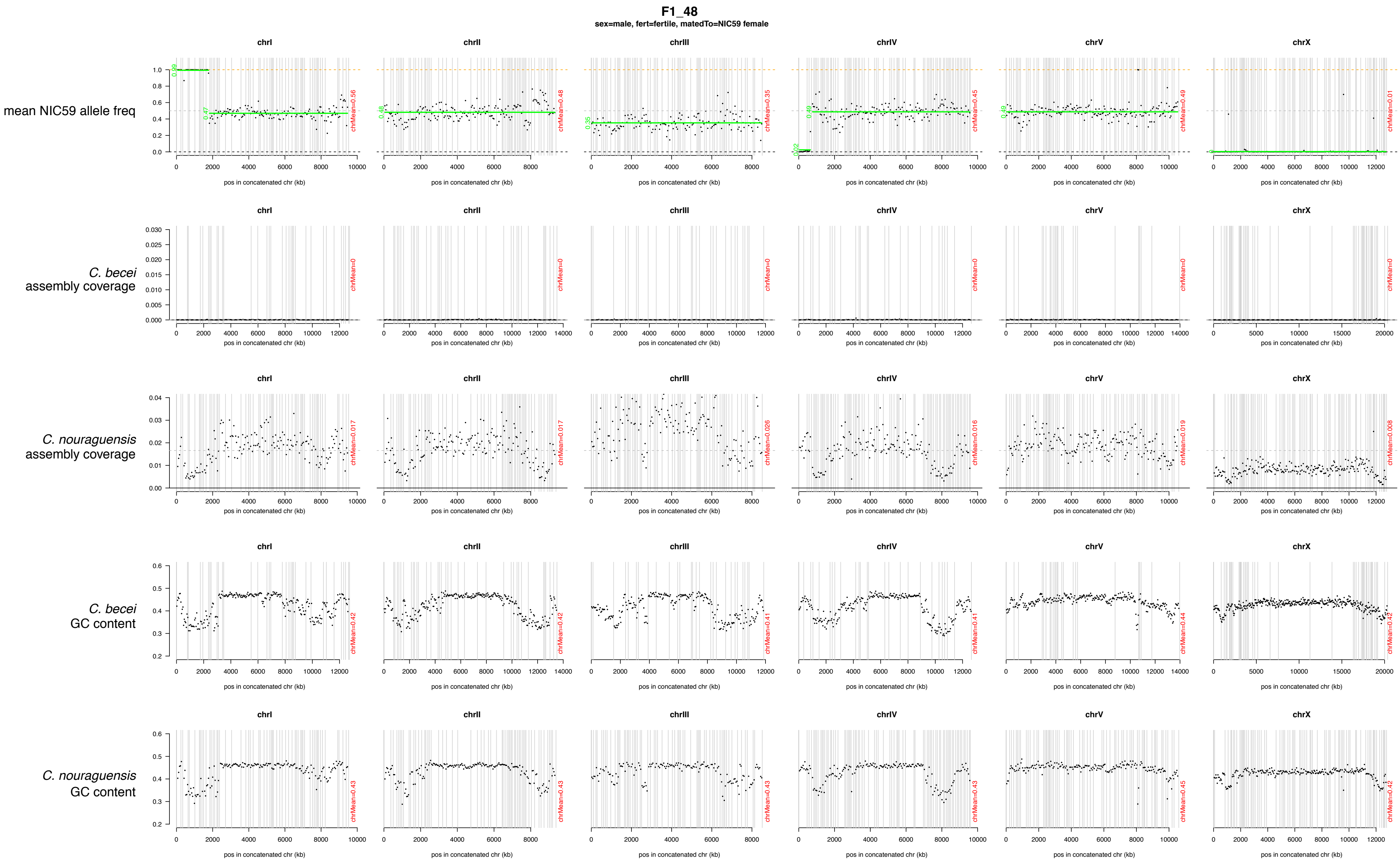
F1_48
sex=male, fert=fertile, matedTo=NIC59 female
chrI
chrII
chrIII
chrIV
chrV
chrX
mean NIC59 allele freq
C. becei
assembly coverage
C. nouraguensis
assembly coverage
C. becei
GC content
C. nouraguensis
GC content
pos in concatenated chr (kb)
chrMean=0.56
chrMean=0.48
chrMean=0.35
chrMean=0.45
chrMean=0.49
chrMean=0.01
chrMean=0
chrMean=0.017
chrMean=0.017
chrMean=0.026
chrMean=0.016
chrMean=0.019
chrMean=0.008
chrMean=0.42
chrMean=0.42
chrMean=0.41
chrMean=0.41
chrMean=0.44
chrMean=0.42
chrMean=0.43
chrMean=0.43
chrMean=0.43
chrMean=0.43
chrMean=0.45
chrMean=0.42

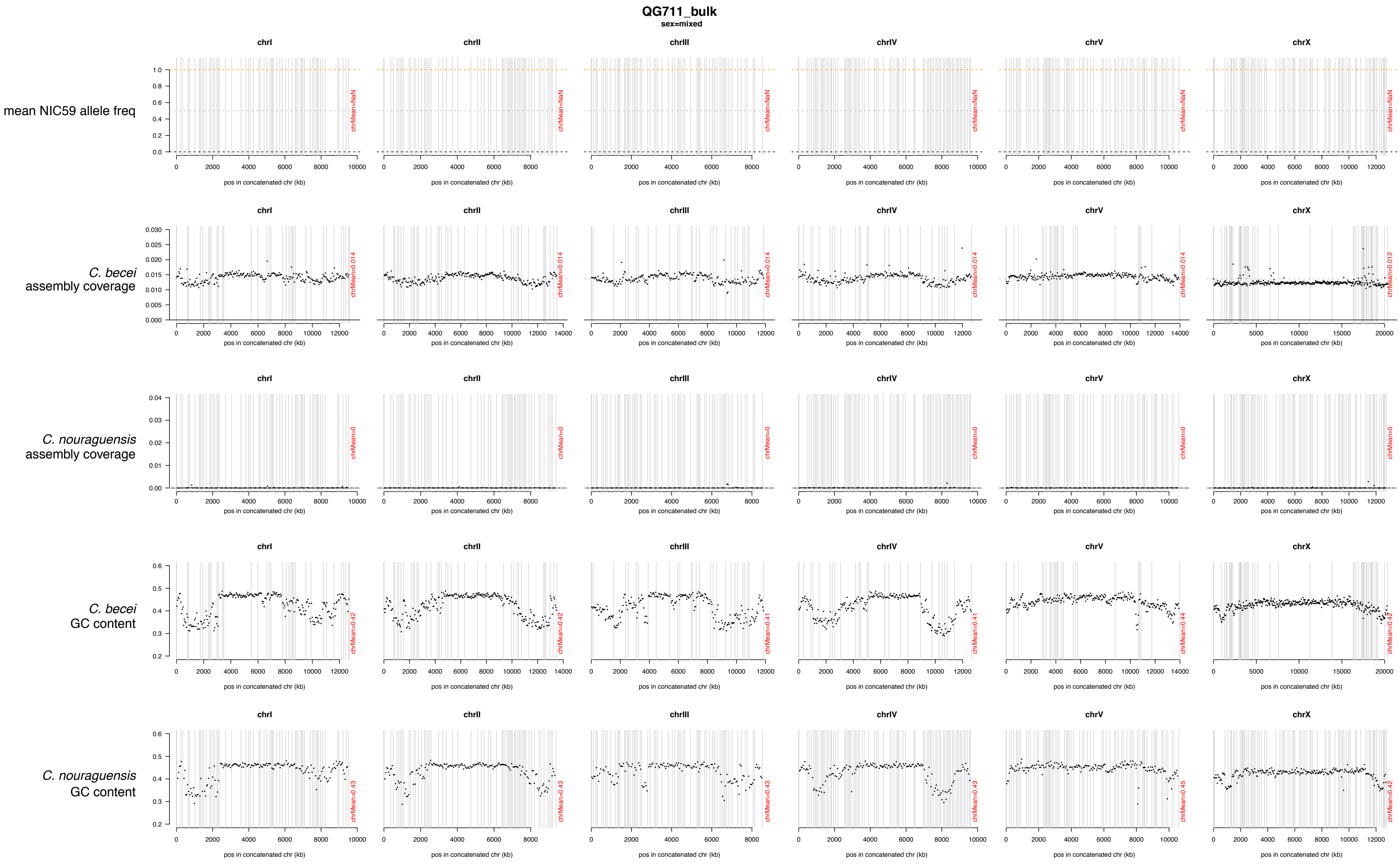
QG711_bulk
sex=mixed
chrI
chrII
chrIII
chrIV
chrV
chrX
mean NIC59 allele freq
chrMean=NaN
C. becei assembly coverage
chrMean=0.014
chrMean=0.012
C. nouraguensis assembly coverage
chrMean=0
C. becei GC content
chrMean=0.42
chrMean=0.41
chrMean=0.44
C. nouraguensis GC content
chrMean=0.43
chrMean=0.45
pos in concatenated chr (kb)

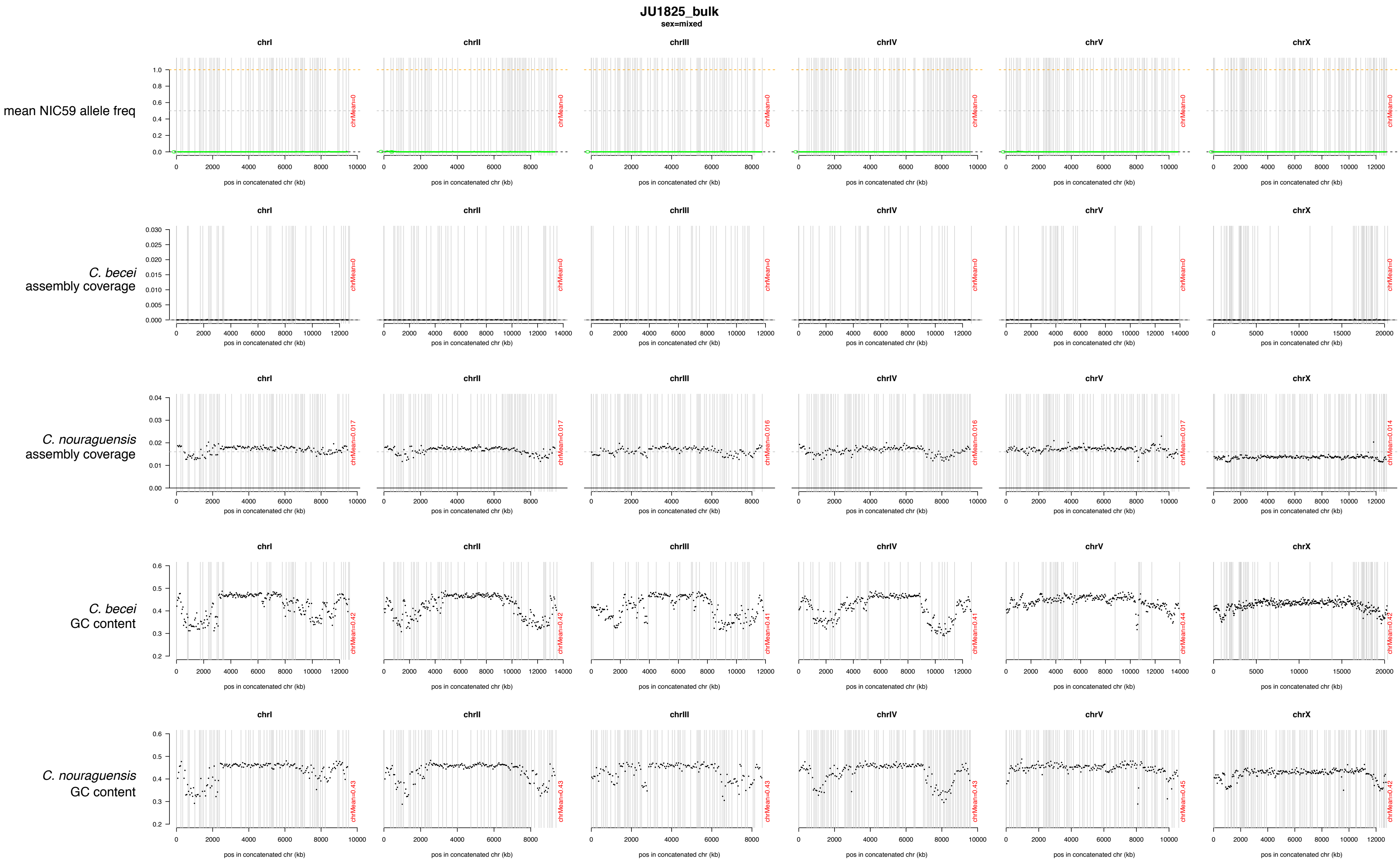
JU1825_bulk
sex=mixed
chrI
chrII
chrIII
chrIV
chrV
chrX
mean NIC59 allele freq
C. becei assembly coverage
C. nouraguensis assembly coverage
C. becei GC content
C. nouraguensis GC content
pos in concatenated chr (kb)
chrMean=0
chrMean=0.017
chrMean=0.016
chrMean=0.014
chrMean=0.42
chrMean=0.41
chrMean=0.44
chrMean=0.43
chrMean=0.45

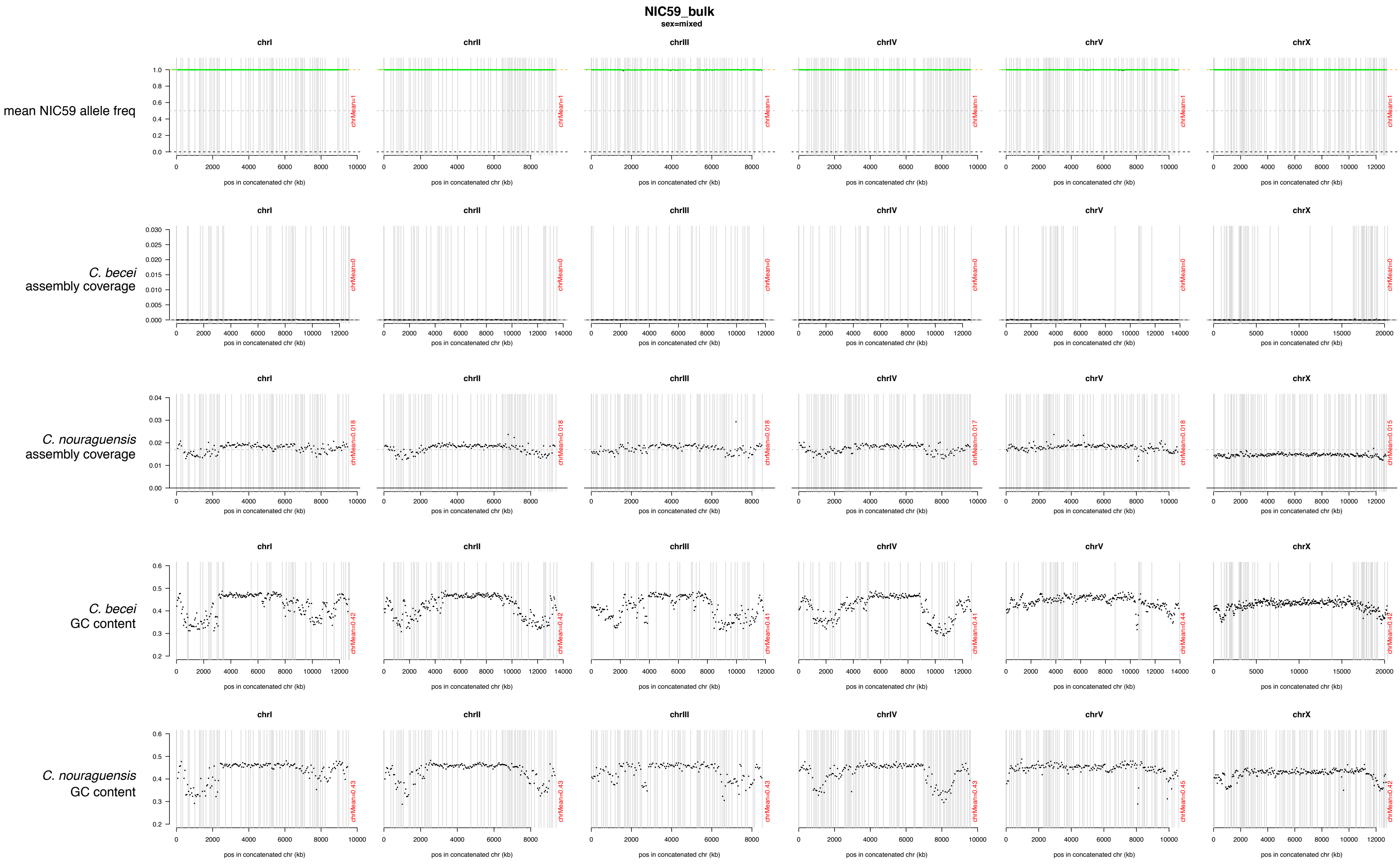
NIC59_bulk
sex=mixed
mean NIC59 allele freq
C. becei assembly coverage
C. nouraguensis assembly coverage
C. becei GC content
C. nouraguensis GC content
chrI
chrII
chrIII
chrIV
chrV
chrX
pos in concatenated chr (kb)

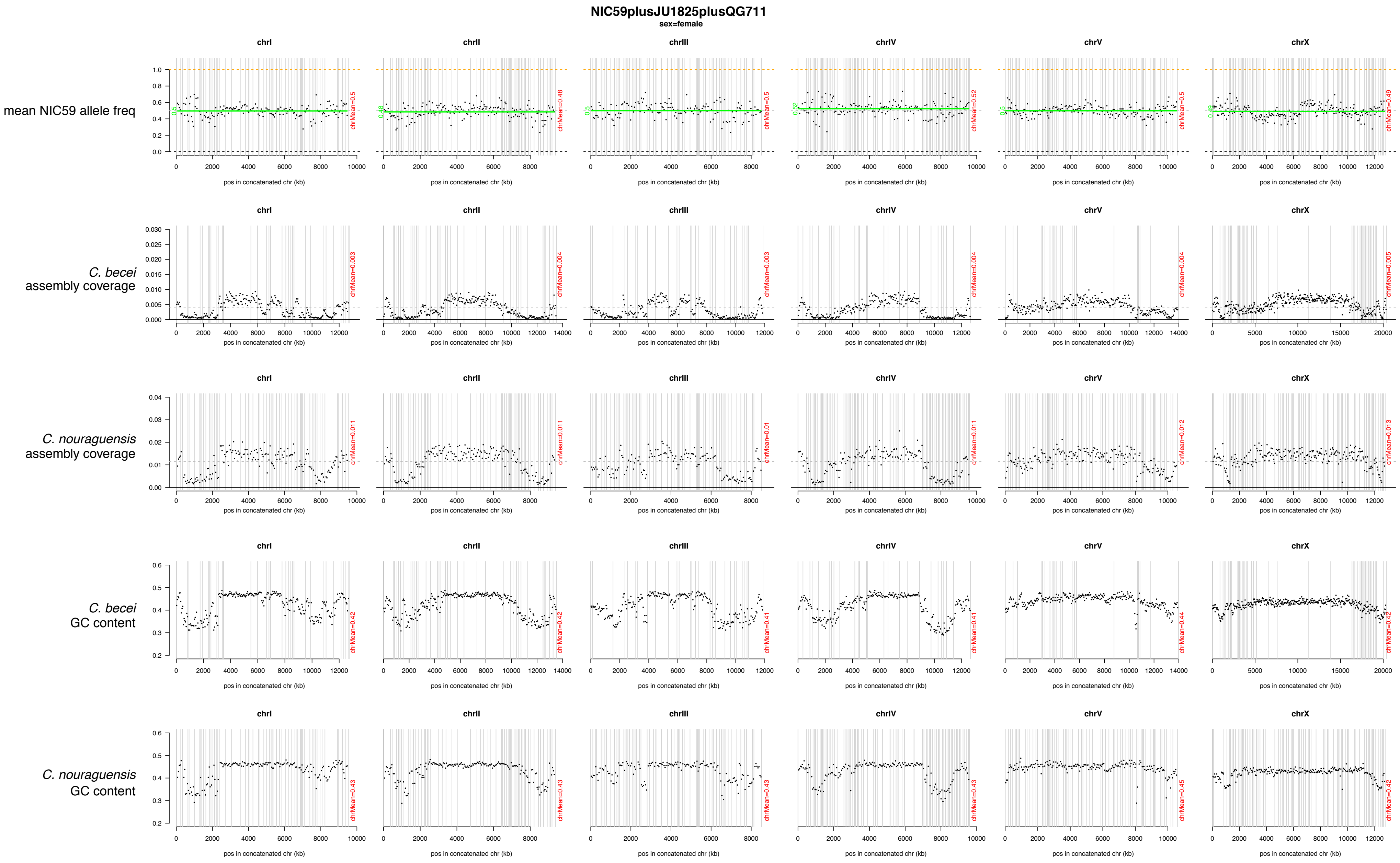

NIC59plusJU1825plusQG711
sex=female
mean NIC59 allele freq
C. becei assembly coverage
C. nouraguensis assembly coverage
C. becei GC content
C. nouraguensis GC content
chrI
chrII
chrIII
chrIV
chrV
chrX
pos in concatenated chr (kb)

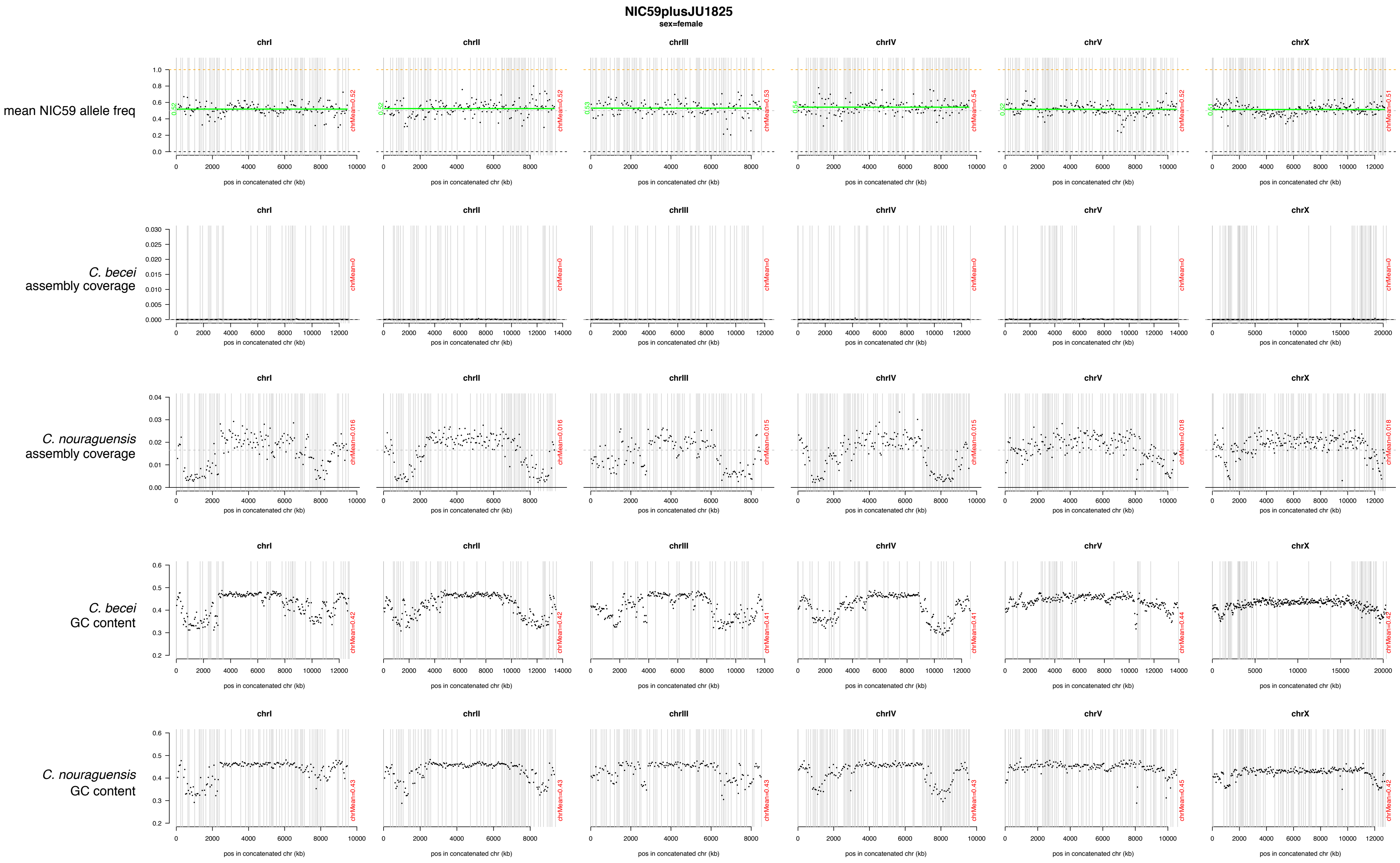

NIC59plusJU1825
sex=female
chrI
chrII
chrIII
chrIV
chrV
chrX
mean NIC59 allele freq
C. becei assembly coverage
C. nouraguensis assembly coverage
C. becei GC content
C. nouraguensis GC content
pos in concatenated chr (kb)
chrMean=0.52
chrMean=0.52
chrMean=0.53
chrMean=0.54
chrMean=0.52
chrMean=0.51
chrMean=0
chrMean=0.016
chrMean=0.016
chrMean=0.015
chrMean=0.015
chrMean=0.018
chrMean=0.018
chrMean=0.42
chrMean=0.42
chrMean=0.41
chrMean=0.41
chrMean=0.44
chrMean=0.42
chrMean=0.43
chrMean=0.43
chrMean=0.43
chrMean=0.43
chrMean=0.45
chrMean=0.42

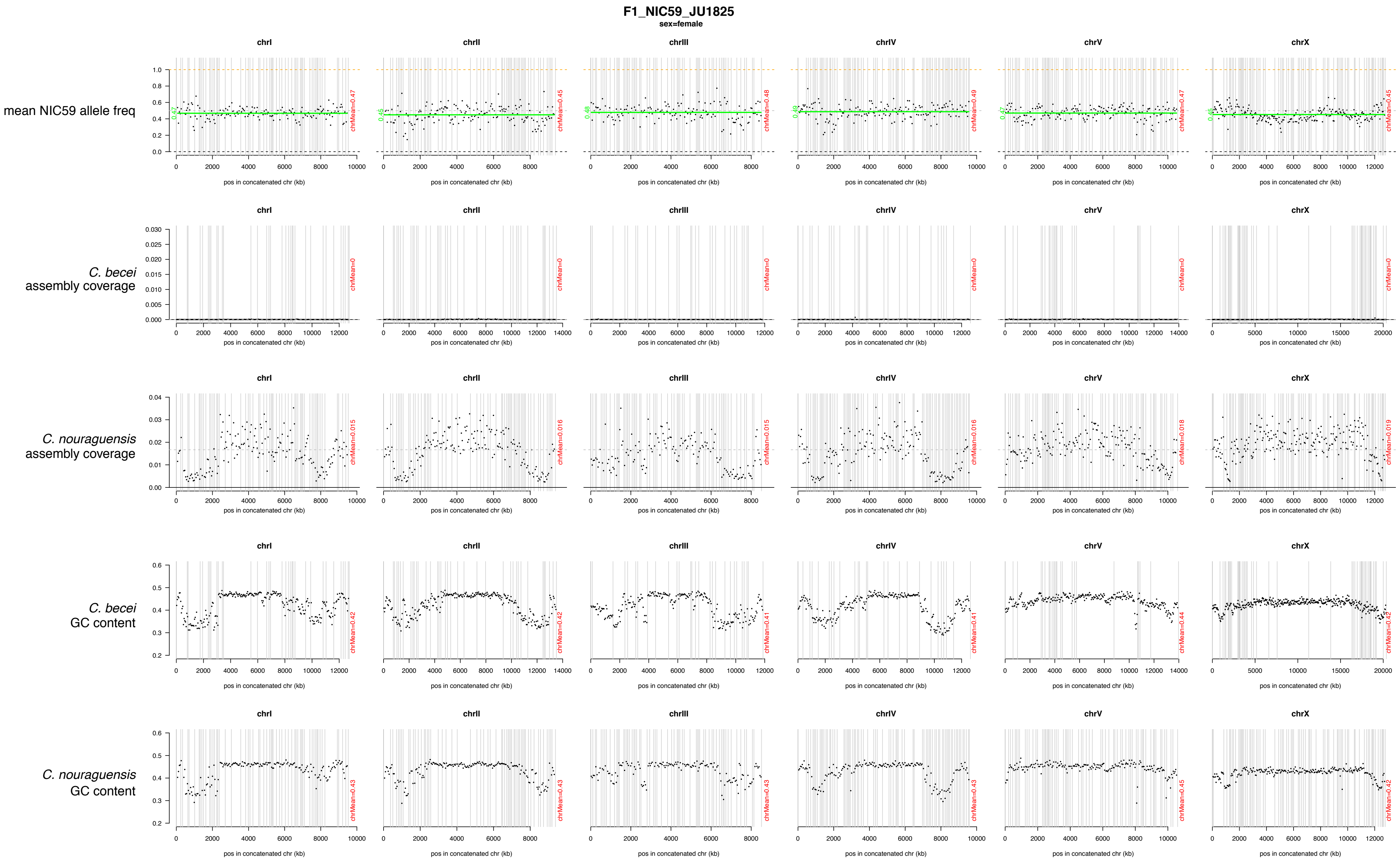

F1_NIC59_JU1825
sex=female
chrI
chrII
chrIII
chrIV
chrV
chrX
mean NIC59 allele freq
C. becei assembly coverage
C. nouraguensis assembly coverage
C. becei GC content
C. nouraguensis GC content
pos in concatenated chr (kb)

### Figure S8

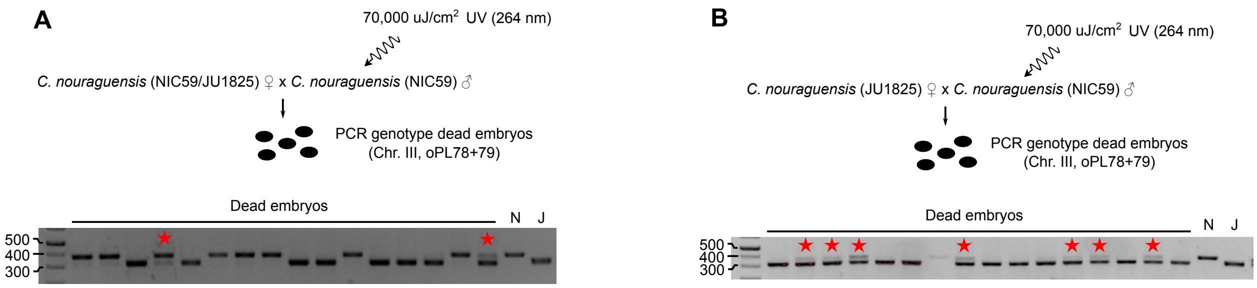


**Figure S8. Fertilization with UV-irradiated sperm does not cause frequent diploid maternal inheritance in intraspecies crosses. (A)** Fertilization with UV-irradiated sperm does not significantly increase diploid maternal inheritance in *C. nouraguensis* intraspecies crosses. Heterozygous *C. nouraguensis* (NIC59/JU1825) females were mated to UV-irradiated *C. nouraguensis* (NIC59) males. Dead embryos produced by this cross were PCR genotyped at a single autosomal locus (oPL78+79). The gel shows the genotypes of a fraction of the dead embryos assayed. Most embryos had either a JU1825 (11/31) or NIC59 genotype (17/31), consistent with haploid maternal inheritance and destruction of the paternal NIC59 genome. Although 3/31 embryos had a heterozygous genotype (red stars), we found that UV-irradiation can fail to completely destroy the paternal genome (see Figure S8B), suggesting that these three heterozygotes may result from haploid maternal inheritance of a JU1825 allele and haploid paternal inheritance of a NIC59 allele due to incomplete destruction of paternal DNA. **(B)** The paternal genome is not always eliminated by UV irradiation. *C. nouraguensis* (JU1825) females were mated to UV-irradiated *C. nouraguensis* (NIC59) males. Dead embryos produced by this cross were PCR genotyped at a single autosomal locus (oPL78+79). The gel shows the genotypes of a fraction of the dead embryos assayed. Red stars indicate cases in which the UV-irradiated paternal NIC59 DNA can be detected.

| | | | | | | After filtering, number of reads matching | | | | | | After filtering, percent of all assigned reads matching | | | After filtering, percent of all worm nuclear genome reads matching | | Approximate coverage | |
|---|---|---|---|---|---|---|---|---|---|---|---|---|---|---|---|---|---|---|
| **Sample name** | Cross F1 derived from | F1 sex | F1 crossed to | F1 fertility | Total # reads | *E. coli* | Firmicutes | *C. nouraguensis* nuclear genome | *C. becei* nuclear genome | *C. nouraguensis* mitochondrial genome | *C. becei* mitochondrial genome | *E. coli* | *C. nouraguensis* nuclear genome | C. becei nuclear genome | *C. nouraguensis* | *C. becei* | *C. nouraguensis* nuclear genome | *C. becei* nuclear genome |
| **F1_1** | (N); N/J F1 female x QG711 male Plate 1 | female | NIC59 male | fertile | 18,405,750 | 11,551,300 | 19 | 3,529,806 | 10,854 | 9,412 | 103 | 76.5 | 23.4 | 0.1 | 99.7 | 0.3 | 2.4 | 0.0 |
| **F1_5** | (N); N/J F1 female x QG711 male Plate 2 | female | NIC59 male | fertile | 4,709,920 | 306,730 | 6 | 3,291,061 | 6,828 | 5,605 | 61 | 8.5 | 91.2 | 0.2 | 99.8 | 0.2 | 2.2 | 0.0 |
| **F1_8** | (N); N/J F1 female x QG711 male Plate 2 | female | NIC59 male | fertile | 6,219,960 | 2,499,294 | 4 | 2,528,892 | 5,174 | 4,160 | 48 | 49.6 | 50.2 | 0.1 | 99.8 | 0.2 | 1.7 | 0.0 |
| **F1_11** | (N); N/J F1 female x QG711 male Plate 3 | female | NIC59 male | fertile | 5,556,004 | 1,228,558 | 3 | 3,169,866 | 6,505 | 1,879 | 20 | 27.9 | 71.9 | 0.1 | 99.8 | 0.2 | 2.2 | 0.0 |
| **F1_25** | (J); N/J F1 female x QG711 male Plate 3 | female | JU1825 male | fertile | 6,489,980 | 5,089,384 | 3 | 369,912 | 8,714 | 499 | 8 | 93.1 | 6.8 | 0.2 | 97.7 | 2.3 | 0.3 | 0.0 |
| **F1_29** | (J); N/J F1 female x QG711 male Plate 4 | female | JU1825 male | fertile | 7,097,772 | 667,344 | 13 | 4,884,750 | 9,066 | 1,495 | 16 | 12.0 | 87.8 | 0.2 | 99.8 | 0.2 | 3.3 | 0.0 |
| **F1_39** | (J); N/J F1 female x QG711 male Plate 1 | female | JU1825 male | fertile | 4,858,964 | 688,291 | 5 | 3,186,443 | 7,190 | 3,346 | 14 | 17.7 | 82.0 | 0.2 | 99.8 | 0.2 | 2.2 | 0.0 |
| **F1_41** | (J); N/J F1 female x QG711 male Plate 2 | female | JU1825 male | fertile | 18,011,610 | 1,112,932 | 20 | 12,756,921 | 21,799 | 2,298 | 19 | 8.0 | 91.8 | 0.2 | 99.8 | 0.2 | 8.7 | 0.0 |
| **F1_4** | (N); N/J F1 female x QG711 male Plate 1 | male | NIC59 female | fertile | 13,552,874 | 151,066 | 6 | 10,195,147 | 22,476 | 5,451 | 51 | 1.5 | 98.3 | 0.2 | 99.8 | 0.2 | 7.0 | 0.0 |
| **F1_46** | (N); N/J F1 female x QG711 male Plate 1 | male | NIC59 female | fertile | 20,496,550 | 4,494 | 15 | 15,463,159 | 35,591 | 7,759 | 83 | 0.0 | 99.7 | 0.2 | 99.8 | 0.2 | 10.6 | 0.0 |
| **F1_48** | (N); N/J F1 female x QG711 male Plate 4 | male | NIC59 female | fertile | 5,484,366 | 13,691 | 3 | 4,151,082 | 9,759 | 661 | 6 | 0.3 | 99.4 | 0.2 | 99.8 | 0.2 | 2.8 | 0.0 |
| **F1_6** | (N); N/J F1 female x QG711 male Plate 2 | female | NIC59 male | sterile | 4,477,900 | 930,700 | 7 | 1,932,322 | 705,081 | 1,202 | 15 | 26.1 | 54.1 | 19.8 | 73.3 | 26.7 | 1.3 | 0.4 |
| **F1_12** | (N); N/J F1 female x QG711 male Plate 3 | female | NIC59 male | sterile | 7,884,868 | 753,139 | 7 | 3,978,335 | 1,483,331 | 2,085 | 19 | 12.1 | 64.0 | 23.9 | 72.8 | 27.2 | 2.7 | 0.8 |
| **F1_18** | (J); N/J F1 female x QG711 male Plate 1 | female | JU1825 male | sterile | 3,468,706 | 1,496,149 | 3 | 1,373,037 | 7,652 | 96 | 0 | 52.0 | 47.7 | 0.3 | 99.5 | 0.6 | 0.9 | 0.0 |
| **F1_20** | (J); N/J F1 female x QG711 male Plate 2 | female | JU1825 male | sterile | 10,834,856 | 1,163,152 | 16 | 5,203,891 | 1,822,729 | 4,026 | 23 | 14.2 | 63.5 | 22.2 | 74.1 | 25.9 | 3.6 | 1.0 |
| **F1_26** | (J); N/J F1 female x QG711 male Plate 3 | female | JU1825 male | sterile | 3,594,908 | 891,072 | 3 | 1,923,182 | 92,400 | 690 | 4 | 30.6 | 66.1 | 3.2 | 95.4 | 4.6 | 1.3 | 0.0 |
| **F1_10** | (N); N/J F1 female x QG711 male Plate 2 | male | NIC59 female | sterile | 13,848,208 | 400,238 | 7 | 8,508,436 | 1,589,956 | 1,200 | 9 | 3.8 | 81.0 | 15.1 | 84.3 | 15.7 | 5.8 | 0.9 |
| **F1_16** | (N); N/J F1 female x QG711 male Plate 3 | male | NIC59 female | sterile | 15,639,042 | 288,535 | 2 | 10,408,444 | 1,046,790 | 16,142 | 172 | 2.5 | 88.5 | 8.9 | 90.9 | 9.1 | 7.1 | 0.6 |
| **F1_17** | (N); N/J F1 female x QG711 male Plate 3 | male | NIC59 female | sterile | 6,247,186 | 79,293 | 15 | 3,443,948 | 1,272,137 | 322 | 9 | 1.7 | 71.8 | 26.5 | 73.0 | 27.0 | 2.4 | 0.7 |
| **F1_21** | (J); N/J F1 female x QG711 male Plate 2 | male | JU1825 female | sterile | 9,834,614 | 137,887 | 6 | 5,701,336 | 1,741,990 | 480 | 5 | 1.8 | 75.2 | 23.0 | 76.6 | 23.4 | 3.9 | 0.9 |
| **F1_23** | (J); N/J F1 female x QG711 male Plate 2 | male | JU1825 female | sterile | 12,059,396 | 1,570,440 | 6 | 7,254,493 | 382,635 | 11,575 | 96 | 17.0 | 78.7 | 4.2 | 95.0 | 5.0 | 5.0 | 0.2 |
| **F1_NIC59_JU1825** | JU1825 female x NIC59 male | female | | | 8,237,250 | 75,877 | 14 | 6,344,331 | 12,349 | 3,180 | 38 | 1.2 | 98.6 | 0.2 | 99.8 | 0.2 | 4.3 | 0.0 |
| **NIC59plusJU1825** | | female | | | 7,045,822 | 319,354 | 15 | 5,091,335 | 10,238 | 2,668 | 21 | 5.9 | 93.9 | 0.2 | 99.8 | 0.2 | 3.5 | 0.0 |
| **NIC59plusJU1825plusQG711** | | female | | | 11,160,274 | 275,223 | 7 | 5,895,338 | 2,620,526 | 1,864 | 620 | 3.1 | 67.0 | 29.8 | 69.2 | 30.8 | 4.0 | 1.4 |
| **NIC59_bulk** | | mixed | | | 32,250,274 | 257,304 | 31 | 22,649,102 | 52,717 | 50,912 | 437 | 1.1 | 98.4 | 0.2 | 99.8 | 0.2 | 15.5 | 0.0 |
| **JU1825_bulk** | | mixed | | | 36,926,106 | 357,701 | 394 | 27,761,970 | 44,405 | 58,749 | 279 | 1.3 | 98.4 | 0.2 | 99.8 | 0.2 | 19.0 | 0.0 |
| **QG711_bulk** | | mixed | | | 28,828,734 | 235,533 | 37 | 120,477 | 21,138,043 | 17 | 41,249 | 1.1 | 0.6 | 98.2 | 0.6 | 99.4 | 0.1 | 11.4 |

**Table S1. Characteristics of the samples submitted for whole-genome sequencing.** Whole-genome amplified F1 individuals derived from the interspecies cross are labeled as "F1_" followed by a barcode number. Sample "F1_NIC59_JU1825" is one heterozygous NIC59/JU1825 female that was whole-genome amplified and serves as a control for genome-wide heterozygosity. Sample "NIC59plusJU1825" comprises one NIC59 and one JU1825 female placed in the same tube and whole-genome amplified. This also serves as a control for genome-wide heterozygosity. Sample "NIC59plusJU1825plusQG711" comprises one NIC59, one JU1825 and one QG711 female placed into the same tube and whole-genome amplified. This serves as a triploid (diploid *C. nouraguensis* and haploid *C. becei*) control. The "NIC59_bulk", "JU1825_bulk" and "QG711_bulk" samples are genomic preps made from large populations of each strain containing mixed developmental stages and sexes: these libraries were prepared without whole-genome amplification. The reads from the "NIC59_bulk" and "JU1825_bulk" samples were used to identify fixed SNPs between the two strains. The second column shows the interspecies cross that produced each viable F1 animal. In each case, heterozygous N/J *C. nouraguensis* females were crossed to *C. becei* QG711 males. However, the heterozygous *C. nouraguensis* females could have either a NIC59 or JU1825 mitochondrial genotype (denoted in parentheses). A fraction of each sample's reads (Illumina 50-bp paired end) derive from *E.coli* (their food). The *E. coli* fraction is quite high in some samples, likely due to inefficient cleaning of single worms (see Materials and Methods). Of the reads that map to *Caenorhabditis* nuclear genomes, the approximate coverage and percent of all worm reads that map to each *Caenorhabditis* nuclear genome are shown.
